# Intrinsic Gestational Timing Governs Human Cerebellar Development After Preterm Birth

**DOI:** 10.1101/2025.09.01.673244

**Authors:** Georgios Sanidas, Gabriele Simonti, Javid Ghaemmaghami, Kyla Woyshner, Antonis Polyviou, Robinson Vidva, Courtney Lowe, Abhya Vij, Daniel Bittel, Maria Triantafyllou, Nora Wolff, Rodolfo Cardenas, Helen Chen, Francisco Almeida, Sniya Sudhakar, Kshitij Mankad, Ioannis Koutroulis, Henrik Sahlin Pettersen, Genevieve Stein-O’Brien, Dimitrios N Sidiropoulos, Vittorio Gallo, Panagiotis Kratimenos

## Abstract

Intrinsic programs shape brain maturation, yet which are perturbed by prematurity, and the molecular pathways involved remain unknown. The human cerebellum serves as a paradigm of extrauterine development; its accelerated growth and circuit formation align with the third trimester, a period disrupted by preterm delivery. Through multimodal datasets encompassing in vivo neuroimaging and neurodevelopmental outcomes with postmortem spatial-transcriptomic and histopathological profiling, we show that prematurity diverts the cerebellar developmental trajectory. Cerebellar growth and functional outcomes scaled with gestational age, despite modifying perinatal exposures. Spatially resolved developmental programs underlying macroscopic trajectories were marked by incomplete granule cell maturation and impaired Purkinje cell structural refinement, indicating lineage-specific developmental asynchrony. Thus, prematurity constitutes biologically displaced maturation, in which infants of equivalent post-menstrual age occupy divergent developmental states.

## Introduction

Developmental timing, fundamental to biology, orchestrates genetic programs that produce complex tissues and organs (*1*). In the human brain, neuronal migration and circuit assembly proceed in a tightly coordinated sequence of maturation and growth acceleration as gestation advances (*2*). Notably, between 24 and 40 weeks of gestation, the cerebellum expands more rapidly than any other brain region, increasing about fivefold in volume (*3, 4*).

In preterm-born infants, this period of peak cerebellar growth is completed ex utero, providing a unique opportunity to study how early birth reshapes an actively unfolding developmental program (*4, 5*). While this disruption is known to alter cerebellar and neurodevelopmental trajectories (*6, 7, 8, 9*), its molecular and cellular basis remains undefined (*4, 10*).

Answering this question requires molecular characterization of human extrauterine development, a distinct state that cannot be fully recapitulated in model systems.

To bridge this gap, we assembled an extensive postmortem cerebellar series combined with an independent in vivo cohort spanning a broad range of gestational ages. Using spatial transcriptomics and deep-learning–based histopathology, combined with longitudinal imaging and subsequent neurodevelopmental assessment, we resolved developmental trajectories and showed that birth timing constrains lineage-specific maturation processes. These findings position gestational age at birth as a key temporal axis of postnatal cerebellar maturation, with enduring structural and neurodevelopmental correlates.

## Results

### Gestational age at birth scales cerebellar growth and neurodevelopment across neonatal exposures

We assembled a gestational age-stratified cohort of 176 infants lacking major destructive brain injury, with detailed perinatal exposure profiling (fig. S1; table S1; data S1). Infants with major destructive brain injury were excluded to isolate developmental variation from acquired injury. Term-equivalent brain MRI (*n* = 120), serial cranial ultrasonography (*n* = 146), and neurodevelopmental follow-up (*n* = 145) were available, with 102 infants assessed by both imaging modalities and 84 across all three measures (fig. S1; fig. S2).

At term-equivalent age (37 to 42 weeks post-menstrual age), cerebellar volume was 230 mm³ greater for each additional gestational week at birth (Fig. 1A), in proportion to overall brain size (fig. S3). This association was robust to variation in the perinatal exposures, although cumulative bronchopulmonary dysplasia (BPD) burden partially attenuated the effect (fig. S4A; table S2).

**Fig. 1.**
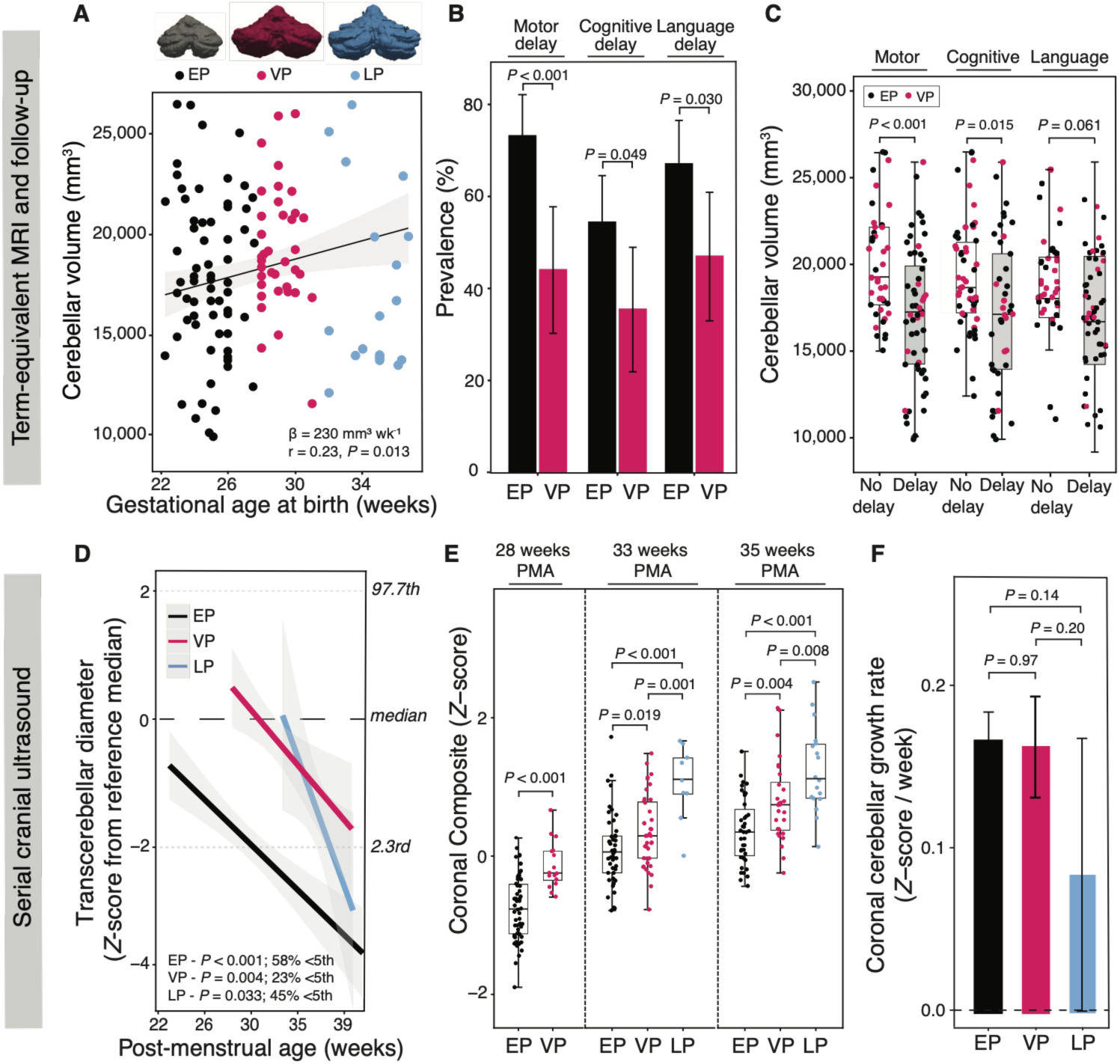
Gestational age at birth organizes cerebellar growth and neurodevelopment after preterm birth. **(A)** Cerebellar volume at term-equivalent age (TEA) plotted against gestational age at birth in extremely preterm (EP, <28 weeks; black), very preterm (VP, 28 to <32 weeks; magenta), and late preterm (LP, 32 to <37 weeks; light blue) infants (*n* = 120; EP = 65, VP = 37, LP = 18). Representative three-dimensional cerebellar reconstructions from each gestational-age group are shown above the plot. Cerebellar volume increased with gestational age at birth (multivariable OLS regression adjusted for post-menstrual age at MRI; β = 230 mm³ week⁻¹; partial *r* = 0.23; *P* = 0.013); shaded band, 95% CI. **(B)** Prevalence of motor, cognitive, and language delay in EP (*n* = 94) and VP (*n* = 51) infants, assessed by Bayley-III or ASQ-3. Bars, proportions; error bars, 95% CI; *P* values, two-sided Fisher’s exact tests. **(C)** Cerebellar volume at TEA in infants with or without motor (left), cognitive (center), or language (right) delay among EP and VP infants with paired MRI and neurodevelopmental follow-up (*n* = 98); dots colored by gestational-age group. Box plots, median and IQR; *P* values, two-sided Wilcoxon rank-sum tests. **(D)** Longitudinal ultrasonographic measurements of transcerebellar diameter, expressed as post-menstrual age–specific *Z*-scores relative to fetal reference standards. Lines, group-specific fits; shaded bands, 95% CI; dashed lines, reference median and 2.3rd/97.7th centiles. *Z*-scores declined progressively with advancing PMA in each group (linear mixed-effects model); percentages indicate the proportion of observations below the 5th reference centile. **(E)** Ultrasonography-derived coronal composite cerebellar size (*Z*-score) at 28, 33, and 35 weeks PMA; the 28-week comparison includes EP and VP infants only. Box plots, median and IQR; *P* values, pairwise two-sided Wilcoxon rank-sum tests. **(F)** Longitudinal coronal cerebellar growth rate derived from serial ultrasonography (*Z*-score week⁻¹) across gestational-age groups. Bars, mean; error bars, 95% CI; *P* values, pairwise slope contrasts from linear mixed-effects models with Tukey adjustment. (D to F), *n* = 146 (EP = 80, VP = 45, LP = 21).

In the same cohort, this structural gradient extended to later neurodevelopment, with lower gestational age at birth associated with higher rates of motor, cognitive, and language delay at 9 to 36 months corrected age (Fig. 1B) and smaller absolute cerebellar volume selectively associated with motor and cognitive delay (Fig. 1C). Across variation in perinatal exposures, these relationships were largely preserved, with gestational age continuing to be associated with motor and language delay and cerebellar volume independently associated with motor and cognitive delay. Neither relationship differed by sex (fig. S4B; table S3; table S4), and twin status did not account for differences in cerebellar volume or neurodevelopmental delay (fig. S5).

Serial cranial ultrasonography measurements were concordant with paired MRI assessments (fig. S6A-B). Transcerebellar diameter progressively diverged from fetal normative growth references across gestational-age groups, with no significant difference in the degree of divergence among groups (*11*) (Fig. 1D). Gestational age–dependent differences in cerebellar size were evident by the earliest assessed time point, 28 weeks post-menstrual age, and persisted to term-equivalent age in both the coronal (Fig. 1E) and midsagittal planes (fig. S6C). Postnatal growth rates were comparable across gestational age groups (Fig. 1F; fig. S6D; fig. S7A-B), indicating that early differences were maintained rather than progressively amplified. The same size patterns remained evident after adjustment for individual clinical exposures (fig. S4A; table S5). Sex did not alter these trajectories, although coronal dimensions were consistently greater in males (table S3).

Together, these findings identify gestational age at birth as a major correlate of early cerebellar growth and subsequent motor and cognitive outcomes, with individual features of the neonatal intensive care course modifying rather than accounting for this relationship.

### Gestational age at birth organizes lineage-specific cerebellar molecular programs

The gestational age–dependent scaling of cerebellar growth and neurodevelopment implicated developmental programs whose progression is altered by the timing of birth. To define these programs in an independent human cohort, we analyzed cerebellar tissue from 11 subjects within an 83-subject postmortem series, spanning the gestational-age spectrum and assessed at comparable post-menstrual ages (fig. S1; fig. S8; table S6; table S7; data S2). Spatial transcriptomic profiling of the developing cerebellar cortex, coupled with CoGAPS (*12*), resolved gene expression into 28 coordinated, unsupervised transcriptional patterns (H-1 to H-28; Fig. 2A; data S3). Three patterns showed both gestational age–dependent variation and coherent biological annotation and were designated Cerebellar Maturation–Associated Patterns (CMAPs) (Fig. 2B–E; see Materials and Methods). These localized to the Purkinje cell layer (CMAP_PCL [H-27]) and internal granule layer (CMAP_IGL_IMM [H-16; IMM, immature] and CMAP_IGL [H-17]; fig. S9A–E; data S4), together delineating a coordinated maturation trajectory across Purkinje and granule cell lineages.

**Fig. 2.**
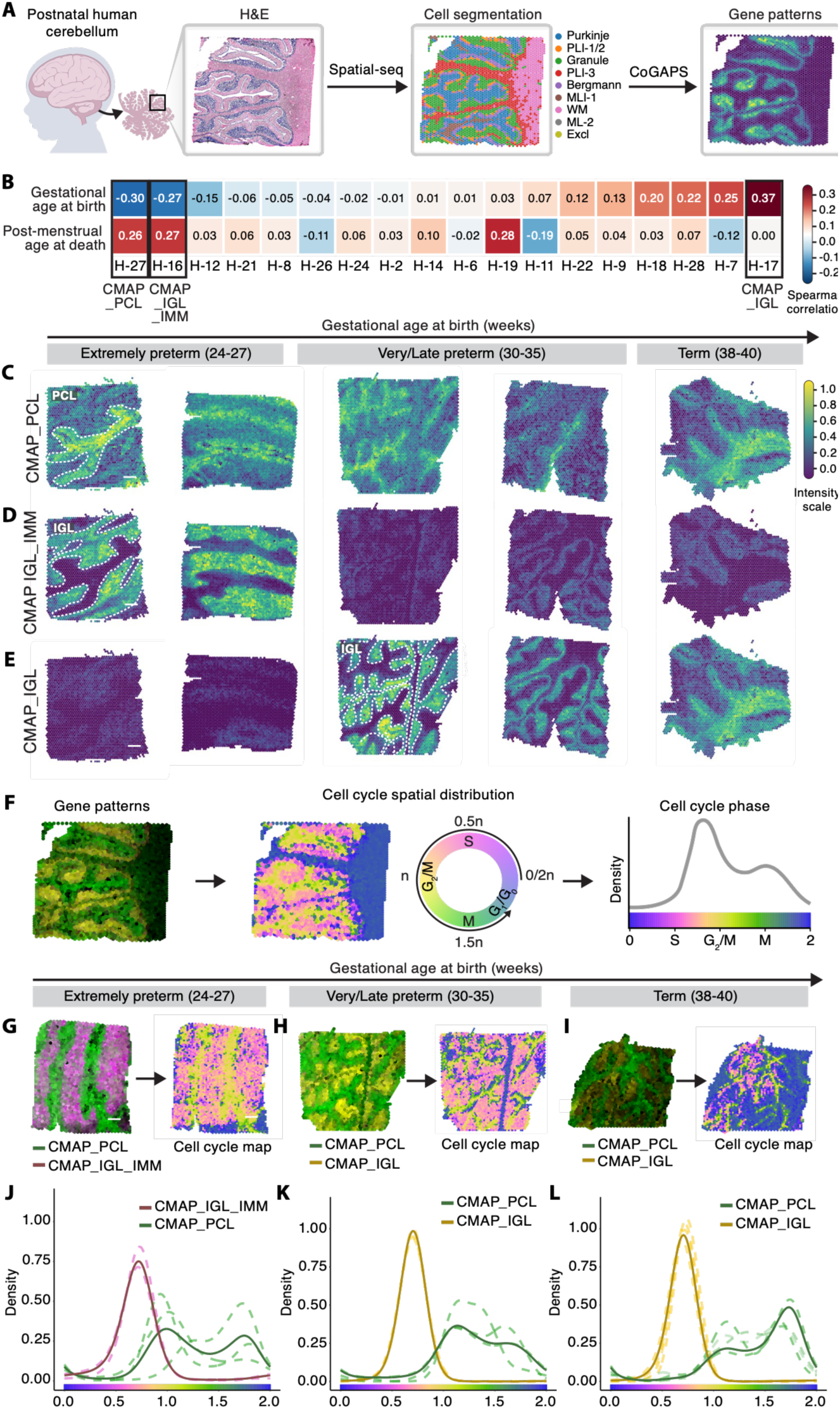
Gestational age at birth organizes lineage-specific cerebellar maturation programs and phase-associated states. **(A)** Spatial transcriptomic workflow in postnatal human cerebellum. Cerebellar sections from 11 subjects (7 preterm and 4 term; 22 sections, two per subject; tables S6 and S7) were annotated by laminar and cellular compartments and decomposed by Bayesian non-negative matrix factorization (CoGAPS) into transcriptional patterns, including three cerebellar maturation-associated patterns (CMAPs). H&E, hematoxylin and eosin. **(B)** Spearman correlations between subject-level CoGAPS pattern activity and gestational age (GA) at birth or post-menstrual age (PMA) at death. H-27 (CMAP_PCL) and H-16 (CMAP_IGL_IMM) were negatively associated with GA, whereas H-17 (CMAP_IGL) was positively associated with GA. **(C to E)** Representative spatial activity maps of CMAP_PCL (C), CMAP_IGL_IMM (D), and CMAP_IGL (E) across increasing GA. White dashed contours delineate the corresponding Purkinje cell layer (PCL) or internal granule layer (IGL). Scale bar, 1 mm. **(F)** Projection of CMAP-associated spots onto the continuous cell-cycle–based transcriptional axis inferred by Tricycle, spanning G0/G1, S, G2/M, and M phase-associated positions. **(G to I)** CMAP activity (left) and inferred phase-associated transcriptional position (right) in extremely preterm (G; GA 24–27 weeks), very/late preterm (H; GA 30–35 weeks), and term (I; GA 38–40 weeks) cerebella. Scale bar, 1 mm. **(J to L)** Kernel density distributions of CMAP-associated spots along the Tricycle continuum in extremely preterm (J, *n* = 3), very/late preterm (K, *n* = 4), and term (L, *n* = 4) subjects. Dashed lines indicate individual subjects and solid lines pooled distributions; spots were restricted to those above the 95th percentile of pattern weight. CMAP_IGL_IMM occupied earlier phase-associated positions in extremely preterm cerebella, whereas CMAP_IGL predominated with advancing gestation at a narrower S-to-G2 distribution. CMAP_PCL co-regionalized with CMAP_IGL_IMM in the extremely preterm cerebella and shifted toward later phase-associated positions with advancing GA. Color bars indicate inferred Tricycle position.

CMAP_PCL, defined by canonical Purkinje cell layer genes (*CALB1, ITPR1, PCP2*, and *ALDOC*), was enriched in infants born at lower gestational ages and increased with longer ex utero survival (Fig. 2B-C; fig. S9C). This pattern was enriched for genes associated with ribosomal, biosynthetic, and metabolic pathways (fig. S10), consistent with the high anabolic and biosynthetic demands of the Purkinje layer during early development.

Within the internal granule layer, two transcriptional programs sequentially occupied the same laminar compartment, consistent with temporal program replacement rather than spatial reorganization (Fig. 2D-E; fig. S9B). At lower gestational ages, CMAP_IGL_IMM predominated and was defined by regulators of granule cell differentiation (*NEUROD1*, *ZIC2*, *CBLN1*, and *RELN)* and was functionally weighted toward neurogenesis and synaptic organization pathways (Fig. 2B, D; fig. S9D; fig. S10).

Across subjects, increasing gestational age at birth was associated with a transition from the nascent CMAP_IGL_IMM to the more mature CMAP_IGL state, marked by synaptic and vesicle-associated genes (*SNAP25*, *CBLN3*, *VSNL*1, and *CHGB*; Fig. 2B, E; fig. S9E; fig. S10) and coordinated divergence in pathways related to synaptic activity and neurotransmitter release. Consistent with this shift toward a mature granule-cell synaptic program, VGLUT1⁺ parallel-fiber labeling was higher across samples with increasing gestational age at birth. In parallel, VGLUT2⁺ climbing-fiber labeling was lower across samples with increasing gestational age at birth (fig. S11A–D; data S5). Together, these patterns are consistent with the canonical developmental VGLUT2-to-VGLUT1 transition (*13, 14*) and progressive maturation of excitatory cerebellar circuitry.

Next, we applied partitioned heritability analysis to map adult-derived genome wide association studies to CMAP gene sets based on genomic proximity. The functional relevance of these programs is underscored by the enrichment of CMAP_PCL genes for adult loci associated with cerebellar volume, while CMAP_IGL and CMAP_IGL_IMM were enriched for loci associated with cognitive reaction time (fig. S12; table S8), supporting the broader biological relevance of these gestationally ordered programs, beyond the neonatal period. The associations of CMAP_IGL and CMAP_PCL with gestational age at birth persisted across strata defined by sex and perinatal exposures, whereas associations with post-menstrual age were less consistent, particularly for CMAP_PCL (table S9).

Together, these findings delineate lineage-specific, gestationally coupled programs that track human cerebellar ontogeny, encode biosynthetic and synaptic maturation states, remain robust across clinical heterogeneity, and align with genetic variation associated with cerebellar volume and cognitive function in the general population.

### Preterm birth is associated with altered phase-associated maturation states across cerebellar lineages

Having identified lineage-specific gene expression patterns (CMAPs) graded by gestational age at birth, we next asked whether these programs reflect constrained progression through developmental transcriptional states. To address this, we applied Tricycle to map CMAPs onto a continuous, cell-cycle-based transcriptional topology, enabling lineage-resolved positioning across phase-associated states (Fig. 2F).

CMAP_IGL_IMM, characteristic of extremely preterm cerebella, was broadly distributed from G1/S through S-G2 positions corresponding to early differentiation and immature transcriptional states. In subjects born at later gestational ages, the IGL program transitioned from CMAP_IGL_IMM to a predominant CMAP_IGL signature, which was more narrowly restricted to S-G2 phases positions (Fig. 2G–L).

These phase-associated states were validated in situ at subcellular resolution using RNAscope in an independent validation cohort (fig. S1). To distinguish the proliferative profile of granule cell progenitors from cell cycle–associated transcriptional states in immature granule cells in the IGL, we first examined the external granule layer (EGL). In the EGL, canonical proliferative markers, including MCM2, were higher across late preterm and term samples than in extremely preterm samples, consistent with progenitor maturation at later gestational ages (fig. S13A-C).

We then examined the IGL to determine whether immature granule cells followed a similar trajectory. In contrast to the EGL, *MCM2* expression was higher across extremely preterm samples and lower across late preterm and term samples, paralleling the early-phase distribution of CMAP_IGL_IMM (fig. S13D-F). This gestational age–dependent pattern of MCM2 expression within the IGL is consistent with an intermediate cell-cycle-associated state between proliferative precursors and mature granule cells in extreme prematurity. In support of this transition, the late-phase G2/M regulators *CCNB1* and *TOP2A* showed greater expression with advancing gestational age at birth in the mature IGL, aligning with progression toward the later-phase transcriptional state marked by CMAP_IGL (fig. S13D-F).

Unlike IGL programs, CMAP_PCL showed a gestational age-dependent redistribution across the cell-cycle phase continuum. In extreme preterm cerebella, CMAP_PCL was centered near the G2/M-to-M-phase region and shifted toward later M-phase positions with increasing gestational age at birth, consistent with maturational progression in the Purkinje cell layer (fig. 2G–L). As chromatin-condensation is a defining feature of M phase, we quantified phosphorylated histone H3 (PHH3) as an independent marker of chromatin state (fig. S13G-H). PHH3 correlated inversely with gestational age at birth (fig. S13H-I): in extremely preterm cerebella, PHH3 appeared as condensed puncta within Purkinje cell nuclei (fig. S13H, left panel) whereas in late preterm and term subjects, nuclear PHH3 was reduced with relative cytoplasmic redistribution (fig. S13H, right panel). This pattern is consistent with persistent nuclear chromatin compaction in extremely preterm, post-mitotic Purkinje cells, linking sustained condensation to immature transcriptional states and delayed maturational progression.

These findings show that preterm birth is associated with gestational age-graded immature states, consistent with limited advancement along the maturation continuum.

### Gestational age at birth and extrauterine time differentially shape cerebellar laminar maturation in a large human neonatal cohort

To validate the molecular findings at the cellular level, we analyzed cerebella from 77 subjects within the 83-subject postmortem cohort, spanning the preterm-to-term gestational spectrum. Seventy-two subjects were independent of the spatial-transcriptomic discovery cohort and five overlapped, providing predominantly orthogonal validation (fig. S1; fig. S14; table S10; table S11; table S12).

Cerebellar laminar architecture was quantified from whole-slide images using deep-learning– based quantitative neuropathology (*15*), with associations between gestational age at birth and layer-level measures adjusted for infant sex and individual perinatal exposures (Fig. 3A; fig. S15; fig. S16A-D; table S13; table S14).

**Fig. 3.**
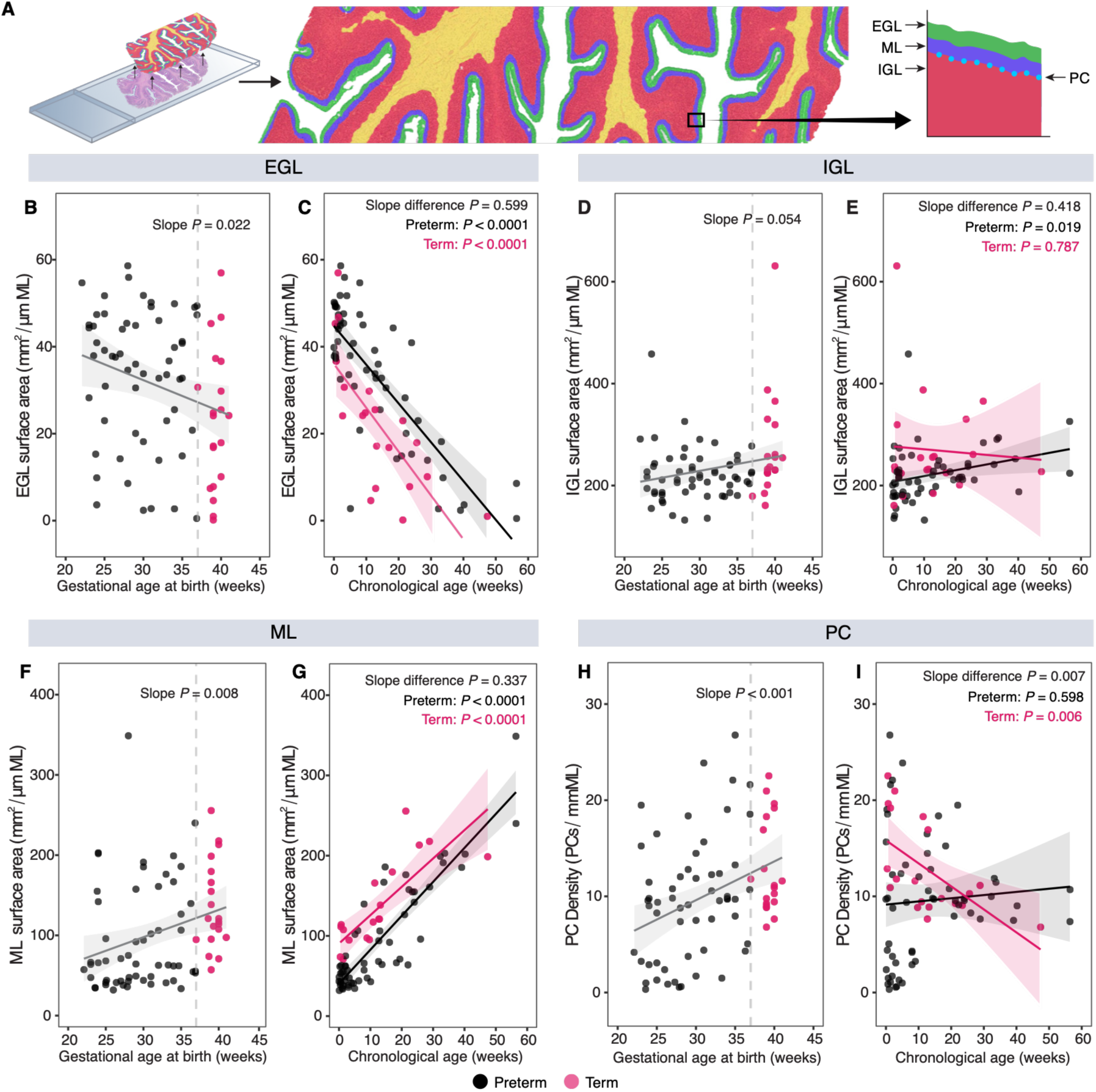
Cerebellar laminar architecture is structured by gestational age at birth. **(A)** Deep-learning quantitative neuropathology workflow. Whole-slide hematoxylin and eosin (H&E)– stained postmortem cerebellar sections were segmented to identify the external granule layer (EGL), internal granule layer (IGL), molecular layer (ML), and Purkinje cell (PC) layer. **(B to I)** Associations of cerebellar laminar measures with gestational age at birth (left) and chronological age (right). EGL surface area (B and C), IGL surface area (D and E), and ML surface area (F and G) were normalized to ML length (mm² per µm ML); PC density (H and I) was expressed as PCs per mm ML. Each dot represents one subject (*n* = 77; preterm, *n* = 57, black; term, *n* = 20, magenta). Lines, ordinary least-squares regression fits; shaded bands, 95% CI. Left panels report the *P* value for the overall association with gestational age at birth; dashed lines mark 37 weeks. Right panels show preterm– and term-specific trajectories and the *P* value for the difference between slopes: EGL (C), preterm β = −0.89 [−1.07 to −0.71], *P* < 0.0001, term β = −1.01 [−1.40 to −0.61], *P* < 0.0001, slope difference *P* = 0.599; IGL (E), preterm β = 1.14 [0.21 to 2.07], *P* = 0.019, term β = −0.55 [−4.52 to 3.41], *P* = 0.787, slope difference *P* = 0.418; ML (G), preterm β = 4.23 [3.65 to 4.81], *P* < 0.0001, term β = 3.53 [2.24 to 4.82], *P* < 0.0001, slope difference *P* = 0.337; PC (I), preterm β = 0.03 [−0.09 to 0.16], *P* = 0.598, term β = −0.24 [−0.39 to −0.09], *P* = 0.006, slope difference *P* = 0.007.

Infants born at lower gestational ages exhibited greater postnatal persistence of the EGL than term-born infants (Fig. 3B). Although EGL surface area declined with chronological age in both groups, the trajectories remained parallel but offset, indicating that postnatal EGL persistence was strongly associated with gestational age at birth (Fig. 3C), with additional modification by perinatal factors (fig. S16A; table S14). The expanded EGL in preterm infants, despite comparable cellular density, reflected a larger residual granule cell progenitor pool rather than a dilution effect, consistent with delayed EGL depletion and inward migration after preterm birth (fig. S17A-B).

In contrast, postnatal IGL surface area was greater with increasing gestational age at birth (Fig. 3D; fig. S16B) and followed similar trajectories in term and preterm infants across chronological age (Fig. 3E). This preservation of laminar growth, despite gestationally displaced molecular states (Fig. 2D–E), suggests that IGL maturation proceeds through transcriptional state transitions within a preserved laminar scaffold rather than through overt laminar reorganization.

The molecular layer, which contains the Purkinje cell dendritic arbor, was smaller in extremely preterm cerebella (Fig. 3F; fig. S16C). Molecular layer surface area increased with chronological age in both groups, with no significant difference in slope between preterm and term infants (Fig. 3G). Purkinje cell density followed a divergent trajectory: it scaled with gestational age at birth (Fig. 3H; fig. S16D), whereas with advancing chronological age, Purkinje cell density declined in term infants, consistent with structural refinement, but remained unchanged in preterm infants over the same interval (Fig. 3I). Thus, after preterm birth, dendritic expansion proceeded without the coordinated decrease in Purkinje cell density seen in term cerebella, indicating growth uncoupled from synaptic pruning and optimization.

Collectively, these data demonstrate that preterm birth is characterized by the prolonged persistence of GC progenitors (*16*) and a concomitant disruption of the physiological PC pruning (*17, 18*), fundamental to the maturation process inherent to term infants.

### Single-gene readouts resolve lineage-specific cerebellar maturation

By modeling transcript expression within the spatial-transcriptomic cohort (fig. S1) as a function of gestational age across developmentally matched preterm and term subjects after term-equivalent age, we sought to identify marker genes that track lineage maturation in Purkinje and granule cells (Fig. 4A-H). Lower gestational age at birth was associated with enrichment of early-lineage and stress-associated genes, including ribosomal and oxidative stress markers such as *RPS26*, *HSPA1A*, and *MT3* (fig. S18A–B), consistent with a reactive state in the most preterm cerebella. With increasing gestational age at birth, expression shifted toward genetic programs supporting neuronal differentiation, synaptic integration, and circuit maturation (Fig. 4A, E).

**Fig. 4.**
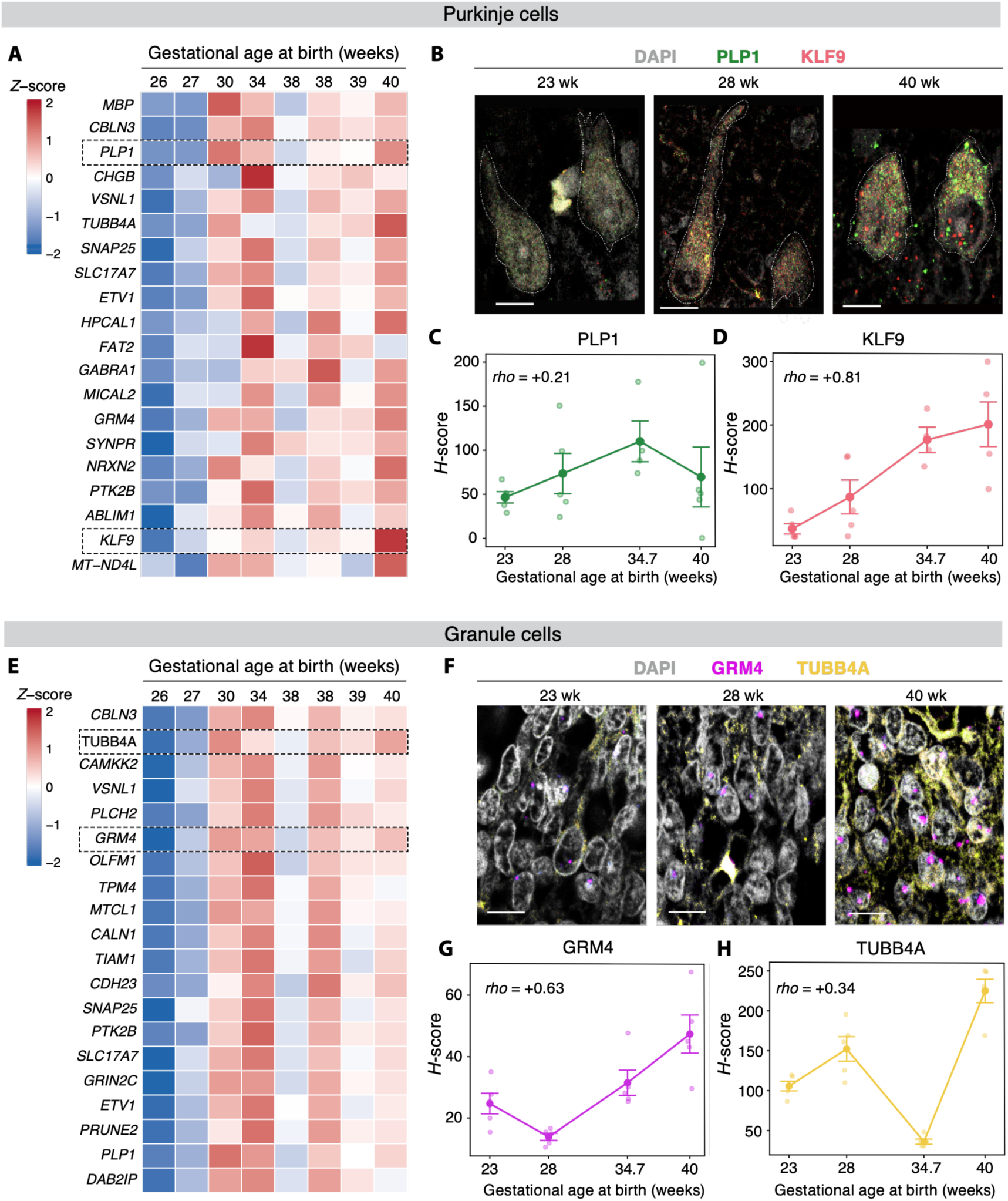
Lineage-specific gene expression marks gestational age-dependent cerebellar maturation. **(A)** Heatmap of Purkinje cell transcripts with the strongest positive gestational-age coefficients from linear mixed-effects models in post-menstrual age–matched spatial transcriptomic samples (*n* = 8; tables S6 and S7). Values are *Z*-scored mean expression per subject, ordered by gestational age at birth. Dashed boxes indicate genes selected for in situ validation. **(B)** Representative RNAscope images of Purkinje cells from infants born at 23, 28, and 40 weeks, showing *PLP1* (green), *KLF9* (magenta), and DAPI (gray). Scale bar, 10 µm. **(C and D)** RNAscope H-scores for *PLP1* (C) and *KLF9* (D) across gestational age at birth (*n* = 4 subjects; 4–5 regions of interest [ROIs] per subject). **(E)** Heatmap of granule cell transcripts with the strongest positive gestational-age coefficients from linear mixed-effects models, displayed as in (A). **(F)** Representative RNAscope images of granule cells from infants born at 23, 28, and 40 weeks, showing *GRM4* (magenta), *TUBB4A* (yellow), and DAPI (gray). Scale bar, 10 µm. (G and H) RNAscope H-scores for *GRM4* **(G)** and *TUBB4A* **(H)** across gestational age at birth, as in (C and D). In (C), (D), (G), and (H), each dot represents one ROI; large symbols indicate subject means, error bars denote SEM across ROIs, and lines connect subject means. Spearman rho between subject-mean H-score and gestational age at birth is shown in each panel.

Gestational age–associated expression patterns were inversely related to chronological age (fig. S18C–F), a relationship inherent to the post-term-equivalent sampling design, in which the most premature infants required the longest ex-utero survival to reach the study endpoint.

In the Purkinje cell lineage, transcripts increasing with gestational age at birth marked late cellular maturation and circuit stabilization (Fig. 4A). To validate these readouts in situ, we selected transcripts based on biological relevance and signal robustness. We assessed *KLF9*, a regulator of late Purkinje cell maturation and survival (*19*), and *PLP1*, a myelination-associated gene marking circuit-level maturation within the Purkinje cell layer (*20, 21*) (Fig. 4B). Both transcripts showed lower expression in extremely preterm cerebella and greater expression at later gestational ages (Fig. 4B–D), supporting their use as gestational age–associated molecular readouts of Purkinje maturation and indicating delayed circuit stabilization after preterm birth.

Within the granule cell lineage, genes correlated with gestational age were related to synaptic signaling and cytoskeletal stabilization (Fig. 4E). To capture this maturational transition in situ, we measured *GRM4*, a marker of parallel fiber synaptic signaling (*22*), and *TUBB4A*, a mediator of cytoskeletal maturation in granule neurons (*23*) (Fig. 4F). Both markers showed lower expression at earlier gestational ages and greater expression at later gestational ages (Fig. 4F–H). These results support delayed IGL progression toward a synaptically integrated and structurally stabilized state following extreme preterm birth.

Together, these analyses resolve continuous, lineage-specific readouts of gestationally ordered maturation programs and identify candidate molecular markers of postnatal circuit maturation.

## Discussion

Preterm birth reshapes the developing brain well beyond the neonatal period (*24*), (*25*) yet how the premature transition to extrauterine life influences the intrinsic cellular and molecular programs that orchestrate maturation remains unknown. By pairing in vivo developmental trajectories with an extensive collection of human postmortem cerebellar tissue, we traced the imprint of prematurity across scales, revealing a distinct ontogenetic program in which gestational age at birth shapes the baseline developmental configuration. This concept challenges a central assumption in prematurity research. After preterm birth, extrauterine development is not merely an incomplete third trimester modified by injury and neonatal complications, nor does it proceed independently of them (*26*). Rather, it reflects an interaction between the developmental state at birth and the physiological conditions under which maturation proceeds.

Our data support displacement of the cerebellar developmental trajectory, with the preterm cerebellum remaining stratified by gestational age at birth from early postnatal life through term-equivalent age across independent cohorts. Infants matched at this time point are therefore not biologically equivalent, as post-menstrual equivalence represents a chronological benchmark rather than a shared developmental endpoint. The persistence of this gestational ordering across clinical heterogeneity indicates that gestational age and neonatal morbidity are not competing explanations: birth timing influences the developmental context in which subsequent exposures act.

The observed cerebellar developmental divergence was both gestationally graded and lineage specific. While term Purkinje cells undergo a programmed reduction in density with advancing age, preterm Purkinje cells exhibit a paradoxical persistence, coupled with attenuated cellular refinement consistent with impaired synaptic pruning (*27*), (*28*). This persistence may reflect a molecular bioenergetic constraint in which the metabolic demands of pruning and chromatin remodeling outpace the capacity of the extrauterine environment, particularly as the cessation of placental support intersects with the energetic demands of extrauterine adaptation and rapid cerebellar development (*29*). The same interval exposes these circuits to premature sensory solicitation. Extrauterine life recruits cerebellar cortical circuits while neurogenesis and afferent refinement remain active, placing additional demand on climbing fiber–Purkinje cell connections whose maturation is itself pruning dependent (*26*), (*27*). Energetic constraint and premature activity may therefore converge on the same synaptic compartment in which we observe attenuated refinement, in the cerebellar cortex’s sole output neuron, offering a candidate route to the motor and cognitive phenotypes seen in preterm survivors (*30*), (*25*).

Granule cells, which provide critical trophic cues for Purkinje cell maturation, remain in an intermediate postmitotic state that may limit the synaptic integration required to drive refinement (*31*), (*32*). This immaturity may be compounded by the physiological transition to extrauterine life, which alters oxygen delivery, perfusion, nutrient supply, and sensory activity while cerebellar vascularization and circuit assembly remain ongoing (*33*), (*26*). These coincident demands could further modulate granule-cell progression and reduce the trophic and synaptic support available for Purkinje cell refinement, reinforcing a model of disrupted co-maturation across lineages rather than isolated loss of a single population.

These reciprocal lineage interactions offer a cellular level at which prematurity may redirect cerebellar maturation. Cerebellar development relies on the synchronized progression of the Purkinje and granule lineages, which exchange the trophic and synaptic cues each requires to advance (*31*), (*34*). Accordingly, maturation is reflected less by physical growth alone than by how far these programs have progressed through biosynthetic and synaptic assembly. Because the phenotype reflects impaired coordination rather than destruction of either lineage, the preterm cerebellum can retain its overall architecture while its developmental trajectory diverges. A programmatic shift of this kind may be incompletely captured by volumetric imaging, which is sensitive to macroscopic size but not to molecular state.

Because this divergence is organized by developmental state rather than chronology alone, it also reframes the context in which neuroprotective interventions act. Therapies delivered around preterm birth, including antenatal magnesium sulfate and postnatal caffeine, encounter a developmental substrate that varies with gestational age at exposure (*35*), (*36*). Viewed through this framework, their established benefits raise a broader question: whether the effect of an intervention depends not only on what is delivered, but also on the maturational state of the tissue receiving it. The gestationally ordered programs identified here may therefore help define that state and provide a basis for aligning intervention timing with biological maturity.

Although gestational age at birth shapes the initial developmental trajectory, biological insults associated with early delivery, including inflammation, hypoxia, and metabolic stress, together with broader postnatal influences such as the extrauterine environment and sensorimotor experience, modulate its subsequent course (*37*), (*26*), (*38*), (*39*). These converging influences cannot be fully disentangled in human cohorts, and our study therefore serves as a discovery platform that defines the developmental architecture associated with preterm birth and provides a human reference for mechanistic testing. Cerebellar tissue cannot be sampled in vivo, necessitating integration across separate molecular and imaging cohorts rather than within subjects. Volumetric MRI is cross-sectional and does not include formal brain-age gap estimation, although longitudinal ultrasound and neurodevelopmental follow-up position these findings along a developmental trajectory. The persistence of gestational-age stratification through term-equivalent age does not establish its permanence; whether it persists, attenuates, or undergoes compensation later in development remains unknown. The same protracted maturation that confers vulnerability may also extend the window for experience-dependent plasticity, during which developmental course remains modifiable.

Across independent human cohorts, molecular, histological, and imaging data converge on a common principle: extrauterine cerebellar maturation remains organized by the developmental state at birth. This principle shifts the translational focus from elapsed time to the biological maturity of the cerebellum at the time of assessment or intervention. Post-menstrual age, the benchmark against which preterm infants are assessed, monitored, and enrolled, indexes chronology rather than developmental state, and interventions timed to it may act on developmentally non-equivalent substrates.

Prematurity should therefore be understood not solely as a condition of vulnerability to injury, but as a state of biologically displaced maturation, in which the challenge is less to compensate for a lost intrauterine environment than to decode the developmental substrate that ex utero survival creates.

## Supporting information

Supplemental Material for manuscript

## Acknowledgments

We thank the CNH and NIH NeuroBioBank for providing postmortem cerebellar tissue and the families who consented to autopsy. We honor the memory of the deceased infants whose profound contribution has advanced our understanding of human cerebellar development; through their gift, we strive to improve the quality of life for future generations of children.

## Author contributions

G.S. and G.Si. contributed equally to this work. G.S. and G.Si. conceived and designed the study, performed experiments, analyzed and interpreted data, and wrote the manuscript. J.G. performed experimental work and contributed to data acquisition. K.W., R.V., G.S.-O’B., and D.N.S. performed bioinformatic analyses and computational modeling. C.L., R.C., H.C., and A.V. curated clinical data and contributed to cohort composition and clinical data extraction. D.B. contributed to data acquisition. F.A., S.S., and K.M. performed neuroradiological assessment and cerebellar volumetric evaluation. H.S.P. developed and implemented the deep learning–based neuropathological segmentation pipeline. M.T., N.W., and I.K. contributed to data acquisition. A.P. contributed to manuscript preparation. V.G. supervised the study and contributed to manuscript revision. P.K. conceived and supervised the study, designed experiments, interpreted results, and wrote the manuscript.

## Data and materials availability

All data underlying the figures and tables are provided with the article as data S1 to S5: in vivo clinical, neuroimaging, and neurodevelopmental data (data S1); postmortem histology and quantitative neuropathology data (data S2); processed spatial transcriptomic analysis tables (data S3); spot-level CoGAPS pattern weights and metadata (data S4); and in situ validation data (data S5). Subjects are identified only by study identifiers. All analysis code and scripts are available at https://github.com/kratimenoslab/Human-Preterm-Cerebellum and are permanently archived at Zenodo (*40*). No new materials were generated in this study.

## Supplementary materials

Materials and methods

Figs. S1-S18

Tables S1-S20

Data S1-S5 References (*41–68*)

## Supplementary Materials for

### This PDF file includes

– Materials and Methods
– Figs. S1 to S18
– Tables S1-S10 and tables S13-S20

### Other Supplementary Materials for this manuscript include the following

– Tables S11 and S12 (Excel files)
– Data S1. In-vivo cohort data (Excel file)
– Data S2. Postmortem cohort and histology data (Excel file)
– Data S3. Spatial transcriptomics analysis tables (Excel file)
– Data S4. Spot-level CoGAPS pattern weights and metadata (ZIP file)
– Data S5. In-situ validation data (Excel file)

## Materials and Methods

### I. Experimental design

We designed this study to determine whether premature birth redefines the intrinsic programs that regulate postnatal cerebellar development. We integrated data from complementary in vivo and postmortem human subjects to examine the relationships among birth timing, cerebellar structure, molecular state, and neurodevelopmental outcome. In vivo, we characterized cerebellar growth using term-equivalent magnetic resonance imaging (MRI), serial cranial ultrasonography, and standardized developmental follow-up assessments across gestational age strata. In parallel, we used spatial transcriptomics to identify gestational age–associated cerebellar programs, which were then evaluated in independent postmortem cohorts by whole-slide neuropathological analysis, RNAscope, and immunofluorescence. Across modalities, our design separated discovery from validation to determine whether gestational age–linked molecular programs were recapitulated in cerebellar architecture and lineage maturation.

#### Definitions

Gestational age (GA) refers to the estimated age of the fetus at birth, expressed in completed weeks from the first day of the last menstrual period. Chronological age (CA) refers to the time elapsed since birth. Post-menstrual age (PMA) is the sum of gestational age and chronological age (PMA = GA + CA) and represents the infant’s total age measured from the first day of the last menstrual period. Corrected age refers to chronological age adjusted for prematurity (corrected age = CA − [40 weeks − GA]) and was used to assess neurodevelopmental outcomes, accounting for the shorter gestational duration of preterm infants. Term-equivalent age (TEA) refers to a PMA of 37 to 42 weeks and denotes the developmental window at which preterm infants reach the maturational equivalent of full-term birth.

### **II.** In Vivo Neuroimaging and Neurodevelopmental Assessment

#### Ethics

This study was approved by the Institutional Review Board of Children’s National Research Institute (protocol #00011850) and conducted in accordance with institutional guidelines.

#### Study design and cohort

The in vivo cohort comprised 176 preterm infants, classified by gestational age at birth as extremely preterm (< 28 weeks), very preterm (28 to < 32 weeks), or late preterm (32 to < 37 weeks). Subjects contributed to one or more modalities: term-equivalent age (TEA) MRI with analyzable cerebellar volumetric segmentation (*n* = 120), serial cranial ultrasonography (*n* = 146), and standardized neurodevelopmental follow-up (*n* = 145) – with partial overlap across modalities (fig. S1). By modality, group composition (extremely/very/late preterm) was: MRI, 65/37/18 (*n* = 120); ultrasound, 80/45/21 (*n* = 146); neurodevelopmental, 94/51 (*n* = 145, extremely and very preterm only). Baseline and demographic metadata are in table S1.

#### Eligibility

The cohort was designed to isolate cerebellar developmental trajectories from the confounding effects of major destructive brain injury and congenital disease. We therefore excluded infants with major destructive brain injury (high-grade intraventricular hemorrhage [grade III/IV], post-hemorrhagic hydrocephalus, cerebellar hemorrhage, or cystic periventricular leukomalacia) or a congenital or genetic disorder. Genetic evaluation followed standard-of-care protocols, directed by clinical indication and comprising karyotyping and chromosomal microarray, with exome or genome sequencing and formal clinical-genetics assessment where indicated. Of 946 infants screened, 99 born at term (≥ 37 weeks) were excluded; of the remaining 847 preterm infants, 671 were excluded for absence of a TEA MRI (*n* = 403) or incomplete follow-up / clinical exclusion (*n* = 268), yielding the in vivo cohort of 176. The full selection process is shown in fig. S1.

#### Cohort derivation by modality

MRI, all 176 infants underwent TEA MRI; 56 scans were excluded from volumetric analysis (30 lacking adequate 3D T1-/T2-weighted sequences; 24 with motion or reconstruction artifact or supratentorial lesions precluding registration; 2 with post-menstrual age at scan outside the TEA window), yielding 120 with analyzable cerebellar volumes. Ultrasound, cranial ultrasound was available for 149 infants; 3 were excluded for incomplete longitudinal scanning, yielding 146. Neurodevelopment, developmental follow-up is not standard of care for late-preterm infants, and available late-preterm data represent a referral-biased subset; accordingly, follow-up analysis was restricted to extremely and very preterm infants. Of 176 infants, 1 lacked a valid assessment within the corrected-age window and 24 late-preterm infants were excluded (*n* = 151); a further 6 with only an overall delay classification and no domain-specific sub-scores were excluded, yielding 145 infants with analyzable domain data (motor, *n* = 143; cognitive, *n* = 140; language, *n* = 143).

### Brain MRI

#### MRI acquisition

MRI was performed at Children’s National Hospital on 3.0-T Discovery MR750 scanners (GE Healthcare, Milwaukee, WI) using a 32-channel receive-only head coil. Structural images for cerebellar segmentation were acquired using a: i) T1-weighted 3D SPGR sequence with following parameters: repetition time = 6.74 ms, echo time = 2.48 ms, flip angle = 12°, voxel size = 1 × 1× 1 = 1 mm^3^. Matrix is 160 × 160 (square), and field of view = 16 cm, slice thickness = 1 mm; and ii) a T2-weighted 3D Cube (fast spin echo) sequence with the following parameters: repetition time = 3000 ms, echo time = 90 ms, isotropic voxel size = 1 × 1× 1 = 1 mm^3^. Matrix is 160 × 160 (square), and field of view = 16 cm, slice thickness = 1 mm. Infants were scanned during natural sleep using a standardized feed-and-wrap protocol without pharmacological sedation. Infants were fed before scanning, swaddled in a vacuum immobilization device, fitted with hearing protection, and monitored throughout the examination. All MRI images were reviewed by experienced neuroradiologists before analysis. Studies were excluded if artifacts prevented reliable identification of cerebellar boundaries or landmarks, including posterior fossa ghosting, substantial signal dropout, incomplete cerebellar coverage, or geometric distortion. All MRIs included in the final cohort met our predefined quality criteria.

#### MRI volumetric analysis

Structural MRI data were processed using the Developing Human Connectome Project (dHCP) neonatal structural pipeline (*41*), which requires 3D T1-weighted and T2-weighted images. Raw images underwent bias field correction, motion correction, and super-resolution reconstruction, followed by brain extraction. Tissue segmentation was performed using the Draw-EM (Developing brain Region Annotation with Expectation-Maximization) algorithm (*42*), which classifies voxels into major tissue classes, including cortical grey matter, white matter, and cerebrospinal fluid. Segmentation labels generated by the dHCP pipeline were each reviewed, and cerebellar masks were manually corrected in ITK-SNAP v4.4.0 (*43*) by experienced neuroradiologists. Cerebellar volumes were extracted from the corrected masks (fig. S2A).

#### Statistical Analysis

Associations between cerebellar volume and gestational age were assessed using multivariable ordinary least-squares regression, including GA (weeks) and PMA at MRI (centered), with HC3 robust standard errors. Partial correlation coefficients for gestational age independent of PMA were derived from model t-statistics, and effect size was quantified using Cohen’s f². Influential observations were assessed using Cook’s distance. PMA– and sex-adjusted cerebellar volume residuals were compared across gestational age groups using Kruskal-Wallis tests with Dunn’s post hoc comparisons. All volumetric analyses were performed in R.

### Cranial ultrasonography

#### Acquisition

Serial cranial ultrasonography was performed as part of routine clinical care (Philips EPIQ Elite, curved-array transducer) at the bedside through the anterior fontanelle, with mastoid fontanelle views for posterior fossa visualization. Standard clinical timepoints were within 72 h of birth, 7 days of life, 28 days of life or discharge, and term-equivalent age (TEA). Cerebellar measurements comprised anteroposterior diameter, height, and cross-sectional area in the midsagittal plane, and transverse diameter, anteroposterior diameter, and cross-sectional area in the coronal plane (fig. S2B). All measurements were performed by trained raters blinded to MRI findings, gestational age at birth, and neurodevelopmental outcomes.

#### Composite size scores

Plane-specific composite scores were computed for the coronal (anteroposterior diameter, transverse diameter, and area) and midsagittal (vertical diameter, anteroposterior diameter, and area) planes by standardizing each measurement to a *Z*-score and applying inverse-correlation weighting to reduce redundancy. Pairwise Pearson correlations among the three measurements (cross-sectional area, anteroposterior diameter, and height) were computed across subjects; weights were set proportional to 1 − mean |*r*| for each measurement and normalized to sum to 1.

#### Growth trajectories

Longitudinal composite trajectories were modeled by linear mixed-effects regression with PMA, gestational age group, and their interaction as fixed effects and subject-specific random intercepts and slopes for PMA; interactions were tested by likelihood-ratio tests and pairwise slope differences estimated from marginal means with Tukey adjustment. Individual composite *Z*-scores were plotted against PMA for each gestational age group separately and combined (fig. S7A-B). Group-specific OLS slopes and their pairwise differences were compared using pooled standard errors with 95% confidence intervals. For subjects with ≥2 scans, the absolute change and rate of change (Δ*Z*/week) between first and last ultrasound were compared across groups by one-way ANOVA with Tukey HSD.

#### Cross-sectional comparisons

At 28, 33, and 35 weeks PMA (±2-week windows), the scan nearest each target age was used; because the 33– and 35-week windows overlap, nearest-scan selection ensured one observation per subject per timepoint, with no scan assigned to multiple timepoints. Groups were compared by Kruskal–Wallis and pairwise two-sided Wilcoxon rank-sum tests.

#### Deviation from normative fetal cerebellar growth

Transcerebellar diameter was referenced to the INTERGROWTH-21st international standards for fetal cerebellar growth (*11*) and expressed as a *Z*-score for PMA. Deviation was summarized as the maximal deficit at term-equivalent age, the rate of change in *Z*-score over PMA per gestational age group, and the proportion of measurements below the reference 5th percentile. Overall differences in trajectories between groups were tested using a linear mixed-effects model with fixed effects for PMA, group, and their interaction (PMA × group), a random intercept for participant, and Type III tests with Satterthwaite degrees of freedom (Fig. 1D).

#### Cross-modal validation

To assess whether bedside ultrasound reflects the volumetric gold standard, coronal and midsagittal ultrasound composites were correlated (Pearson) with the segmented 3D cerebellar volume from TEA MRI in infants with both examinations (*n* = 102). Because ultrasound preceded the MRI by a median of 4 weeks, attenuating the association across the full cohort, the comparison was performed in the well-matched subset scanned within 2 weeks of the MRI (*n* = 35; fig. S6A-B).

### Neurodevelopmental assessment

Neurodevelopmental outcomes were assessed at the Children’s National Hospital Neonatal Neurodevelopmental Follow-Up Clinic between 9 and 36 months corrected age. Standardized instruments included the Bayley Scales of Infant and Toddler Development, Third Edition (Bayley-III), and the Ages and Stages Questionnaire, Third Edition (ASQ-3) when a formal Bayley assessment was not feasible. Bayley assessments provided composite scores for the cognitive, motor, and language domains, and corrected age was applied according to standard clinical practice.

#### Delay classification

Outcomes were defined as clinician-adjudicated binary classifications of developmental delay (yes/no), assigned at the time of evaluation by neonatal follow-up specialists. These classifications were derived from standardized instrument results, with Bayley composite scores < 85 used to define delay and ASQ-3 results interpreted using established age-specific cutoffs. All assessments were administered by certified developmental examiners blinded to neonatal neuroimaging measurements.

#### In vivo cohort sex analysis

Sex was evaluated as a covariate in the in vivo cohort (*n* = 176; extremely preterm: 41 females, 57 males; very preterm: 18 females, 36 males; late preterm: 10 females, 14 males). Associations of sex with gestational age group and continuous gestational age were assessed using a *χ*² test and two-sided Welch’s *t* test, respectively (table S3). Cerebellar MRI measures were analyzed by ordinary least-squares regression including centered PMA at MRI, gestational age, sex, and a gestational age × sex interaction. Longitudinal ultrasound growth was analyzed by linear mixed-effects regression, with PMA × sex tested by likelihood-ratio test. Developmental delay was modeled by logistic regression including gestational age group and sex; in infants with MRI and neurodevelopmental data (*n* = 98), cerebellar volume and the cerebellar volume × sex interaction were additionally evaluated. Analyses were performed in R v4.5 using lme4 and lmerTest, with figures generated using ggplot2.

#### Statistical analysis

Rates of developmental delay were compared using Fisher’s exact test, with confidence intervals estimated by the Wald approximation (truncated to 0–100%). To assess potential confounding by plurality, twins were compared with singletons using Wilcoxon rank-sum and Fisher’s exact tests, and regression models were repeated with twin status included; no significant differences were observed, and effect estimates were unchanged (fig. S5). To relate cerebellar structure to outcome, the volumetric MRI cohort was merged with the neurodevelopmental cohort (extremely and very preterm only, gestational age < 32 weeks), restricted to subjects with both MRI and neurodevelopmental data and valid domain-specific delay classifications; point-biserial correlations were computed between cerebellar volume and binary delay status for each domain (motor, cognitive, language), and cerebellar volumes were compared between the delay and no-delay groups using Wilcoxon rank-sum tests (Fig. 1C). All analyses were performed in R (v4.5.2) using lme4, emmeans, DescTools, ggplot2, and patchwork.

### Perinatal and neonatal clinical factors and covariate-adjusted analyses

To determine whether the gestational-age associations reflected birth timing itself or the co-occurring morbidities of prematurity, each infant’s perinatal and neonatal intensive-care course was phenotyped across six domains: respiratory, infectious, gastrointestinal, glycemic, procedural pain, and delivery indication, together with two pharmacologic exposures (antenatal magnesium sulfate and postnatal caffeine) (fig. S4A-B; table S2; table S4; table S5). Respiratory support was summarized by the respiratory severity score (RSS; daily mean airway pressure × FiO₂) and its cumulative form (daily RSS × days of invasive ventilation), with ventilation days summed by mode from respiratory flowsheets. Infectious and inflammatory exposure was summarized by days of therapy (DOT; calendar days receiving ≥1 antimicrobial), per-agent antibiotic days, and peak and mean C-reactive protein over the observation window.

Gastrointestinal morbidity was represented by surgical necrotizing enterocolitis. Procedural pain was quantified from the Premature Infant Pain Profile (PIPP) as the mean, median, and cumulative score and the proportion of assessments above threshold; glycemic control was expressed as the proportion of glucose readings within, below, and above the 70–180 mg/dL range (time in, below, and above range). Models used [cumulative PIPP score] and [time in range], respectively. Delivery indication distinguished infants delivered for a maternal indication (for example, preeclampsia or placental abruption) from those born after spontaneous onset of labor. Respiratory and infectious morbidity were additionally represented by study-defined composite measures: the cumulative burden of bronchopulmonary dysplasia (cumulative RSS and duration of mechanical ventilation) and the cumulative burden of sepsis (peak C-reactive protein, antibiotic days of therapy, and blood-culture positivity). Surgical necrotizing enterocolitis, magnesium sulfate, and caffeine exposure were recorded as categorical variables.

Because neonatal morbidities are mutually correlated and covary with the degree of immaturity, each exposure was entered individually alongside gestational age rather than in a single multivariable model, and these were treated as covariate-adjusted sensitivity analyses without correction for multiple comparisons. For each association, the independent contribution of gestational age was assessed by nested-model comparison. For cerebellar volume, an ordinary least-squares model including post-menstrual age at MRI and the exposure was compared with a model additionally including gestational age, and the gestational-age term was tested by a nested F-test. For developmental delay, a logistic model including the exposure was compared with one additionally including gestational-age group, tested by likelihood-ratio χ². For longitudinal ultrasound growth, a linear mixed-effects model tested the post-menstrual age × gestational-age-group interaction by likelihood-ratio test under adjustment for each exposure. The relationship between cerebellar volume and developmental delay was evaluated by logistic regression with cerebellar volume (per 1,000 mm³) adjusted for each exposure. Association retention was defined as *P* < 0.05 for the gestational-age (or cerebellar-volume) term. Analyses were performed in R (v4.5).

### **III.** Postmortem Tissue Analysis

#### Ethics

This study was conducted in accordance with protocols approved by the Institutional Review Board of Children’s National Research Institute (IRB protocol #00015350). All postmortem subjects were derived from the Children’s National Hospital pathology department and the NIH NeuroBioBank.

#### Cohort organization

Postmortem cerebellar tissue cohorts were stratified by analytic modality and assembled by directed selection from institutional archives, as is standard for human postmortem tissue studies. Subjects were selected on the basis of gestational age, post-menstrual age at death, tissue availability, and preservation quality. Cohort composition by modality was as follows:

#### Eligibility

To parallel the in vivo cohort design and isolate cerebellar developmental trajectories from confounding pathology, subjects with major destructive brain injury (high-grade intraventricular hemorrhage [grade III/IV], post-hemorrhagic hydrocephalus, cerebellar hemorrhage, cystic periventricular leukomalacia, or hypoxic-ischemic encephalopathy) or major congenital or genetic anomalies were excluded, as were cases with significant autolysis, prolonged postmortem interval, or insufficient analyzable cerebellar tissue. Clinical comorbidities, including necrotizing enterocolitis, sepsis, and maternal pre-eclampsia, were documented for each subject (table S7; table S11; table S12). Of 178 autopsy cases screened, 95 were excluded on these criteria, yielding a postmortem cohort of 83 subjects (fig. S1).

i) *Spatial transcriptomic analysis*: fixed-frozen cerebellar tissue from 12 subjects (seven preterm, five term) obtained from the NIH NeuroBioBank. Subjects were selected to approximate equivalent post-menstrual age at death across preterm and term groups. One term subject was excluded from downstream analysis because advanced post-menstrual age at death (> 200 weeks) introduced age-related confounding, yielding a final analytic cohort of 11 subjects. Two sections were analyzed per subject (22 sections total; table S6; table S7).
ii) *Neuropathological analysis*: a paraffin-embedded cerebellar tissue cohort of 77 infants (preterm, *n* = 57; term, *n* = 20; table S10; table S11; table S12). This cohort does not overlap the in vivo cohort and shares five subjects with the spatial transcriptomic cohort (Preterm 1, 2 and 52; Term 1 and 3; table S7). Gestational age at birth ranged from 22.1 to 40.0 weeks, chronological age at death from 0.0 to 56.4 weeks, and post-menstrual age at death from 24.6 to 93.3 weeks. Layer measures were plotted against gestational and chronological age (Fig. 3) and regressed on gestational age at birth in the sex and clinical covariate analyses (fig. S16; table S13; table S14).
iii) Validation: a subset of the neuropathological cohort comprising 12 unique subjects, 4 assessed by RNAscope in situ hybridization (table S15) and 9 by immunofluorescence (table S16), with one subject assessed by both. Immunofluorescence was performed in 8 subjects per marker; because the term subject differed between markers, 9 specimens contributed in total. Validation subjects, which spanned the full gestational age range, were used only to confirm patterns identified in the spatial-transcriptomic discovery analysis.

### Spatial Transcriptomics

#### Tissue acquisition and processing

Fixed-frozen cerebellar tissue was collected at postmortem examination. After legal pronouncement of death, an autopsy was typically initiated within 24–48 hours to limit postmortem degradation, with the postmortem interval reported for all tissue collected within this timeframe. Sections were sampled using a standardized landmark-based strategy to ensure anatomical comparability across subjects. Tissue was embedded in optimal cutting temperature compound, cryosectioned at 10 µm, mounted onto Visium capture slides, and processed according to each manufacturer’s protocols. Sections were stained with hematoxylin and eosin (H&E) before library preparation. Six Visium slides were used, each containing four capture areas. Libraries were generated using the 10× Genomics Visium Spatial Gene Expression reagent kit and sequenced on an Illumina NovaSeq 6000 with 150-bp paired– end reads, targeting 50,000 reads per spot.

#### Data processing

Clinical metadata were abstracted from electronic medical records and included gestational age at birth, chronological age, and post-menstrual age at death. Spatial annotations were derived independently from histological compartmentalization and canonical marker gene expression and were not informed by downstream computational outputs. Subject-level clinical information is provided in table S7. Raw sequencing data were processed using Space Ranger (v1.3.0) with alignment to GRCh38. Gene expression matrices were imported into Seurat (v4.0.1) in R for downstream analysis (*44*). Data were normalized using log normalization, followed by principal component analysis and graph-based clustering. Pathology annotations were incorporated into the Loupe output and used together with expression-based clustering to define spatially distinct compartments and transcriptional states. Differential gene expression was evaluated using a pseudobulk framework implemented in Seurat. Pseudobulk count matrices were modeled in DESeq (*45*) with batch included as a covariate in the design formula; limma (*46*) was used separately to visualize expression values after batch correction. Projection of cell type and developmental reference signatures onto the spatial data was performed using projectR. Analyses were conducted in R (v4.1).

#### Batch structure

24 capture areas from 12 subjects were distributed across six Visium slides. One subject was subsequently excluded, leaving 22 capture areas from 11 subjects in the final analysis. Slide assignments were balanced across gestational age groups to minimize confounding between batch and developmental stage. Batch was included as a covariate in all differential expression models.

#### Visium quality control

Quality control was performed independently on each of the 24 capture areas. Spots were filtered based on three metrics: (1) a minimum total unique molecular identifier (UMI) count of 500 to exclude empty or near-empty spots, (2) a minimum gene count of 250 to remove low-complexity spots, and (3) a maximum mitochondrial gene fraction of 10% to exclude spots overlying damaged or necrotic tissue. These thresholds were applied across all samples and were determined from per-sample knee plots, UMI and gene count histograms, and UMI-versus-gene scatter plots colored by mitochondrial fraction (see Visium quality control, below). Of 74,096 total spots across all 24 capture areas (including the excluded term subject), 2,000 (2.7%) failed at least one quality control (QC) criterion. After QC filtering and removal of the excluded subject’s two capture areas, 72,096 spots across 22 capture areas from 11 subjects were retained for downstream analysis. After filtering, the median UMI count per spot was 10,611, the median number of genes per spot was 4,610, and the total number of UMIs across all retained spots was approximately 834 million.

#### CoGAPS pattern discovery and annotation

Transcriptional pattern discovery was performed using CoGAPS, a Bayesian non-negative matrix factorization framework, applied to the aggregated Visium dataset as previously described (*47*). CoGAPS decomposes the log-transformed expression matrix into an amplitude matrix that captures gene weights and a pattern matrix that captures spot-level activity. Prior to factorization, genes with zero counts across all spots were removed, reducing the feature space from 36,601 to 29,963 genes. Unsupervised factorization identified 28 transcriptional patterns (H-1 to H-28). Three patterns termed Cerebellar Maturation-Associated Patterns (CMAPs) were selected for in-depth characterization and are referred to by their functional annotations throughout: H-16 (CMAP_IGL_IMM), H-17 (CMAP_IGL), and H-27 (CMAP_PCL). Pattern prioritization required concordance across three independent criteria: (i) Spearman rank correlation of pattern activity (P matrix) with developmental timing variables, including gestational age and/or post-menstrual age at death; (ii) inspection of top-weighted genes in the amplitude matrix (A matrix) for coherent biological functions rather than nonspecific gene sets; and (iii) spatial localization of pattern activity consistent with known cerebellar cortical compartments and cellular lineages.

Patterns were excluded in two stages. First, H-1, H-3, H-4, H-5, H-10, H-13, H-15, H-20, H-23, and H-25 were excluded because they failed to meet the three prioritization criteria above. After the first exclusion pass, remaining patterns were evaluated a second time prior to downstream analysis. Patterns lacking coherent developmental associations, spatial localization to cortical compartments, or biologically interpretable gene expression content, as well as those driven primarily by expression or correlation in two age-discordant subjects, were excluded from individual characterization. All patterns remained in the CoGAPS factorization and contributed to downstream analyses, but only the three prioritized CMAPs were characterized individually.

No clinical or spatial metadata were included during CoGAPS model fitting. Post hoc interpretation of patterns used only gestational age at birth, post-menstrual age at death, and spatial laminar annotations. Spatial laminar annotations were generated by integrating manual neuropathologist delineation on H&E-stained tissue images with expression-based clustering of canonical cell-type markers, independently of CoGAPS outputs. Pattern annotations were confirmed by visualizing spatial feature plots of canonical lineage markers alongside pattern activity maps. Feature plots were generated using Seurat’s SpatialFeaturePlot function.

#### Clinical subgroup analysis

Clinical variables for the 11 spatial transcriptomic subjects were coded as binary indicators from autopsy, neuropathology, and neonatal records; for two subjects no primary record was available and coding relied on donor-bank metadata. Subject-level CMAP activity was correlated with gestational age at birth and, separately, with post-menstrual age at death by Spearman rank correlation, first in the full cohort and then within each clinical subgroup. Subgroups of fewer than five subjects were not analyzed. Correlations are descriptive; no *P* values were computed (table S9).

#### Functional annotation (fGSEA)

To characterize the molecular programs captured by CoGAPS patterns CMAP_IGL_IMM (H-16), CMAP_IGL (H-17), and CMAP_PCL (H-27), we performed gene set enrichment analysis using genes ranked by their raw amplitude weights from the CoGAPS A matrix (positive-score type). Gene sets were obtained from MSigDB category C5 (Gene Ontology: Biological Process, Cellular Component, and Molecular Function; Human Phenotype Ontology terms excluded) via the R package msigdbr for *Homo sapiens* (*48*), and the analysis was run with the R package fgsea (v1.28.0) using default parameters and Benjamini-Hochberg-adjusted *P*-values. Each ranked gene list was tested against 10,395 Gene Ontology terms. For each pattern, gene sets were ranked by enrichment *P*-value, and the top 50 terms per pattern were selected and combined into a single union set by removing duplicate terms shared across patterns, enabling cross-pattern comparison without censoring enrichment to a single pattern. Gene sets were ordered by the pattern in which they ranked most significantly (H-16 first, then terms unique to H-17, then H-27) and displayed across all three patterns as a heatmap using a double-log transformation (log_10_(−log_10_[nominal *P* value])) with a diverging blue-red color scale (ComplexHeatmap; colorspace Blue-Red 3 palette) (*49*).

#### Partitioned Heritability (LDSC)

We tested whether cerebellar gene expression patterns were enriched for genetic variants associated with cerebellar/neurodevelopmental traits using LD score regression (LDSC) (*50, 51*). For each of the 28 CoGAPS patterns, genes with amplitudes above the 90th percentile were mapped to genomic coordinates (hg38) using ±100-kb regulatory windows. All 28 patterns were included in the analysis to maintain a conservative multiple-testing denominator; results are reported for the three prioritized CMAPs. The datasets included in these analyses are listed in table S8. These regions were converted to binary annotations for LD score computation using 1000 Genomes Phase 3 European ancestry reference genotypes (*52*), conditioned on the baseline LD model v2.2 (97 functional annotations) (*53*). We performed partitioned heritability regression for 76 traits drawn from the studies listed in table S8, spanning psychiatric and neurological disorders, cognitive and motor traits, and brain imaging phenotypes (*54, 55, 56, 57, 58, 59, 60, 61, 62, 63, 64*). Enrichment was defined as the proportion of single-nucleotide polymorphism (SNP) heritability explained by a pattern divided by the proportion of SNPs in that pattern. Significance was assessed using Benjamini-Hochberg false discovery rate (FDR) correction across 2,128 tests (28 patterns × 76 traits), with FDR < 0.05 considered significant.

#### Tricycle cell-cycle analysis

We applied Tricycle analysis, as previously described in (*65*), to evaluate cell-cycle phases at the spot level using the same cohort used for the Visium spatial transcriptomic analysis. Briefly, Tricycle employs transfer learning to predict continuous cell-cycle position; originally developed for single cell RNA-seq, it was applied here at Visium spot resolution (∼55 µm). The inferred position, represented as an angle ranging from 0 to 2π, denotes the location of a spot within continuous cell cycle progression. Spots were assigned to CMAPs based on pattern weights; for each pattern, we selected the spots whose weights exceeded the pattern’s 95th percentile to enrich for high-confidence assignments. Within each group (extreme preterm, very/late preterm, and term), we estimated phase distributions both per subject and for the pooled set of spots across all subjects in each category, by computing kernel density estimates along the Tricycle continuum.

#### Linear model of gene expression

Linear mixed-effects models were used to assess the relationship between gene expression and gestational age. For this analysis, the spatial transcriptomic cohort was restricted to 8 subjects (4 preterm, 4 term) with comparable PMA at death, between 45 and 52 weeks. Demographic information is provided in table S6; table S7. Spots were assigned to Purkinje cells or granule cells based on independent histological laminar annotations derived from the Visium spatial maps; spots overlying the Purkinje cell layer were classified as Purkinje cell spots, and spots overlying the internal granule cell layer were classified as granule cell spots. Separate models were constructed for each gene in Purkinje and granule cells.

The model was specified as:

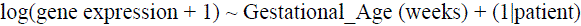

where gestational age was included as a fixed effect and patient as a random intercept to account for within-subject variability. This model was applied iteratively to each gene. For each gene, the fixed-effect coefficient β and its associated *t*-value were extracted. The 20 most strongly positively and the 20 most strongly negatively associated genes, based on the gestational age coefficient β, were selected separately for granule cells and Purkinje cells. For visualization, log-transformed expression values for each gene were averaged across spots of interest (granule cells or Purkinje cells) per patient, and the resulting values were scaled to *Z*-scores. Heatmaps of scaled expression values were ordered by direction and strength of association (genes) and by gestational age (samples), with the color scale clipped at the 5th and 95th percentiles to reduce the influence of outliers. To assess potential confounding by postnatal duration, the same genes were plotted against PMA at death, confirming the expected inverse relationship with gestational age across subjects under the study’s sampling design. All analyses were conducted in R using the lme4 package (v1.1-35).

### Deep-Learning–Based Neuropathological Analysis

#### Pipeline organization

The neuropathological cohort described above (*n* = 77) was analyzed using a deep learning–based histopathology pipeline. Neuropathological review assessed cellular morphology within the cerebellar layers: external granule cell layer (EGL), molecular layer (ML), internal granule cell layer (IGL), and the Purkinje cell (PC) layer. Whole-slide imaging of H&E-stained sections was performed using an Olympus VS120 microscope at 20× magnification. The entire tissue area on each slide was quantified using deep learning–based segmentation.

#### Tissue acquisition, processing, and standardization of section levels

Cerebellar specimens were obtained at autopsy at the Children’s National Hospital Department of Pathology following the same autopsy protocol described for the spatial transcriptomic cohort, with the following processing steps for formalin-fixed, paraffin-embedded (FFPE) samples. The brain was removed en bloc, weighed, inspected, and immediately immersed in 10% neutral-buffered formalin for at least 7 days to ensure uniform fixation. After fixation, the cerebellum was sharply separated from the brainstem at the level of the superior cerebellar peduncles and bisected precisely along the midline. For each subject, tissue blocks were collected from two consistent anatomical planes: (i) a parasagittal hemispheric section at the level of the dentate nucleus, used as a reproducible internal landmark, and (ii) a midsagittal vermian section. The parasagittal block was sampled from the first well-preserved ribbon immediately lateral to the vermis, corresponding anatomically to lobules VI-VIII, a region with highly conserved folial architecture. Consistency of findings across both parasagittal and vermian sections within each subject supports generalizability across cerebellar subregions. Tissue sections were processed as FFPE specimens according to CAP-accredited histopathology protocols. Fixed tissue underwent graded ethanol dehydration, xylene clearing, and paraffin infiltration under vacuum before being embedded in fresh paraffin with standardized orientation to maintain a perpendicular cutting plane across the folial axis. Blocks were stored in a temperature– and humidity-controlled archive until microtomy. Sections were cut at 5 µm thickness on a rotary microtome using ribbons generated orthogonally to the folial curvature to ensure consistent representation of the EGL, ML, IGL, PC layer, and deep white matter. Sections were floated on a 40°C water bath, mounted on positively charged slides, dried, and stained with H&E.

#### Segmentation workflow

We used the active-learning features and annotation import/export utilities of the NoCodeSeg pipeline to perform deep learning–based cerebellar layer and PC segmentation. For this study, the DeepMIB graphical training interface was replaced with a scripted Python/PyTorch ensemble-training implementation to improve scalability and reproducibility. Specifically, we trained an ensemble of three DeepLabV3+ models with ResNet-18 backbones, each with independent random initialization and stochastic augmentation settings. Ensemble predictions were fused to form a consensus segmentation mask, which was used for auto-labeling and iterative expansion of the training set. Low-confidence patches were reviewed and corrected by a pathologist using an interactive workflow in QuPath (*66*) and Microscopy Image Browser (MIB) (*67*). Corrected labels were then reintegrated into subsequent training cycles. This iterative human-in-the-loop refinement was repeated until segmentation performance stabilized across cycles (fig. S15A). Annotation export/import scripts were retrieved from the NoCodeSeg repository; data were processed using Python (pandas, numpy) for cleaning and formatting, and statistical modeling and visualization were performed in R.

#### Segmentation performance

The ensemble model was evaluated on a 20% held-out validation set constructed at the slide level and stratified by gestational age. Collapsing the four cerebellar layers into foreground versus background, quantitative performance was strong: median intersection over union (IoU) = 0.989 (interquartile range [IQR] 0.981–0.992; mean 0.976; *n* = 213 tiles) and median Dice = 0.994 (IQR 0.990–0.996; mean 0.983; *n* = 213).

Boundary-level accuracy was high, with a symmetric mean surface distance of 5.86 px (IQR 4.26–7.71; mean 6.30; *n* = 211). Topology-preserving accuracy, assessed using the topology-break rate (normalized skeleton connectivity mismatch between prediction and ground truth), yielded a median of 0.533 (IQR 0.242–1.000; mean 0.853; *n* = 213). Class-wise confusion-matrix analysis demonstrated high fidelity for individual cerebellar layers (EGL: 0.89, IGL: 0.98, ML: 0.98, white matter: 0.99 correct classification probabilities), confirming low cross-layer misclassification (fig. S15B–C). EGL misclassification occurred predominantly at the EGL–ML boundary in older specimens where the EGL is thinnest; all segmentations were reviewed and corrected by an expert neuropathologist before quantification. Model performance was consistent across cortical regions, indicating that accuracy was not driven by overfitting to a particular anatomical configuration. The DeepLabV3+ architecture was selected for its multi-scale feature extraction capability, which accommodates the variable thickness of cerebellar cortical layers across developmental stages.

#### Normalization

Because 2D sections can vary in the relative proportions of EGL, ML, IGL, and underlying white matter due to folial geometry and sampling angle, we applied analytic normalization to reduce confounding from section size and folial extent. Specifically, we quantified ML length on each slide (derived from the ML segmentation mask) as a surrogate for folial extent and normalized layer area measurements (EGL, ML, IGL) to ML length (mm²/µm). This normalization facilitates robust comparisons of cortical architecture across specimens despite unavoidable anatomic variability in fixed tissue.

#### External granule cell layer density

EGL cell density was quantified from whole-slide H&E, using the deep learning-derived EGL segmentation to define the layer region. Within each EGL region, cells were detected in QuPath (v0.6.0) using the built-in StarDist cell detection on the H&E stain, and EGL cell density was computed as the number of detected cells divided by the segmented EGL area (detections/mm²). For subjects with multiple sections, a single per-subject value was obtained by pooling detections and area across sections.

#### Purkinje cell quantification

Layer segmentations were used to define the Purkinje cell analysis region at the molecular layer–internal granule layer (ML–IGL) interface. An active-learning process analogous to that used for cerebellar layer segmentation was applied to PC segmentation, yielding a final dataset of 13,247 annotated patches. Final PC predictions were imported into QuPath. During workflow development, an intermediate 50-µm expansion was generated; this was subsequently replaced by a 100-µm expansion toward the IGL, which defined the final region used for quantification. The larger region provided a more accurate representation of the PC distribution by avoiding restriction of the analysis to the expected monolayer and ensuring that displaced or misaligned PCs were also captured. Based on the mean annotated PC profile diameter of 14.63 µm (median, 14.98 µm), the final region encompassed approximately seven PC soma widths while remaining anatomically centered on the ML–IGL interface. PC detections within this region were included in the analysis. PC-layer length was estimated by dividing the perimeter of the expanded ML annotation by two to account for its two approximately parallel boundaries. PC density was reported as the number of PCs per millimeter of estimated PC-layer length.

#### Statistical analysis

The neuropathological cohort spanned the full postnatal cerebellar developmental trajectory (PMA at death: 24.6–93.3 weeks for preterm; 39.1–86.4 weeks for term). Five cerebellar parameters were analyzed independently: EGL normalized surface area, EGL density, ML normalized surface area, IGL normalized surface area, and PC density. For subjects with multiple tissue sections, measurements were averaged to obtain a single representative value per subject. All 77 subjects (57 preterm, 20 term) were included in the primary analysis. Each parameter was examined using two complementary approaches: (i) linear regression against gestational age across all subjects to assess the overall developmental gradient, and (ii) Within each group, cerebellar measures were regressed on chronological age to estimate postnatal developmental trajectories, and slopes were compared between preterm and term subjects. Slope differences between groups were tested using a two-sample *t-*test of the group-specific ordinary least-squares slope coefficients and their associated standard errors, with statistical significance set at *P* < 0.05. Because the five parameters assess distinct biological compartments and were evaluated as independent hypotheses, no correction for multiple comparisons was applied.

#### Sex analysis

Sex was evaluated as a covariate in the postmortem cohort (*n* = 77; preterm: 28 females, 29 males; term: 8 females, 12 males). Associations of sex with preterm/term status and continuous gestational age were assessed using Fisher’s exact test and two-sided Welch’s *t* test, respectively. Cerebellar layer morphometry, Purkinje cell density, and external granule layer cell density were regressed on gestational age with and without sex. Gestational age × sex and sex × preterm/term interactions were tested. Analyses were performed in R v4.5, with figures generated using ggplot2.

### Clinical covariate analysis

Neonatal exposures were defined as described for the in vivo cohort (Section II) and abstracted from the medical record for subjects with available clinical records (fig. S16; table S14). Each cerebellar parameter, Purkinje cell density, external granule layer and molecular layer morphometry, and layer-specific cell densities, was regressed on gestational age with and without each exposure, and the independent contribution of gestational age was assessed by the same nested-model comparison. Because exposures were evaluated individually and the parameters assess distinct compartments, no correction for multiple comparisons was applied. Analyses were performed in R (v4.5).

### RNAscope and Immunofluorescence Validation

All validation tissue was obtained from the neuropathological cohort as described above (table S15; table S16). Cell segmentation for both RNAscope and immunofluorescence analyses was performed using InstanSeg (Fluorescence Nuclei and Cells 0.1.1) which identifies nuclei via DAPI staining and infers whole-cell boundaries, generating nuclear and cytoplasmic compartments. Following segmentation, cells were manually classified as EGL, PC, or IGL based on morphology and spatial localization. All measurements were exported from QuPath (v0.6.0) for downstream statistical analysis in R (v4.5), with figures generated using ggplot2.

#### RNAscope staining and detection

RNAscope in situ hybridization was performed on FFPE cerebellar sections from the 4 validation subjects described above (table S15), with four to five regions of interest (ROIs) acquired per subject. Multiplex RNA in situ hybridization was performed using the RNAscope Multiomic LS assay on the BOND RX automated staining platform (Leica Biosystems) according to the manufacturer’s protocol. Briefly, staining protocols were configured within the BOND Research Detection System using the ACD Amplification 6 workflow, including epitope retrieval (HIER; ER2, 20 min), protease permeabilization (PretreatPro), and RNAscope 2.5 LSx probe hybridization and amplification steps. Two sequential multiplex panels were run on adjacent sections from each subject, using eight probes (table S17; table S18). RNA probes were registered in the system, and reagent volumes were calculated to account for instrument dead volumes before loading into open containers. Slides were configured with probe identity and tissue metadata, barcoded, and processed through fully automated cycles of probe application, amplification, and detection, with total run times of 20– 26 hours. Upon completion, slides were removed, rinsed, and coverslip-mounted for imaging.

#### RNAscope image acquisition and quantification

Fluorescence images were acquired at 60× magnification using two complementary systems. Channels for DAPI, Opal 520, Opal 570, Opal 620, and Opal 690 were captured on a FLUOVIEW FV4000 with AF405, AF488, Cy3, AF594, and AF647 excitation wavelengths, respectively. To minimize spectral overlap with Opal 520, the Opal 480 channel was captured separately on a STELLARIS LIAchroic equipped with a tunable white light laser. Images from both systems were acquired at matched pixel sizes and co-registered before analysis. RNAscope probe panels targeted PC maturation genes (*PLP1, KLF9*), granule cell maturation and differentiation genes (*TUBB4A*, *GRM4*), and cell cycle genes (*MCM2, PCNA, CCNB1, TOP2A*). ROIs corresponding to cerebellar lobules were manually annotated, and channel identities were standardized with a custom script to match probe assignments. Per-cell measurements included morphological features and fluorescence intensities across nuclear, cytoplasmic, and combined compartments. Subcellular RNA puncta and clusters were detected via intensity thresholding and quantified per cell and compartment, including counts, areas, and fluorescence-intensity metrics.

#### Immunofluorescent staining

FFPE sections from the 9 validation subjects described above, matched for PMA at death (range 42.7–60.3 weeks; table S16), were processed using a standardized immunofluorescence protocol. Sections were baked at 60°C for 45 minutes, washed in xylene, graded ethanol, and 1× phosphate-buffered saline (PBS), and subjected to antigen retrieval in 10 mM citrate buffer (pH 6.0). After PBS-Tween rinses and 3% H₂O₂ incubation, sections were blocked with a commercial blocking solution (Akoya Bioscience, ARD1001EA) at room temperature and incubated with primary antibodies for 24 to 48 hours at 4°C. Antibody details are reported in table S19. Following rinsing, sections were incubated for 2 hours at room temperature with the appropriate secondary antibodies and mounted on 24 × 50 mm No. 1.5 coverslips (Fisherbrand, cat. 12-541-033) with DAPI (Vector Laboratories).

#### Immunofluorescence image acquisition and quantification

To assess excitatory synaptic inputs, cerebellar sections were immunolabeled for calbindin together with VGLUT1 or VGLUT2 and imaged on a Leica STELLARIS confocal microscope (LAS X software; HC PL APO CS2 20×/0.75 objective). Five regions of interest (ROIs) were acquired per subject; each marker was assessed in 8 subjects (7 preterm, 1 term); because the term subject differed between markers, 9 unique specimens contributed in total (gestational age 23–40 weeks).

Three channels were collected using the white-light and 405-nm lasers on Power HyD detectors: DAPI (405 nm), calbindin (Alexa ≈488, analog mode), and VGLUT1 or VGLUT2 (Alexa ≈647, photon-counting mode). Laser power, detector gain, and all acquisition settings were held constant across images to preserve quantitative comparability. Frames were acquired in xyz mode at 2048 × 2048 pixels (field, 581 × 581 µm; pixel size, 283 nm), 200 Hz, zoom 1.0, pinhole 1.0 AU, and 875-ns dwell time, with a five-point manual focus map per section. Images were exported as OME-TIFF and quantified in QuPath (v0.6.0). In each image, the molecular layer (ML) was manually annotated (blind to gestational age) and detected cells removed to restrict analysis to the nuclei-free ML. VGLUT1 and VGLUT2 were quantified using the same pixel-classifier thresholder applied to the respective channel at full resolution, with identical classifier parameters held constant across all images, and expressed as the positive-area fraction (marker-positive area divided by total ML area); the two markers were analyzed and reported separately. Images with a mean punctum area >100 µm², positive area >90%, or fewer than 10 detected puncta were excluded before analysis.

#### RNAscope and immunofluorescence statistical analysis

Per-cell RNA expression levels were scored using the H-score system (*68*) following the ACD/Bio-Techne RNAscope fluorescent multiplex assay scoring guideline (Advanced Cell Diagnostics, Technical Note MK-50-010), in which estimated spot counts per cell were categorized as negative (0 spots), low (1–3 spots), medium (4–9 spots), or high (10 or more spots). The H-score was derived from the weighted sum of the percentage of cells in each bin, yielding a continuous score ranging from 0 to 300. H-scores were computed for each gene and cell type combination, per ROI. This approach was applied to PC maturation genes (*PLP1, KLF9*), granule cell maturation genes (*TUBB4A, GRM4*), and cell cycle transcripts (*MCM2, PCNA, CCNB1, TOP2A*), which were quantified in the EGL. In the IGL, H-scores were computed for *MCM2, CCNB1,* and *TOP2A*; *PCNA* was excluded from IGL quantification because the signal was consistently below the detection threshold in postmigratory granule cells. Monotonic trends in H-scores across gestational age were assessed using Spearman’s rank correlation for each gene and cell type. Plots display the mean H-score with standard error of the mean at each gestational age, with individual ROI-level data points overlaid. Data are shown descriptively for validation. For PHH3 immunofluorescence in PCs, nuclear puncta and clusters were detected via QuPath’s intensity-threshold subcellular detection. The nuclear cluster fraction was defined as the number of nuclear PHH3 clusters divided by total nuclear PHH3 puncta per cell and averaged at the subject level. The nuclear-to-cytoplasmic ratio of PHH3 puncta was also computed per cell and averaged to the subject level. Pearson correlation was used for PHH3 nuclear cluster fraction and nuclear-to-cytoplasmic ratio, given their continuous nature and approximately linear relationships.

For both VGLUT1 and VGLUT2, positive-area fractions were averaged across ROIs to yield one value per subject, avoiding pseudoreplication from nonindependent ROIs. Pearson correlation coefficients were calculated to describe the relationship between positive-area fraction and gestational age at birth (n = 8). Preterm-versus-term comparisons were not performed because only one term subject was available. VGLUT1 and VGLUT2 were quantified using the same detection and normalization approach but analyzed separately for each marker. Given the limited sample size, these analyses were interpreted descriptively. Measurements were exported from QuPath (v0.6.0) and analyzed in R (v4.5; lme4, lmerTest), with figures generated using ggplot2.

### Intra-subject transcriptomic–neuropathological correlation

Five subjects in the postmortem cohort were profiled on the same tissue sections by both spatial transcriptomics and deep learning–based quantitative neuropathology (Preterm 1, Preterm 2, Preterm 52, Term 1, and Term 3), enabling direct comparison of the two modalities within the same subject. For each subject, the activity of each Cerebellar Maturation-Associated Pattern (CMAP) was reduced to a single value: the mean CoGAPS spot weight across the spots annotated to the layer corresponding to that pattern (Purkinje-cell-layer spots for CMAP_PCL; internal granule layer spots for CMAP_IGL and CMAP_IGL_IMM). Associations between transcriptomic pattern activity and the corresponding neuropathological measure were quantified by Spearman rank correlation, with leave-one-subject-out re-estimation to assess stability; two-sided exact-permutation *P* values (all 120 permutations of five observations) are reported for reference (table S20).

**Fig. S1.**
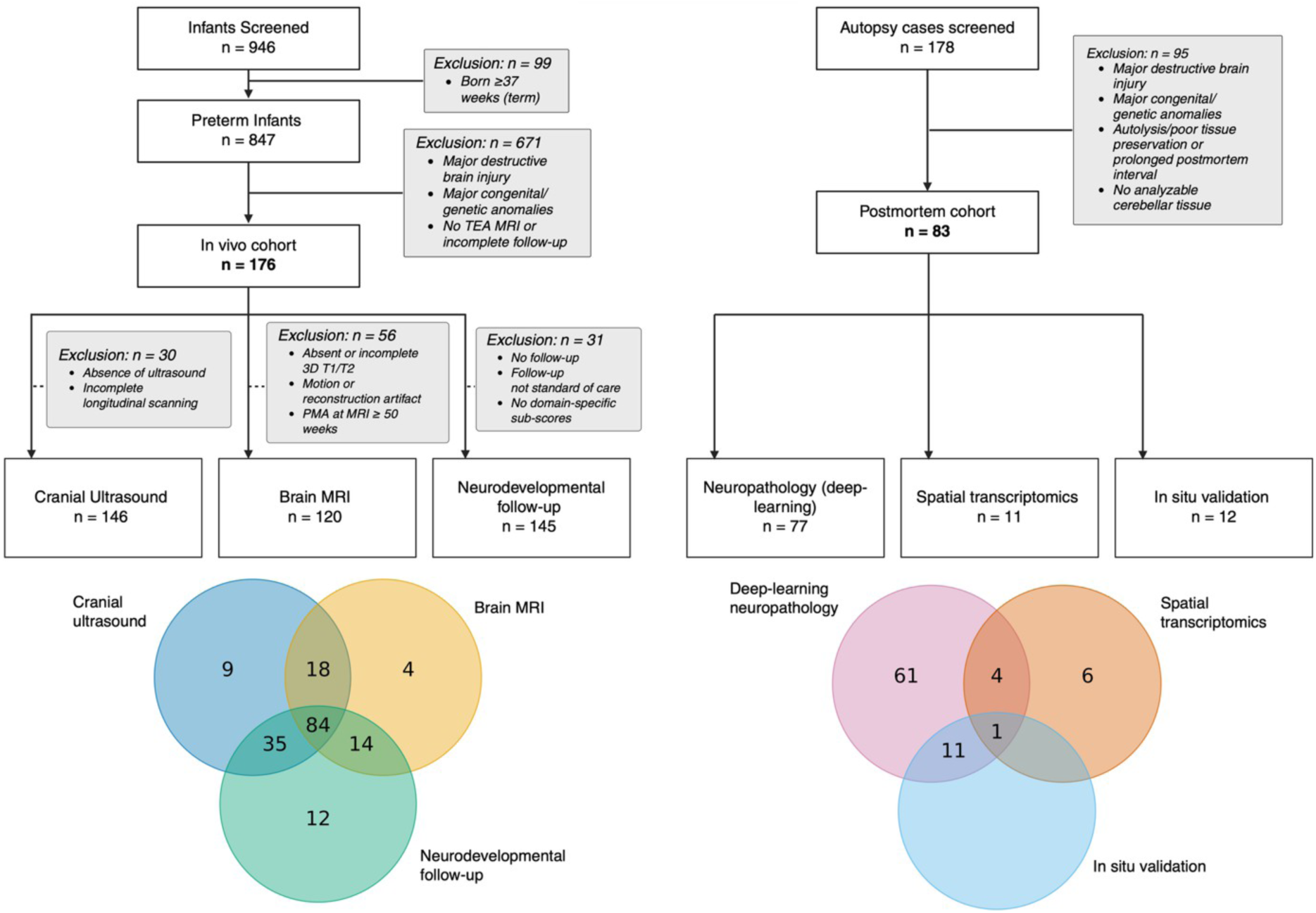
Cohort selection and subject overlap across in vivo and postmortem analyses. Left, in vivo cohort selection. Of 946 infants screened, 99 born at ≥37 weeks were excluded; of the remaining 847 preterm infants, 671 were excluded for major destructive brain injury, major congenital or genetic anomalies, absence of term-equivalent age (TEA) MRI, or incomplete follow-up, yielding the in vivo cohort (*n* = 176). Three analytic cohorts were derived: cranial ultrasound (*n* = 146; EP, 80; VP, 45; LP, 21), brain MRI (*n* = 120; EP, 65; VP, 37; LP, 18), and neurodevelopmental follow-up (*n* = 145; EP, 94; VP, 51). Modality-specific exclusions and reasons are indicated in the flow diagram. Right, postmortem cohort selection. Of 178 autopsy cases screened, 95 were excluded for major destructive brain injury, major congenital or genetic anomalies, autolysis or poor tissue preservation, prolonged postmortem interval, or absence of analyzable cerebellar tissue, yielding the postmortem cohort (*n* = 83). Analytic cohorts comprised deep-learning neuropathology (*n* = 77), spatial transcriptomics (*n* = 11), and in situ validation (*n* = 12; immunofluorescence, *n* = 9; RNAscope, *n* = 4; one subject assessed by both). Bottom, Venn diagrams show subject overlap across analyses within the in vivo (left) and postmortem (right) cohorts; numbers indicate unique or shared subjects. EP, extremely preterm; VP, very preterm; LP, late preterm; MRI, magnetic resonance imaging.

**Fig. S2.**
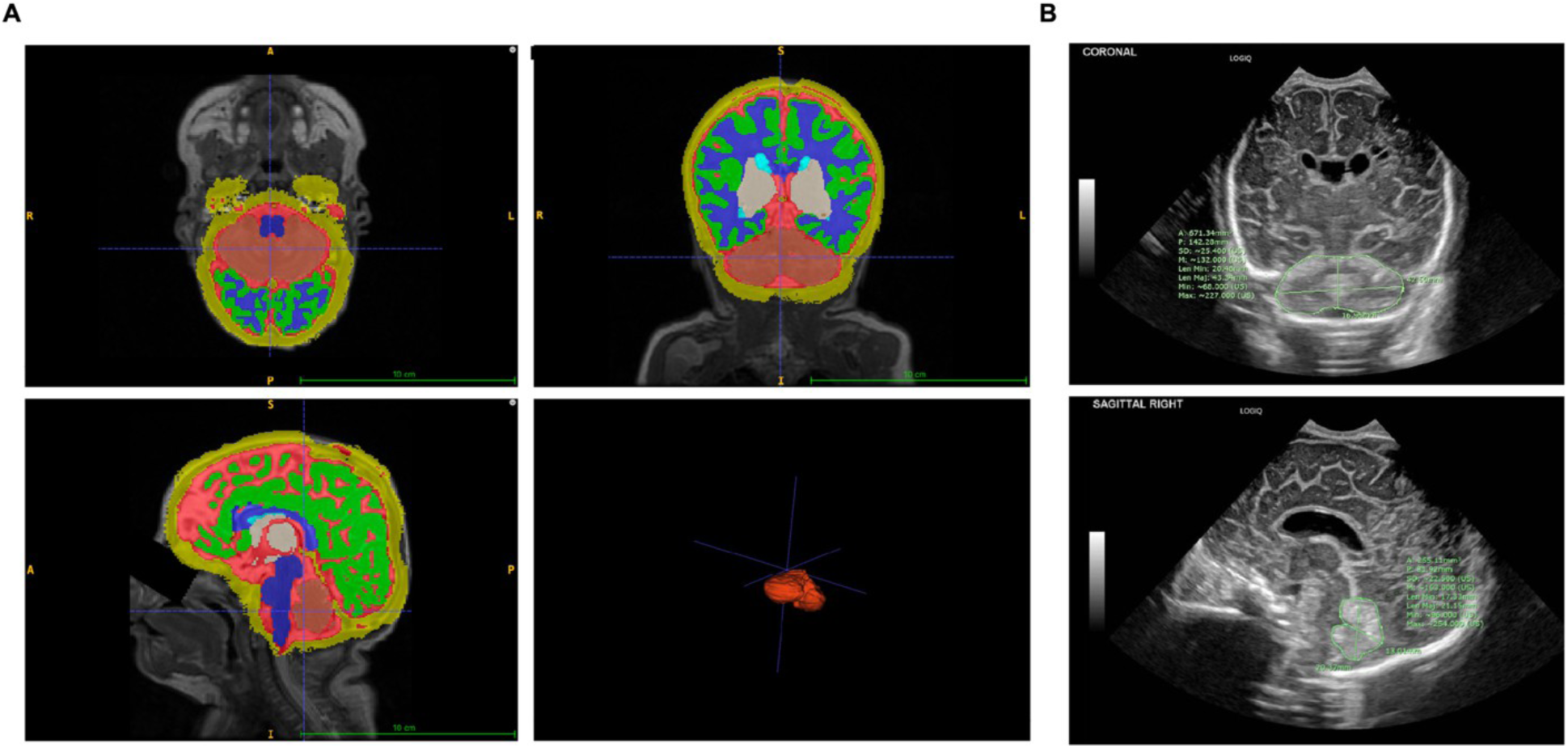
Representative MRI cerebellar segmentation and cranial ultrasound morphometry. **(A)** Representative brain MRI segmentation showing segmented brain structures with the cerebellum highlighted in red in axial (top left), coronal (top right), and sagittal (bottom left) views, with the corresponding three-dimensional cerebellar reconstruction (bottom right). Segmentation labels are overlaid on the anatomical images. Scale bar, 10 cm. **(B)** Representative cranial ultrasound images showing coronal (top) and sagittal (bottom) measurement planes used for cerebellar morphometry. Coronal measurements include transcerebellar diameter and cross-sectional area; sagittal measurements include anteroposterior diameter, height, and cross-sectional area. Measurement axes and contours are shown in green.

**Fig. S3.**
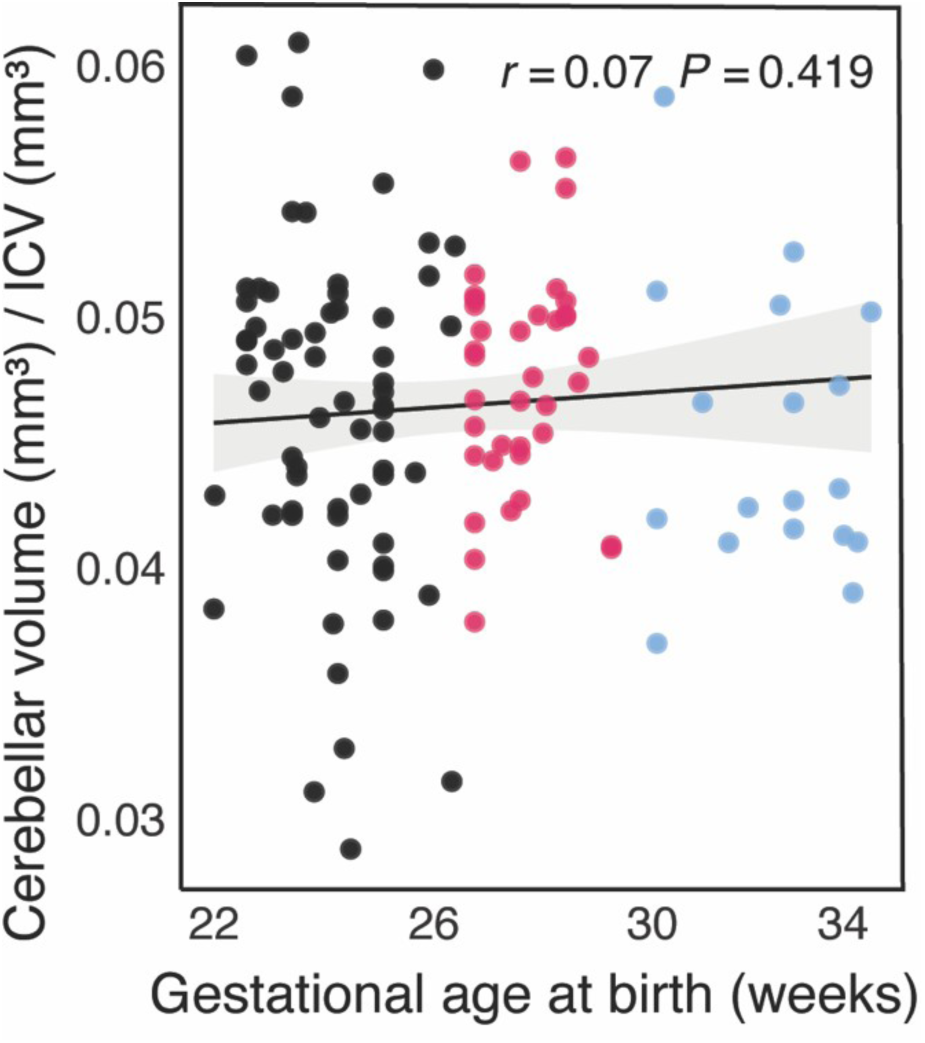
Cerebellar volume normalized to intracranial volume. Cerebellar volume at term-equivalent age normalized to intracranial volume (ICV) and plotted against gestational age at birth in extremely preterm (EP, black), very preterm (VP, magenta), and late preterm (LP, light blue) infants (*n* = 120; EP = 65, VP = 37, LP = 18). Line, ordinary least-squares fit; shaded band, 95% CI; partial *r*, adjusted for post-menstrual age at MRI. The cerebellum-to-ICV ratio was not associated with gestational age at birth (partial *r* = 0.07; *P* = 0.419), indicating proportional scaling of cerebellar and intracranial volumes.

**Fig. S4.**
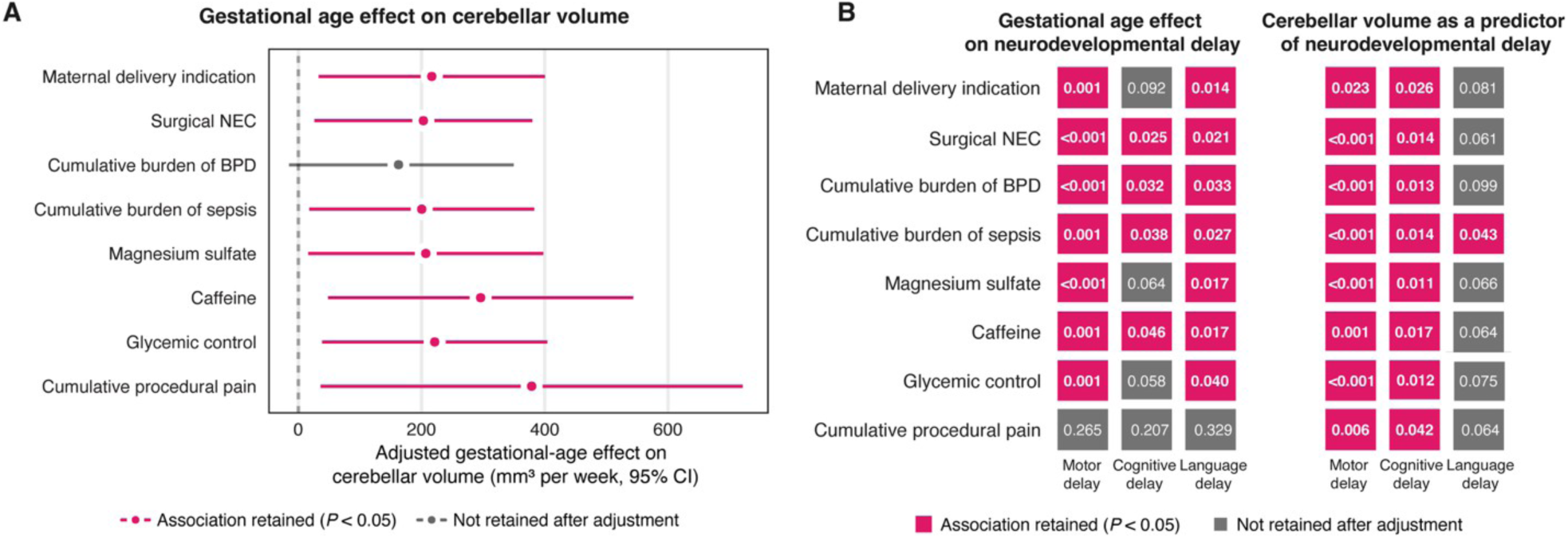
Persistence of gestational age and cerebellar volume associations after adjustment for individual perinatal exposures. **(A)** Association between gestational age at birth (GA) and term-equivalent age (TEA) cerebellar volume after adjustment for individual perinatal exposures. Each row shows the GA coefficient and 95% CI from a separate model including post-menstrual age at MRI and the indicated exposure as covariates. Points, GA coefficients; horizontal lines, 95% CI; dashed line, no association. Magenta, association retained after adjustment (*P* < 0.05); gray, not retained. Model size ranged from *n* = 115 to *n* = 120. **(B)** Persistence of the associations of GA (left) and TEA cerebellar volume (right) with motor, cognitive, and language delay after adjustment for each indicated perinatal exposure. Each cell shows the *P* value from a separate adjusted model. Magenta, association retained (*P* < 0.05); gray, not retained. Model size ranged from *n* = 140 to *n* = 143 for GA models and from *n* = 90 to *n* = 97 for cerebellar-volume models. Perinatal exposures included maternal delivery indication, surgical necrotizing enterocolitis (NEC), cumulative burden of bronchopulmonary dysplasia (BPD), cumulative burden of sepsis, magnesium sulfate, caffeine, glycemic control, and cumulative procedural pain. Each exposure was entered separately as an additive covariate; no interaction terms were included.

**Fig. S5.**
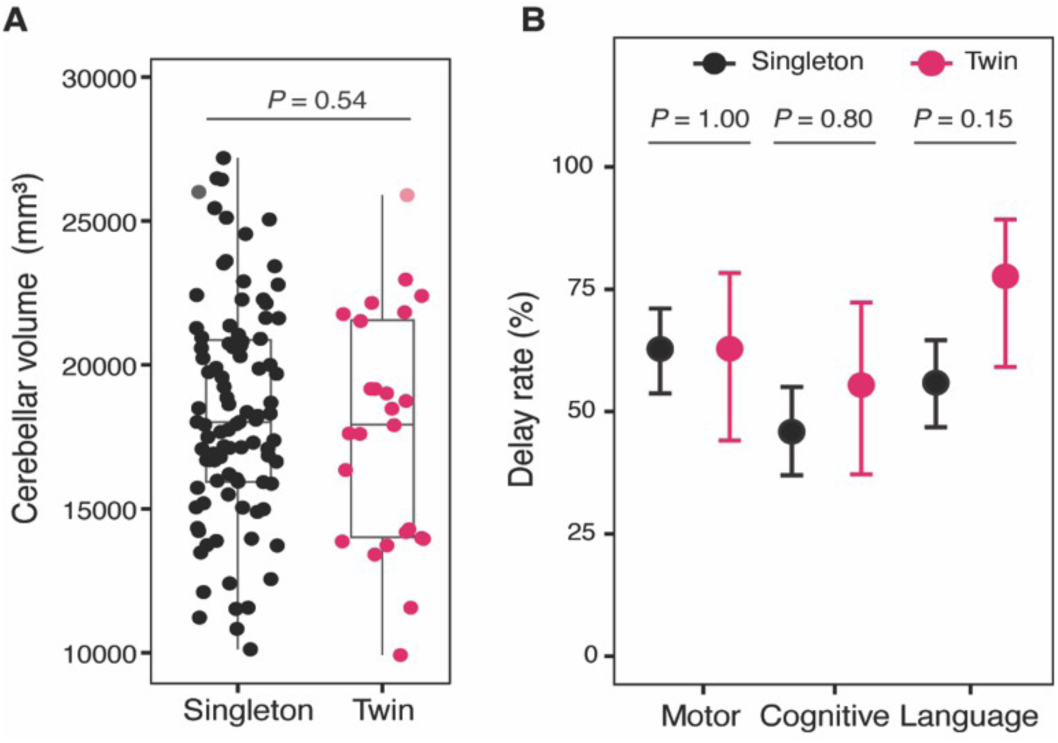
Twin status does not account for cerebellar volume or neurodevelopmental delay. **(A)** Term-equivalent age (TEA) cerebellar volume in singletons (*n* = 95) and twins (*n* = 25) in the volumetric MRI cohort. Two-sided Wilcoxon rank-sum test; box, median and IQR; whiskers, 1.5× IQR; points, individual infants. Gestational age at birth did not differ between groups (*P* = 0.82), and inclusion of twin status did not alter the gestational age–volume association (twin-status coefficient, +136 mm³; *P* = 0.86). **(B)** Motor, cognitive, and language delay rates in extremely and very preterm infants by twin status (singletons, *n* = 118; twins, *n* = 27). Points, proportion with delay; error bars, 95% Wilson CI. *P* values, two-sided Fisher’s exact tests, Holm-adjusted across the three domains. No delay domain differed by twin status after adjustment. Singletons, black; twins, magenta.

**Fig. S6.**
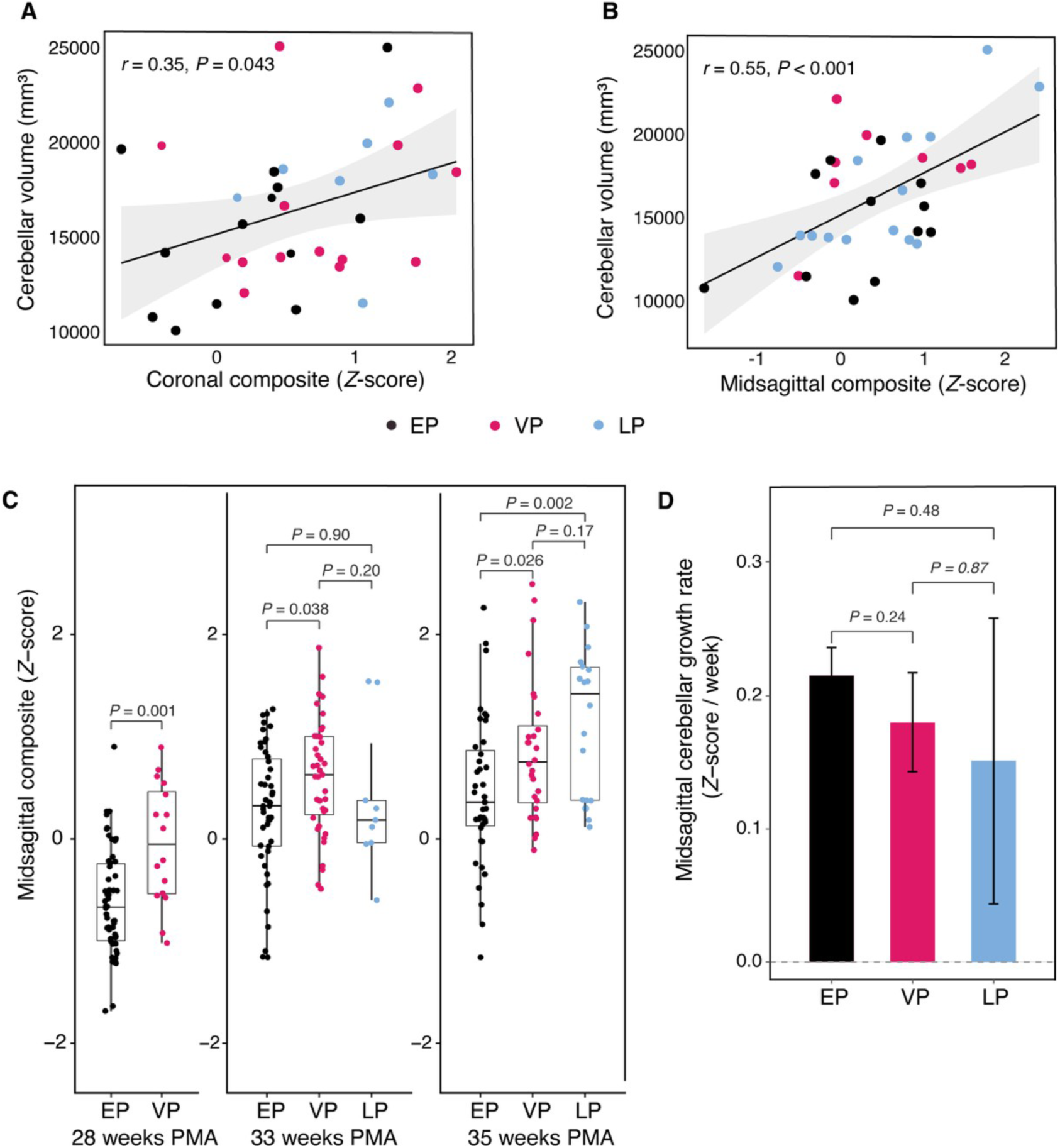
Cross-modal correlation of ultrasound and MRI measures and midsagittal cerebellar growth across preterm groups. Extremely preterm (EP, black), very preterm (VP, magenta), and late preterm (LP, light blue) infants are shown throughout. **(A and B)** Association between bedside ultrasound measures and segmented MRI cerebellar volume in infants with ultrasound performed within 2 weeks of MRI (*n* = 35): coronal composite (A) and midsagittal composite (B), with ultrasound composites expressed as *Z*-scores; two-sided Pearson correlation. **(C)** Midsagittal composite cerebellar size (*Z*-score) at 28, 33, and 35 weeks post-menstrual age (PMA); 28 weeks includes EP and VP only. Box plots, median and IQR with individual subjects overlaid; pairwise two-sided Wilcoxon rank-sum tests. **(D)** Midsagittal cerebellar growth rate (*Z*-score per week) by group. Bars, mean; error bars, 95% CI; pairwise slope contrasts from mixed-effects models, Tukey-adjusted. (C and D), *n* = 146 (EP = 80, VP = 45, LP = 21).

**Fig. S7.**
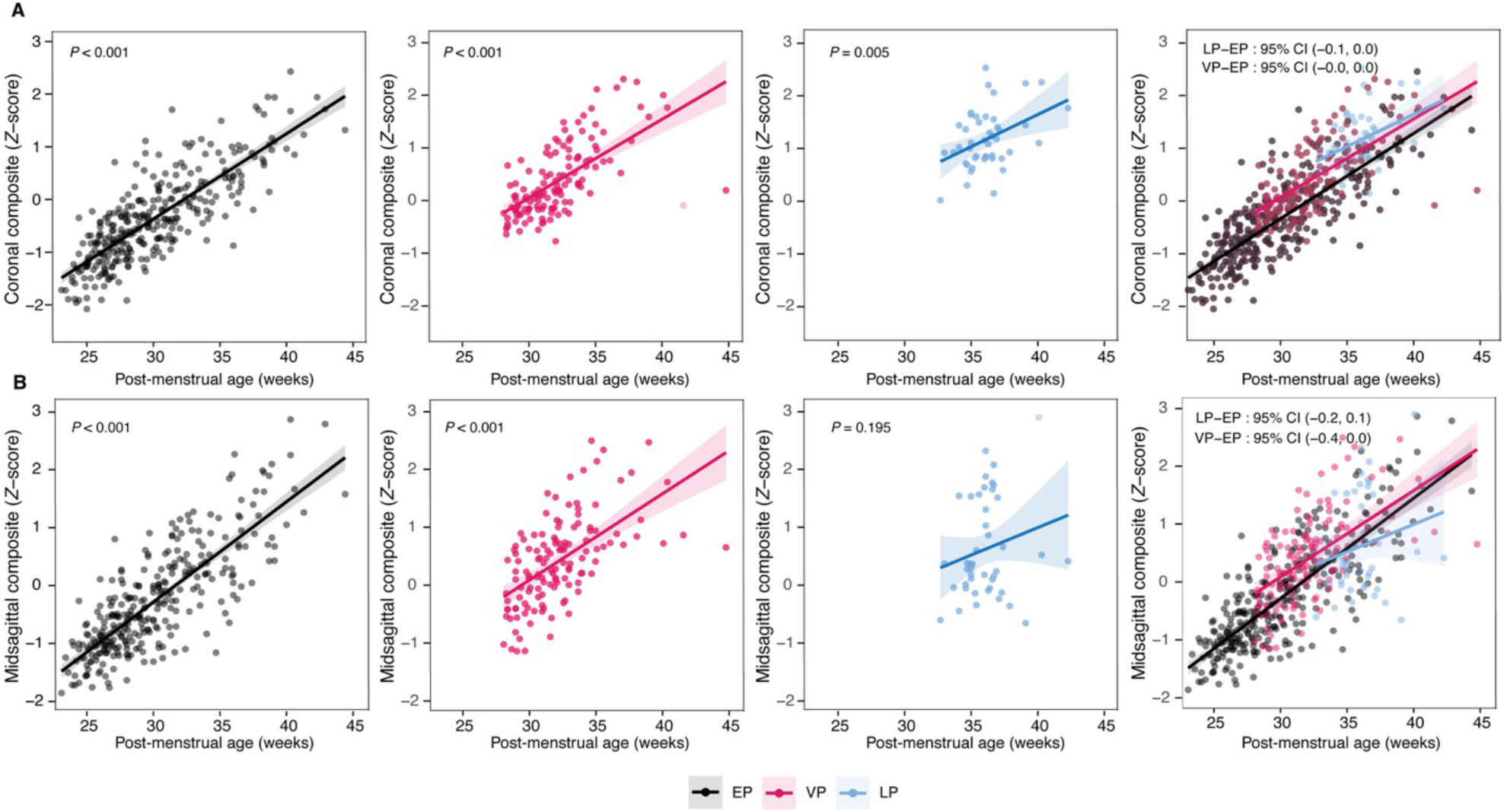
Longitudinal cerebellar growth trajectories across gestational age groups. (**A and B**) Coronal composite cerebellar size (A) and midsagittal composite cerebellar size (B), each expressed as a *Z*-score and plotted against post-menstrual age (PMA) for extremely preterm (EP, black), very preterm (VP, magenta), and late preterm (LP, light blue) infants (*n* = 146; EP = 80, VP = 45, LP = 21). The first three columns show EP, VP, and LP trajectories, respectively; the fourth column shows all groups overlaid. Each point represents one ultrasound measurement. Solid lines, linear mixed-effects model fits (PMA × group, per-infant random effect); shaded bands, 95% CI. The overlaid panels report pairwise slope differences (LP−EP and VP−EP) with Tukey-adjusted 95% CI. *P* values in the group-specific panels test the within-group PMA slope. Growth rates did not differ detectably between groups in either plane.

**Fig. S8.**
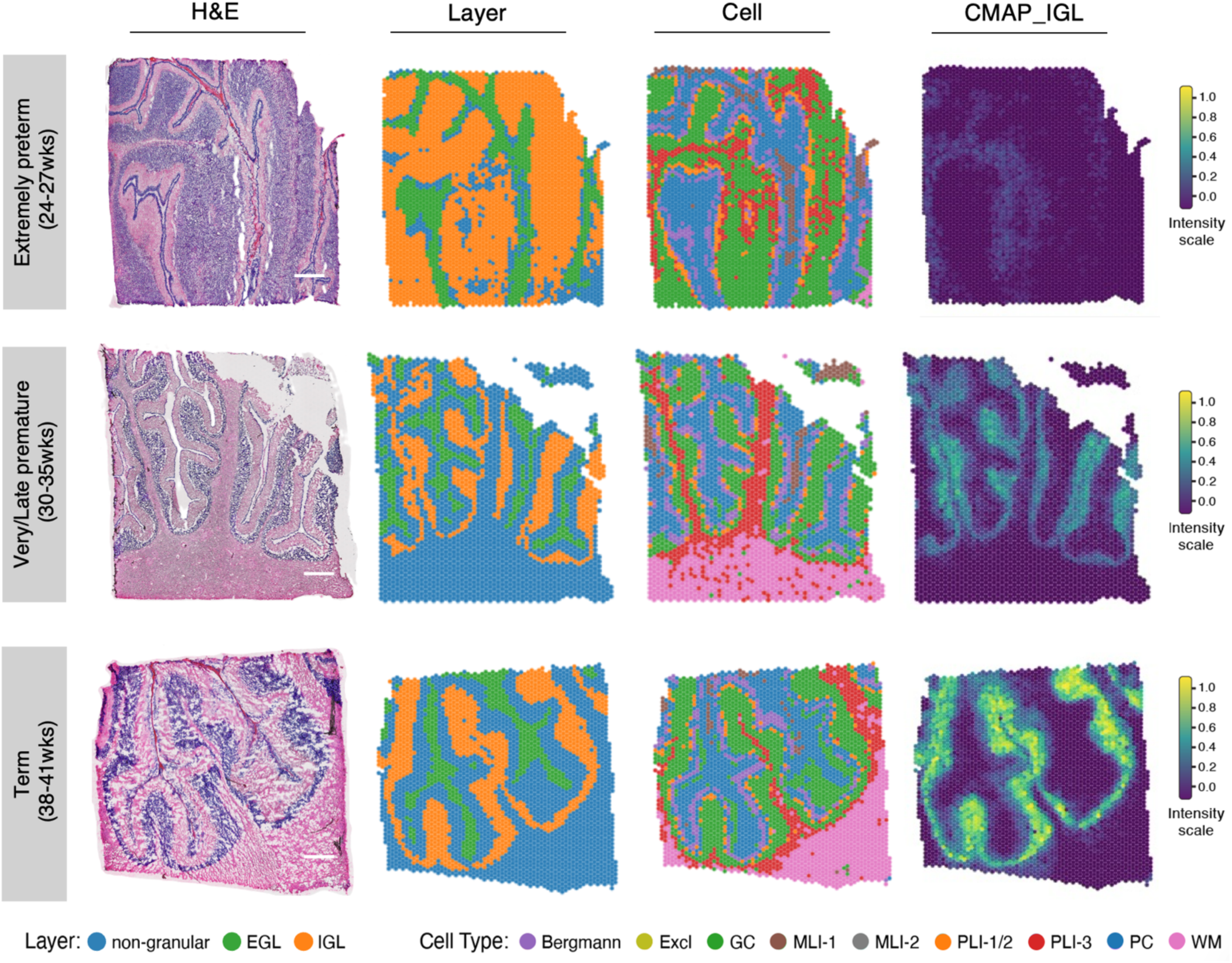
Spatial transcriptomic architecture of the human cerebellar cortex across the gestational spectrum. Representative postmortem cerebellar sections profiled by Visium spatial transcriptomics. Rows (top to bottom): extremely preterm (24–27 weeks gestational age at birth, GA), very/late preterm (30–35 weeks GA), and term (38–40 weeks GA) subjects. Columns (left to right): hematoxylin and eosin (H&E) staining of the profiled section; spatial annotation of cortical layers (Layer); spatial annotation of cell types (Cell); and spatial activity of CMAP_IGL, shown as an example of a cerebellar maturation-associated pattern (CMAP) derived by Bayesian non-negative matrix factorization (CoGAPS). Scale bars, 1 mm. Each spot represents one Visium capture location. Layer annotations: non-granular, external granule layer (EGL), and internal granule layer (IGL). Cell-type annotations: Bergmann glia, excluded tissue (Excl), granule cells (GC), molecular layer interneurons (MLI-1, MLI-2), Purkinje layer interneurons (PLI-1/2, PLI-3), Purkinje cells (PC), and white matter (WM). CMAP activity is shown on a normalized scale from 0.0 to 1.0 (color bar). Analyses were performed on *n* = 11 subjects with two sections per subject (22 sections total; tables S6 and S7).

**Fig. S9.**
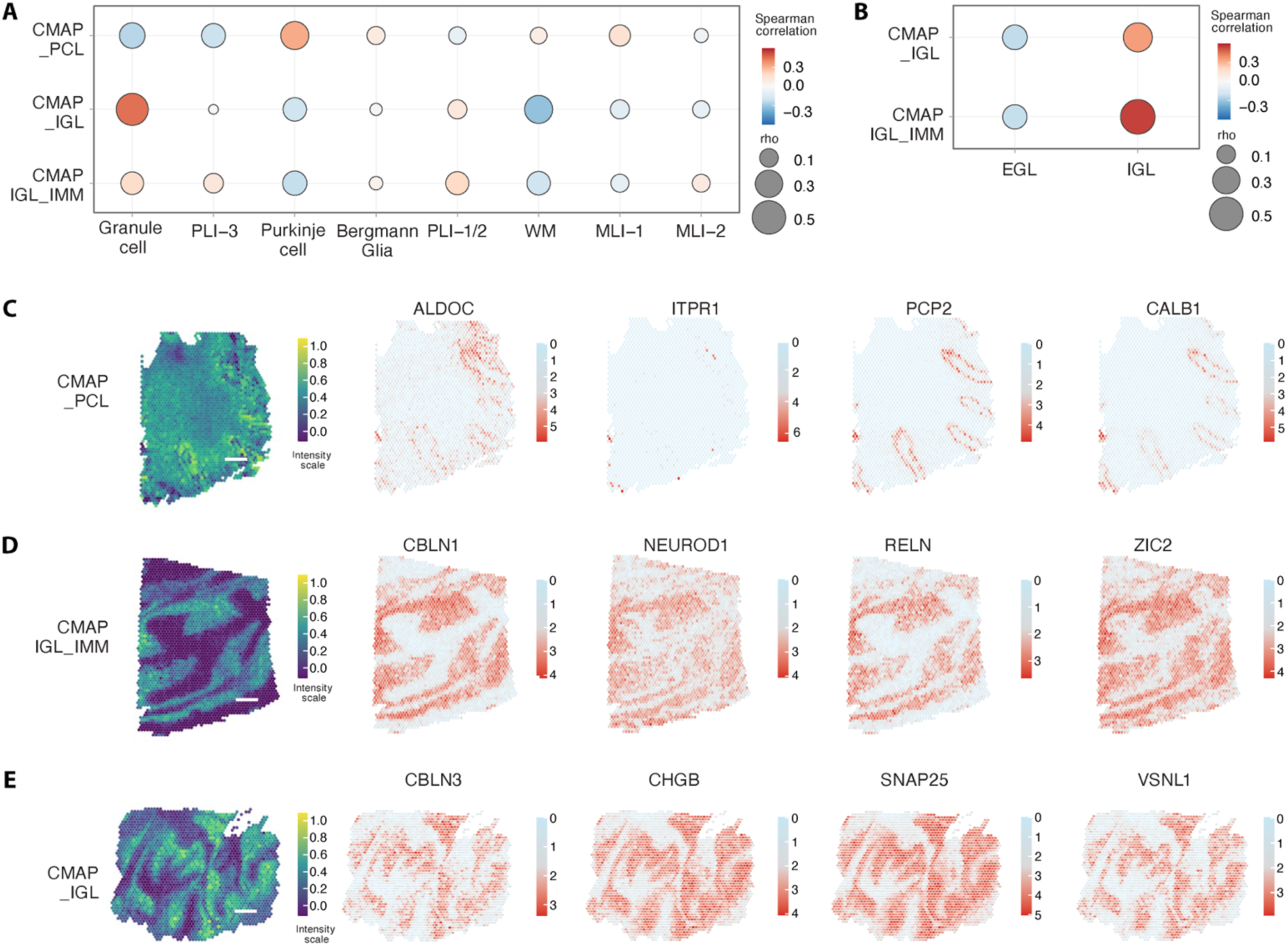
Cerebellar maturation-associated patterns localize to distinct cell types and laminar compartments. **(A)** Spearman correlations between cerebellar maturation-associated pattern (CMAP) activity and cell-type annotations across the spatial transcriptomic cohort. Dot color, Spearman rho; dot size, absolute correlation magnitude (|rho|). MLI, molecular layer interneuron; PLI, Purkinje layer interneuron; WM, white matter. **(B)** Spearman correlations between CMAP_IGL and CMAP_IGL_IMM activity and laminar annotations of the external granule layer (EGL) and internal granule layer (IGL), displayed as in (A). **(C to E)** Representative spatial activity maps of CMAP_PCL (C), CMAP_IGL_IMM (D), and CMAP_IGL (E), shown alongside spot-level expression of canonical marker genes: *ALDOC*, *ITPR1*, *PCP2*, and *CALB1* (C); *CBLN1*, *NEUROD1*, *RELN*, and *ZIC2* (D); and *CBLN3*, *CHGB*, *SNAP25*, and *VSNL1* (E). Scale bars, 1 mm. Color scales indicate normalized CMAP activity (left) and spot-level gene expression (right). Representative sections are from subjects born at 24.4 (C), 26 (D), and 39 (E) weeks of gestation. (A to E) *n* = 11 subjects, two sections per subject (22 sections total).

**Fig. S10.**
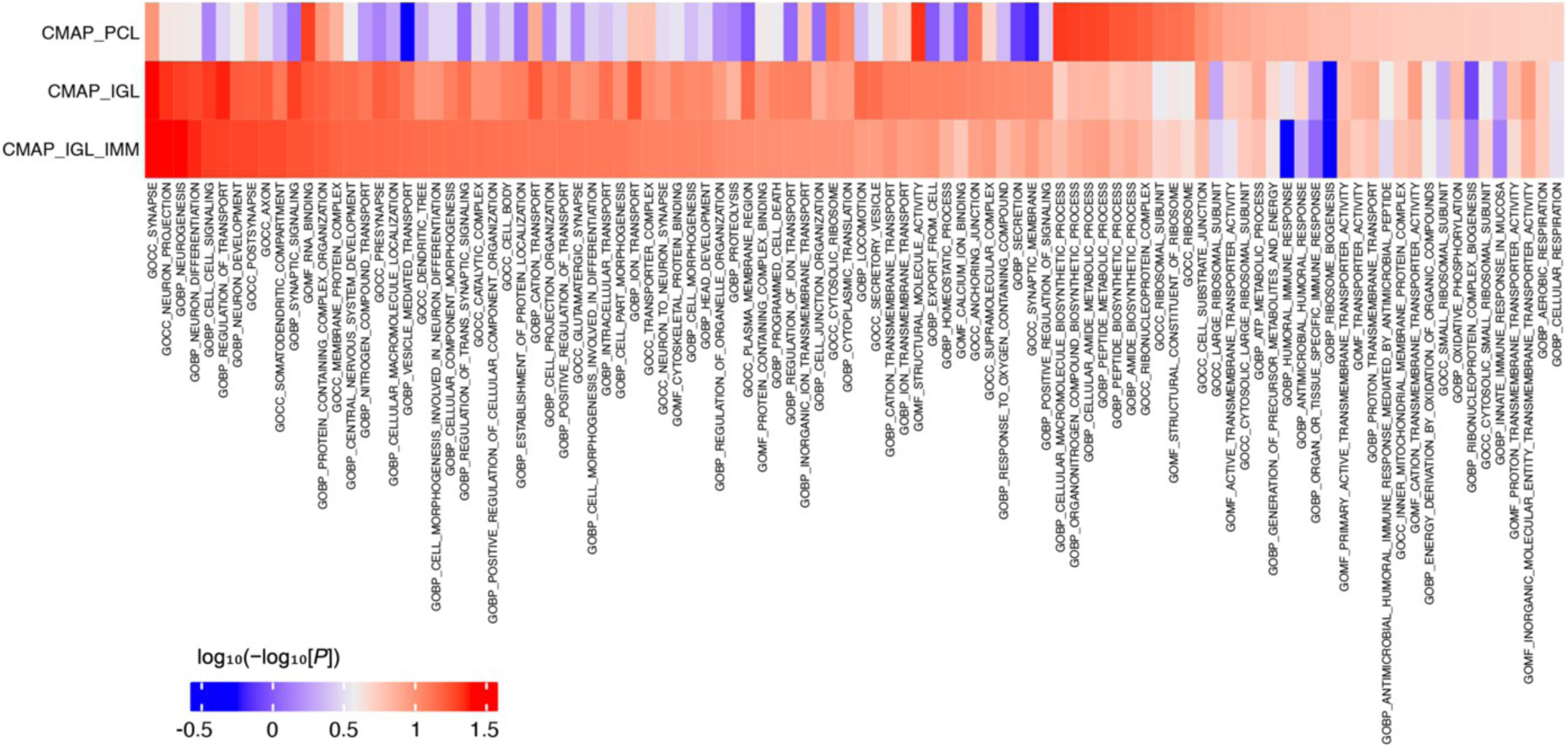
Functional enrichment of cerebellar maturation-associated patterns. Heat map of Gene Ontology (GO) enrichment across gene sets associated with CMAP_PCL, CMAP_IGL, and CMAP_IGL_IMM. Columns represent GO terms enriched in at least one CMAP-associated gene set, spanning biological process (GOBP), cellular component (GOCC), and molecular function (GOMF) categories. Enrichment profiles differed across CMAPs, reflecting distinct developmental, synaptic, metabolic, and cellular programs. Color encodes the transformed nominal *P* value from fast gene set enrichment analysis (fGSEA), log₁₀(−log₁₀[nominal *P*]). CMAPs were derived from 11 subjects with two sections per subject (22 sections total).

**Fig. S11.**
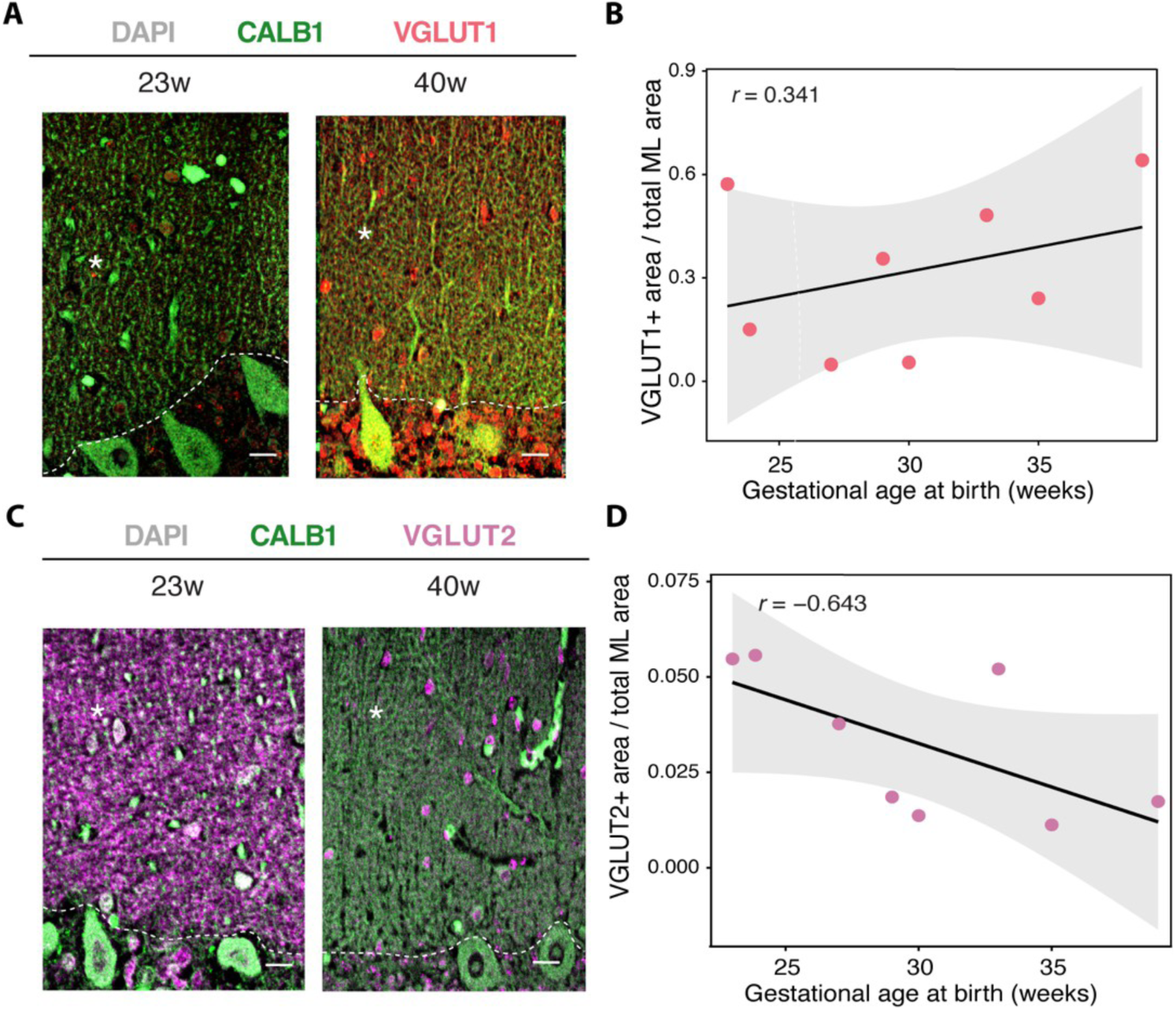
VGLUT1 and VGLUT2 immunoreactivity in the cerebellar molecular layer across gestational age at birth. **(A)** Representative immunofluorescence images of the cerebellar molecular layer (ML) from subjects born at 23 and 40 weeks’ gestation, stained for calbindin (CALB1, green), VGLUT1 (red), and DAPI (gray). Asterisks, molecular layer; dashed lines, Purkinje cell layer–molecular layer border. Scale bar, 10 µm. **(B)** VGLUT1-positive area as a fraction of total ML area plotted against gestational age at birth (Pearson *r* = 0.341). **(C)** Representative immunofluorescence images stained for CALB1 (green), VGLUT2 (magenta), and DAPI (gray), as in (A). Scale bar, 10 µm. **(D)** VGLUT2-positive area as a fraction of total ML area plotted against gestational age at birth (Pearson *r* = −0.643). (B and D) Each point represents one subject and the mean of five regions of interest (ROIs); line, linear fit; shaded band, 95% CI. Eight subjects were analyzed per marker (nine unique subjects; the term subject differed between markers).

**Fig. S12.**
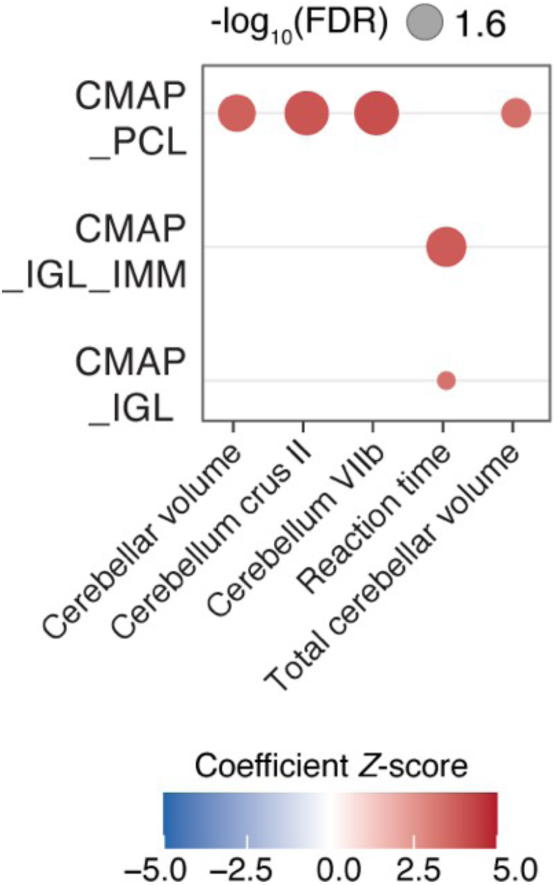
LDSC-based partitioned heritability enrichment of CMAP gene sets across neuroimaging and cognitive traits. Partitioned heritability enrichment of CMAP_PCL, CMAP_IGL_IMM, and CMAP_IGL cerebellar maturation-associated pattern (CMAP) gene sets across cerebellar imaging and cognitive traits was assessed by linkage disequilibrium score regression (LDSC) using published genome-wide association study summary statistics (table S8). Rows indicate CMAP gene sets and columns indicate traits. Only associations passing a false discovery rate (FDR) of <0.05 are shown. Dot size, −log₁₀ FDR; color, regression coefficient *Z*-score. CMAP gene sets were derived from the spatial transcriptomic cohort (*n* = 11 subjects, two sections per subject).

**Fig. S13.**
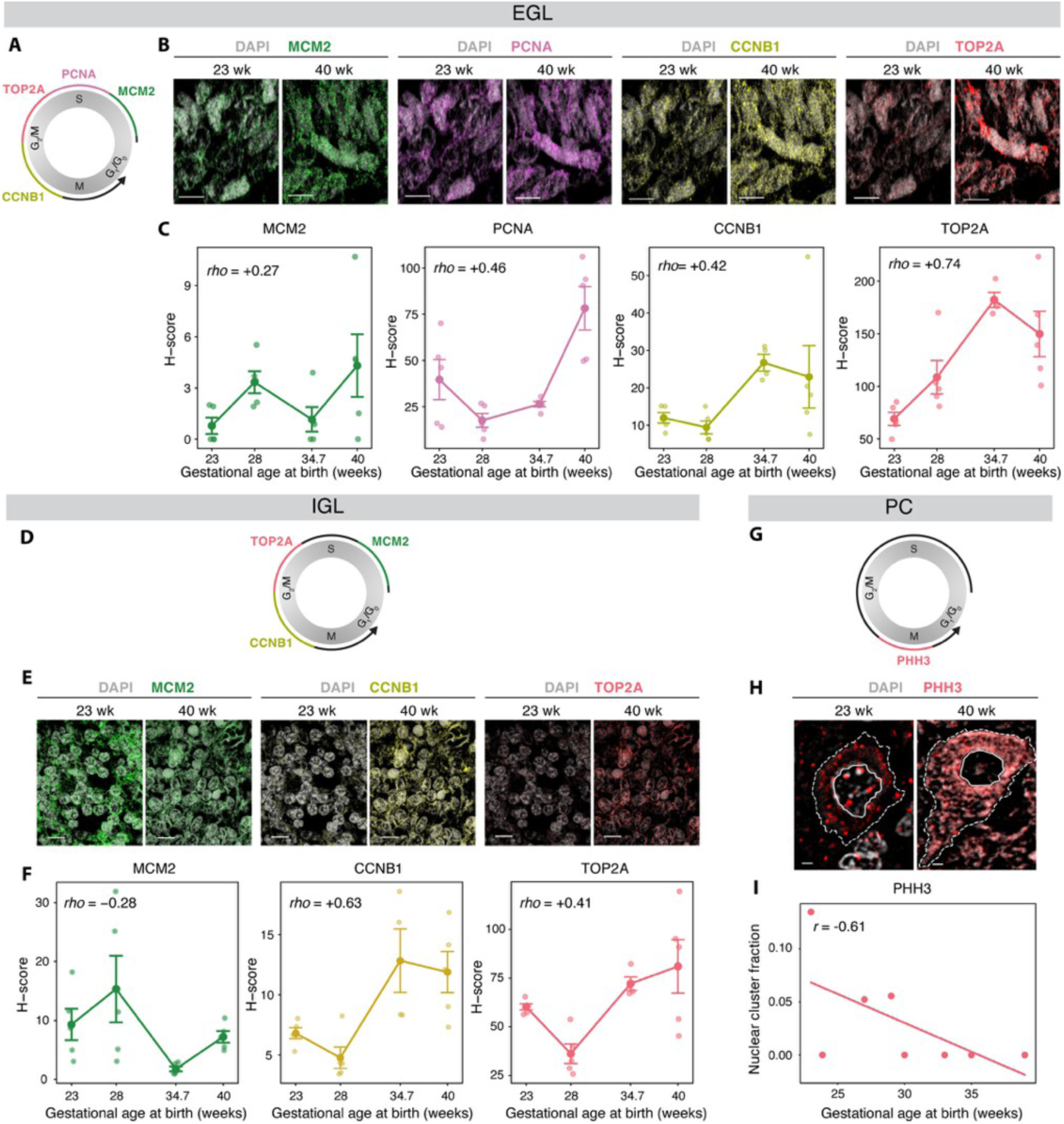
Orthogonal validation of phase-associated molecular states across cerebellar lineages and gestational ages. **(A)** Schematic of *MCM2*, *PCNA*, *CCNB1*, and *TOP2A* transcript positions along the inferred cell cycle continuum in the external granule layer (EGL). **(B)** Representative RNAscope images of the EGL in subjects born at 23 and 40 weeks of gestation, showing *MCM2* (green), *PCNA* (purple), *CCNB1* (yellow), *TOP2A* (magenta), and DAPI (gray). Scale bar, 10 µm. **(C)** H-score quantification of EGL marker expression across gestational age at birth (*n* = 4 subjects; 4–5 regions of interest [ROIs] per subject). Small points, individual ROIs; large points, subject means; lines connect subject means; error bars, SEM across ROIs. Spearman rho between subject-mean H-score and gestational age at birth is shown in each panel. **(D)** Schematic of *MCM2*, *TOP2A*, and *CCNB1* transcript positions along the inferred cell cycle continuum in the internal granule layer (IGL). **(E)** Representative RNAscope images of the IGL for the markers in (D), with colors and staining as in (B). Scale bar, 10 µm. **(F)** H-score quantification of IGL marker expression across gestational age at birth, as in (C). **(G)** Schematic of phosphorylated histone H3 (PHH3) localization along the phase-associated continuum in Purkinje cells (PCs). **(H)** Representative immunofluorescence images of PCs showing PHH3 (magenta) with DAPI (gray); dashed outlines, annotated cells. Scale bar, 2 µm. **(I)** PHH3 nuclear cluster fraction in PCs plotted against gestational age at birth (*n* = 8 subjects; 5 ROIs per subject, 5–6 PCs per ROI). The nuclear cluster fraction was calculated as the number of nuclear PHH3 clusters divided by total nuclear PHH3 puncta per cell and averaged per subject. Two-sided Pearson correlation.

**Fig. S14.**
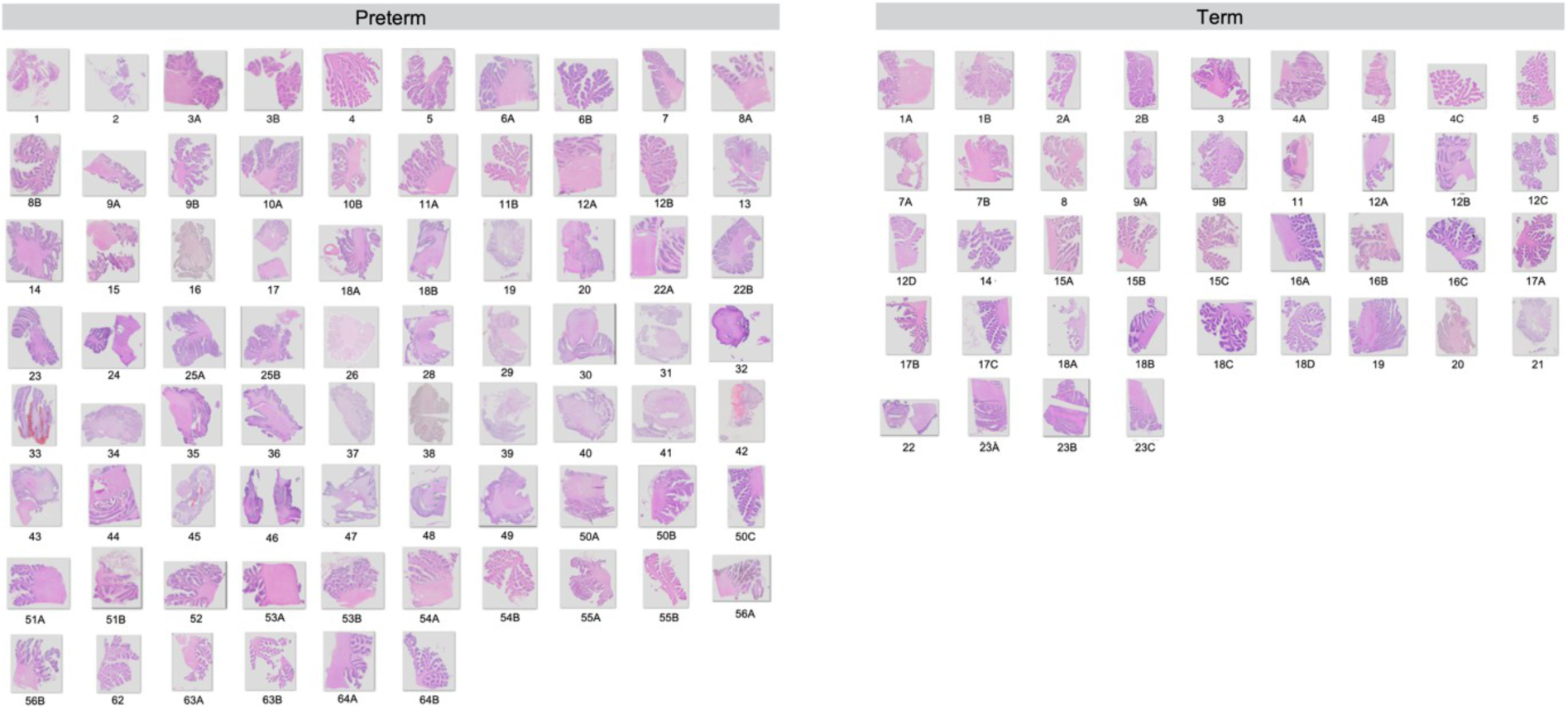
Whole-slide hematoxylin and eosin images used for deep learning–based neuropathological analysis. Whole-slide hematoxylin and eosin (H&E) images of formalin-fixed, paraffin-embedded postmortem cerebellar sections from the deep-learning neuropathology cohort (*n* = 77; preterm, *n* = 57; term, *n* = 20). Five subjects also contributed to the spatial transcriptomic cohort (fig. S1). Each thumbnail represents one cerebellar section. Subject identifiers are numbered within group (tables S11 and S12); suffixes (A, B, C, etc.) denote multiple sections from the same subject. Thumbnails are individually scaled for layout and not shown at a common magnification; quantification was performed on calibrated whole-slide images at native resolution.

**Fig. S15.**
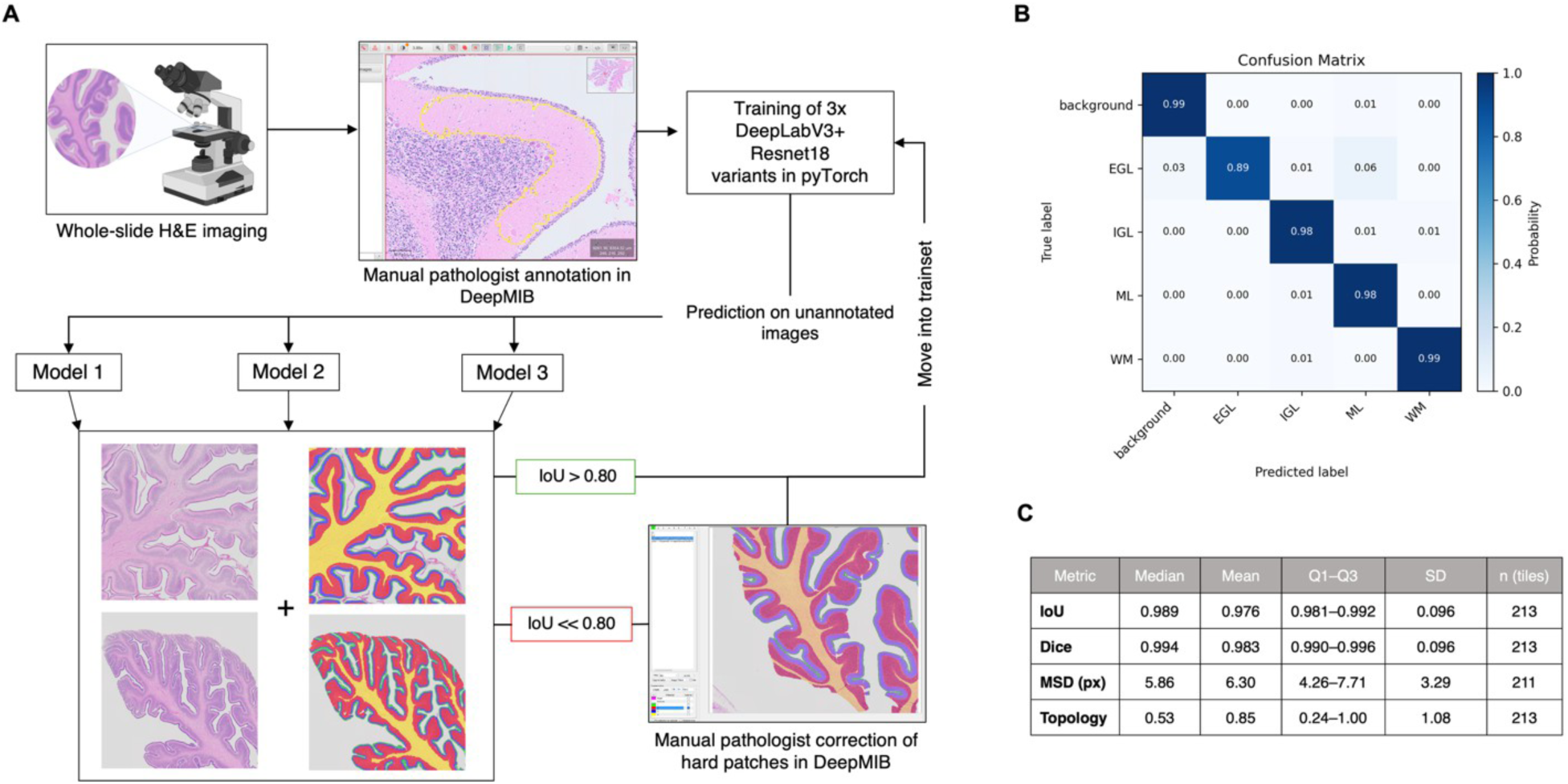
Deep-learning segmentation workflow and performance. **(A)** Pathologist-in-the-loop workflow for cerebellar layer segmentation from whole-slide hematoxylin and eosin (H&E) images. Manually annotated regions in DeepMIB were used to train three DeepLabV3+ ResNet18 models. Predictions were applied to previously unannotated images; regions with IoU > 0.80 were incorporated into the training set, whereas lower-IoU patches were manually reviewed and corrected by a pathologist before retraining. This iterative process was repeated until model performance stabilized. **(B)** Class-wise confusion matrix for segmentation of background, external granule layer (EGL), internal granule layer (IGL), molecular layer (ML), and white matter (WM) on a held-out validation set. Purkinje cells were quantified separately. **(C)** Segmentation performance on the held-out validation set, reported as intersection over union (IoU), Dice coefficient, mean symmetric surface distance (MSD; pixels), and topology score. Validation metrics were calculated from *n* = 213 image tiles (MSD, *n* = 211). Median IoU and Dice coefficients were 0.989 and 0.994, respectively.

**Fig. S16.**
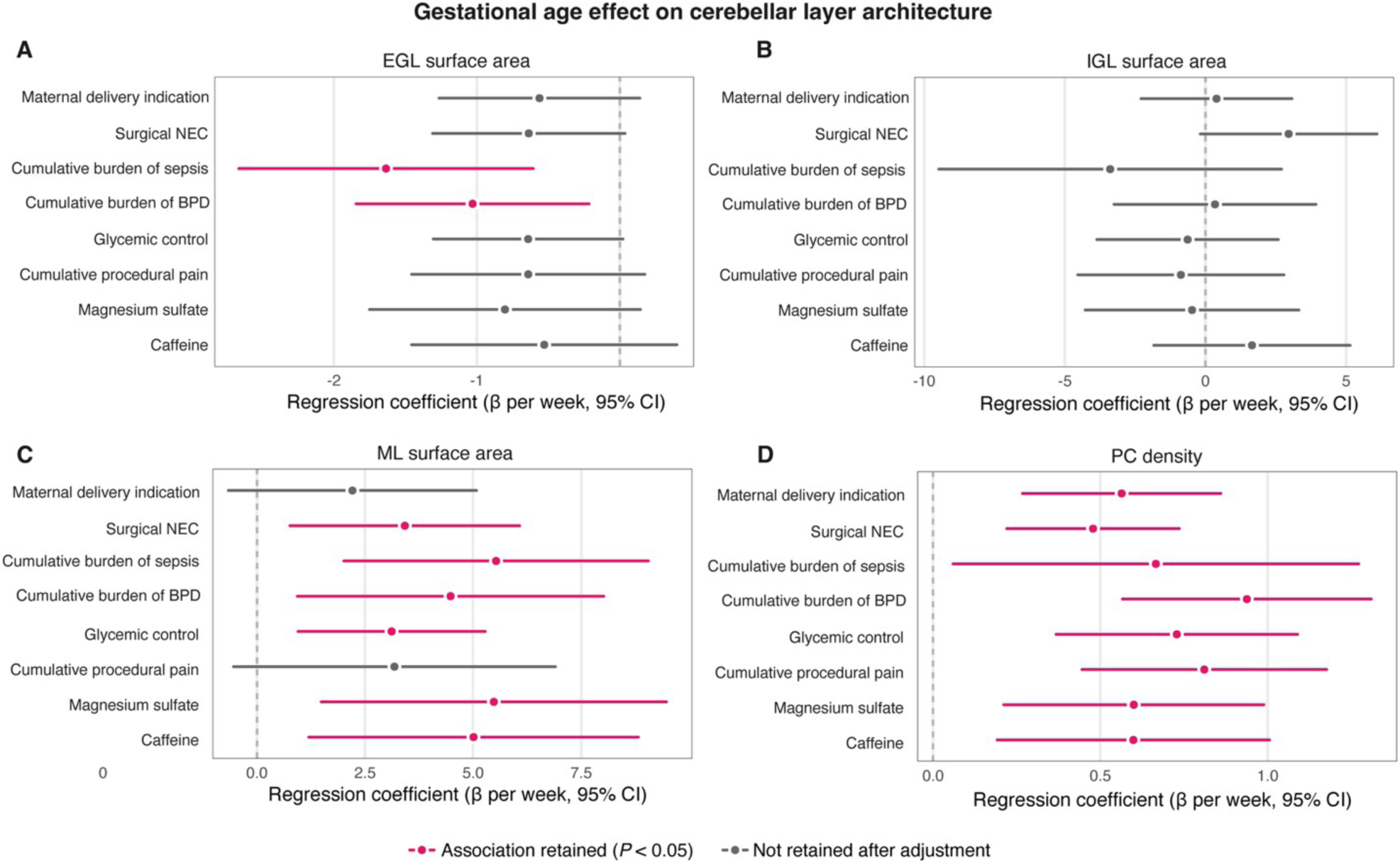
Persistence of gestational age associations with cerebellar layer architecture after adjustment for individual perinatal exposures. (**A to D**) Associations between gestational age at birth (GA) and cerebellar cytoarchitecture after adjustment for individual perinatal exposures: external granule layer (EGL) surface area (A), internal granule layer (IGL) surface area (B), and molecular layer (ML) surface area (C), each normalized to ML length (mm² per µm ML), and Purkinje cell (PC) density (D; PCs per mm ML), in the deep-learning–based neuropathology cohort. Each row shows the GA regression coefficient (β per week) and 95% CI from a separate model including post-menstrual age at death and the indicated exposure as covariates. Points, β; horizontal lines, 95% CI; dashed line, β = 0. Magenta, association retained after adjustment (*P* < 0.05); gray, not retained. Perinatal exposures included maternal delivery indication, surgical necrotizing enterocolitis (NEC), cumulative burden of sepsis, cumulative burden of bronchopulmonary dysplasia (BPD), glycemic control, cumulative procedural pain, magnesium sulfate, and caffeine. Each exposure was entered separately as an additive covariate; no interaction terms were included. Model size ranged from *n* = 21 (sepsis) to *n* = 71 (NEC) and was the same across all four panels for each exposure. Each panel has its own x-scale.

**Fig. S17.**
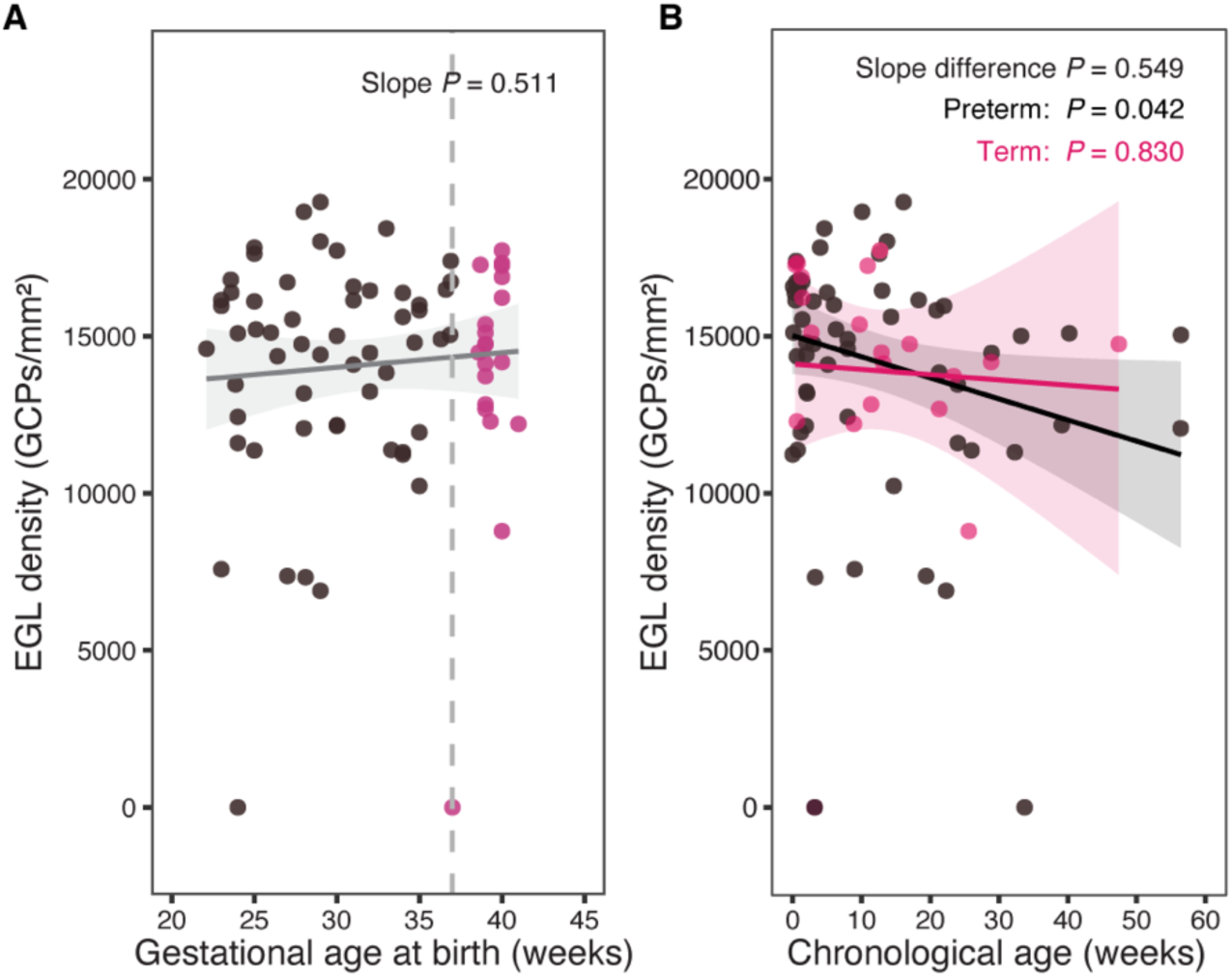
External granule layer density across gestational age at birth and chronological age. External granule layer (EGL) granule cell precursor (GCP) density (GCPs per mm²) in the neuropathology cohort (*n* = 77 subjects; preterm, *n* = 57, black; term, *n* = 20, magenta). Each point represents one subject. **(A)** EGL density plotted against gestational age at birth. Line, ordinary least-squares fit; shaded band, 95% CI; dashed line, 37 weeks (preterm–term boundary). EGL density was not associated with gestational age at birth (slope *P* = 0.511). **(B)** EGL density plotted against chronological age, with group-specific fits (preterm slope, −67.13 [95% CI, −130.31 to −3.95], *P* = 0.042; term slope, −16.80 [95% CI, −168.03 to 134.43], *P* = 0.830). Shaded bands, 95% CI. The preterm and term slopes did not differ (slope-difference *P* = 0.549).

**Fig. S18.**
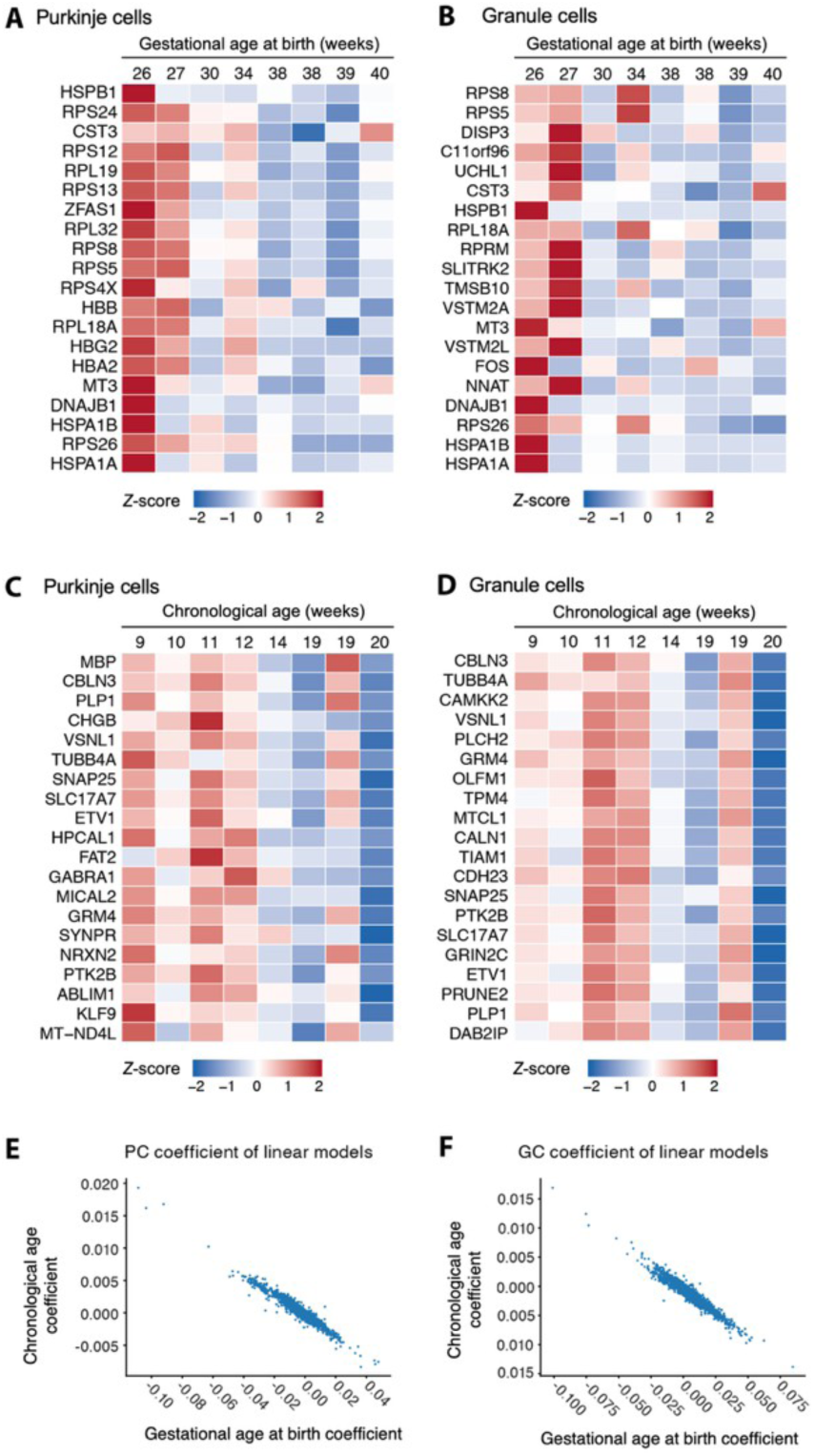
Gestational age– and chronological age-associated gene expression patterns are inversely related. (**A and B**) Heat maps of genes negatively associated with gestational age at birth (GA) in Purkinje cells (A) and granule cells (B), from spatial transcriptomic samples matched by post-menstrual age at death. Values are *Z*-scored mean expression per subject, ordered by GA. **(C and D)** Genes positively associated with GA (Fig. 4A and E) displayed across chronological age (CA) in Purkinje cells (C) and granule cells (D), illustrating the inverse relationship between GA– and CA-associated expression in this matched cohort. **(E and F)** Gene-level regression coefficients for GA (x axis) versus CA (y axis) in Purkinje cells (E) and granule cells (F). Each point represents one gene. (A to F), *n* = 8 PMA-matched subjects (tables S6 and S7).

**Table S1.**
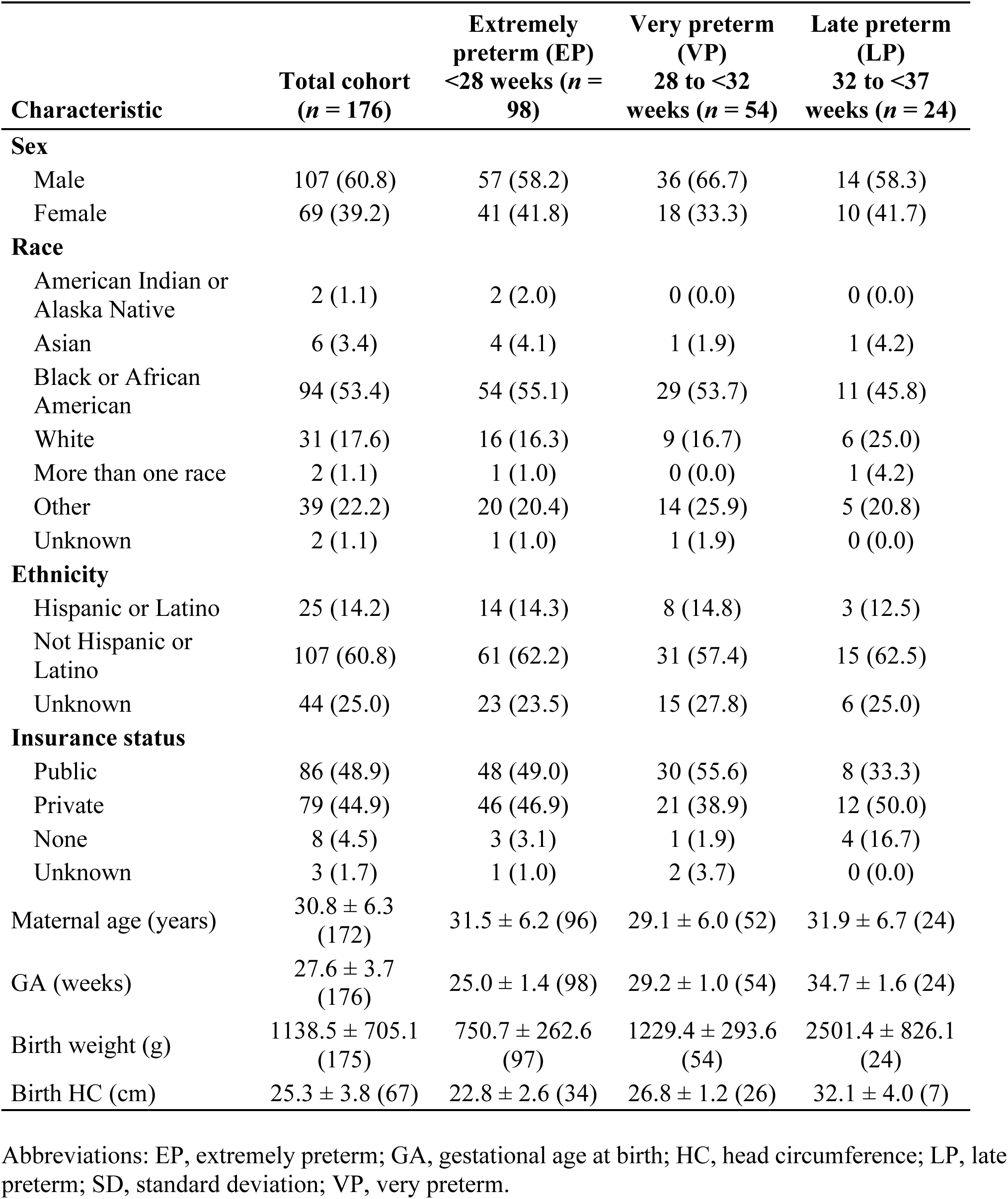
Demographic and clinical characteristics of the preterm infant in vivo cohort. Values are number (percentage) for categorical variables and mean ± SD (number of infants with available data) for continuous variables; percentages include infants of unknown status.

| **Characteristic** | **Total cohort**<br>**(*n* = 176)** | **Extremely**<br>**preterm (EP)**<br>**<28 weeks (*n* =**<br>**98)** | **Very preterm**<br>**(VP)**<br>**28 to <32**<br>**weeks (*n* = 54)** | **Late preterm**<br>**(LP)**<br>**32 to <37**<br>**weeks (*n* = 24)** |
| --- | --- | --- | --- | --- |
| **Sex** |  |  |  |  |
| Male | 107 (60.8) | 57 (58.2) | 36 (66.7) | 14 (58.3) |
| Female | 69 (39.2) | 41 (41.8) | 18 (33.3) | 10 (41.7) |
| **Race** |  |  |  |  |
| American Indian or<br>Alaska Native | 2 (1.1) | 2 (2.0) | 0 (0.0) | 0 (0.0) |
| Asian | 6 (3.4) | 4 (4.1) | 1 (1.9) | 1 (4.2) |
| Black or African<br>American | 94 (53.4) | 54 (55.1) | 29 (53.7) | 11 (45.8) |
| White | 31 (17.6) | 16 (16.3) | 9 (16.7) | 6 (25.0) |
| More than one race | 2 (1.1) | 1 (1.0) | 0 (0.0) | 1 (4.2) |
| Other | 39 (22.2) | 20 (20.4) | 14 (25.9) | 5 (20.8) |
| Unknown | 2 (1.1) | 1 (1.0) | 1 (1.9) | 0 (0.0) |
| **Ethnicity** |  |  |  |  |
| Hispanic or Latino | 25 (14.2) | 14 (14.3) | 8 (14.8) | 3 (12.5) |
| Not Hispanic or<br>Latino | 107 (60.8) | 61 (62.2) | 31 (57.4) | 15 (62.5) |
| Unknown | 44 (25.0) | 23 (23.5) | 15 (27.8) | 6 (25.0) |
| **Insurance status** |  |  |  |  |
| Public | 86 (48.9) | 48 (49.0) | 30 (55.6) | 8 (33.3) |
| Private | 79 (44.9) | 46 (46.9) | 21 (38.9) | 12 (50.0) |
| None | 8 (4.5) | 3 (3.1) | 1 (1.9) | 4 (16.7) |
| Unknown | 3 (1.7) | 1 (1.0) | 2 (3.7) | 0 (0.0) |
| Maternal age (years) | 30.8 ± 6.3<br>(172) | 31.5 ± 6.2 (96) | 29.1 ± 6.0 (52) | 31.9 ± 6.7 (24) |
| GA (weeks) | 27.6 ± 3.7<br>(176) | 25.0 ± 1.4 (98) | 29.2 ± 1.0 (54) | 34.7 ± 1.6 (24) |
| Birth weight (g) | 1138.5 ± 705.1<br>(175) | 750.7 ± 262.6<br>(97) | 1229.4 ± 293.6<br>(54) | 2501.4 ± 826.1<br>(24) |
| Birth HC (cm) | 25.3 ± 3.8 (67) | 22.8 ± 2.6 (34) | 26.8 ± 1.2 (26) | 32.1 ± 4.0 (7) |

Abbreviations: EP, extremely preterm; GA, gestational age at birth; HC, head circumference; LP, late preterm; SD, standard deviation; VP, very preterm.

**Table S2.**
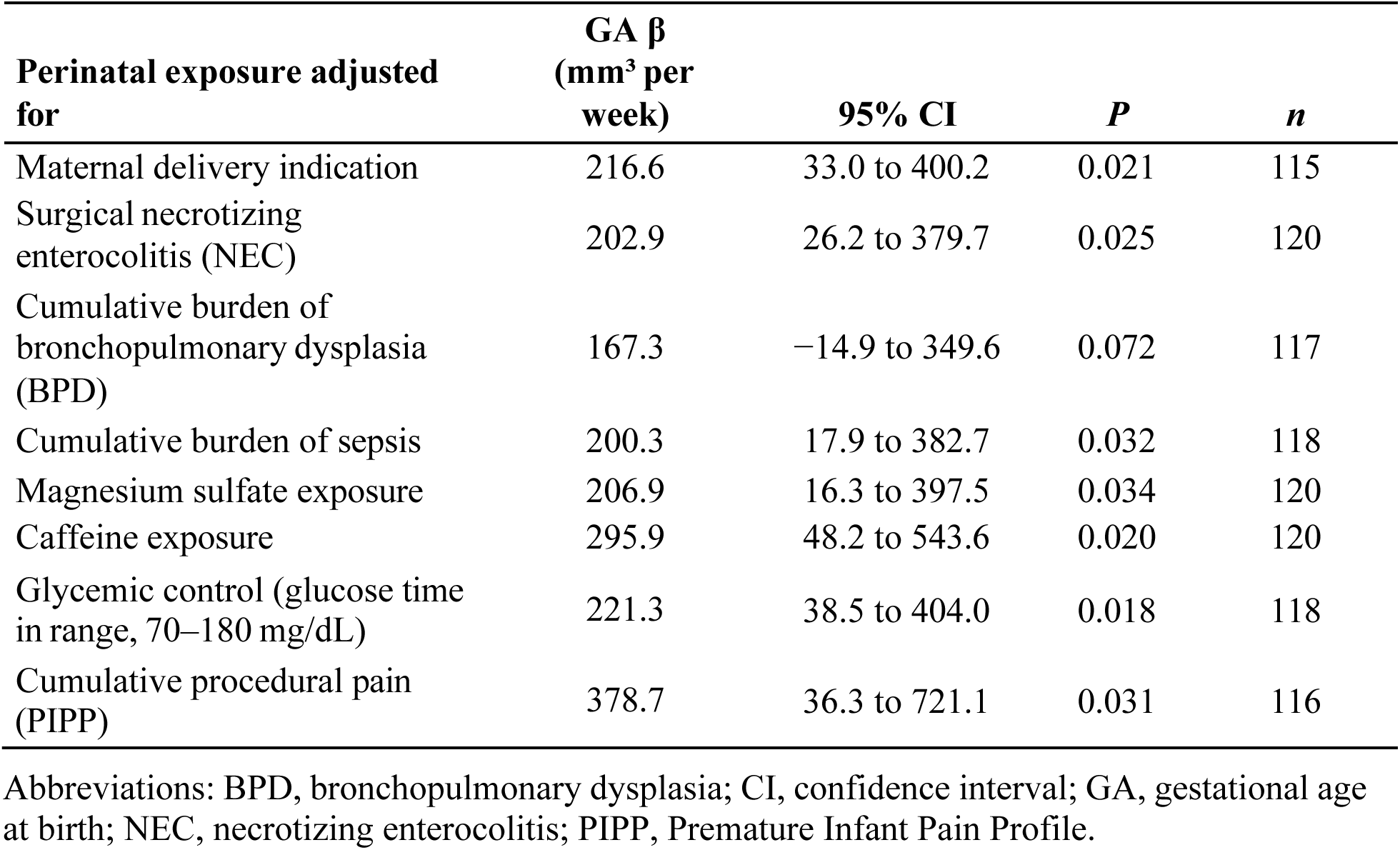
Gestational age effect on term-equivalent age cerebellar volume, adjusted for individual perinatal exposures in the in vivo cohort. Cerebellar volume was modeled on post-menstrual age at scan and one perinatal exposure at a time; gestational age (GA) was then added and tested by nested F test. Unadjusted GA effect: β = 230.2 mm³ per week (95% CI 50.0 to 410.4; *n* = 120).

**Table S3.**
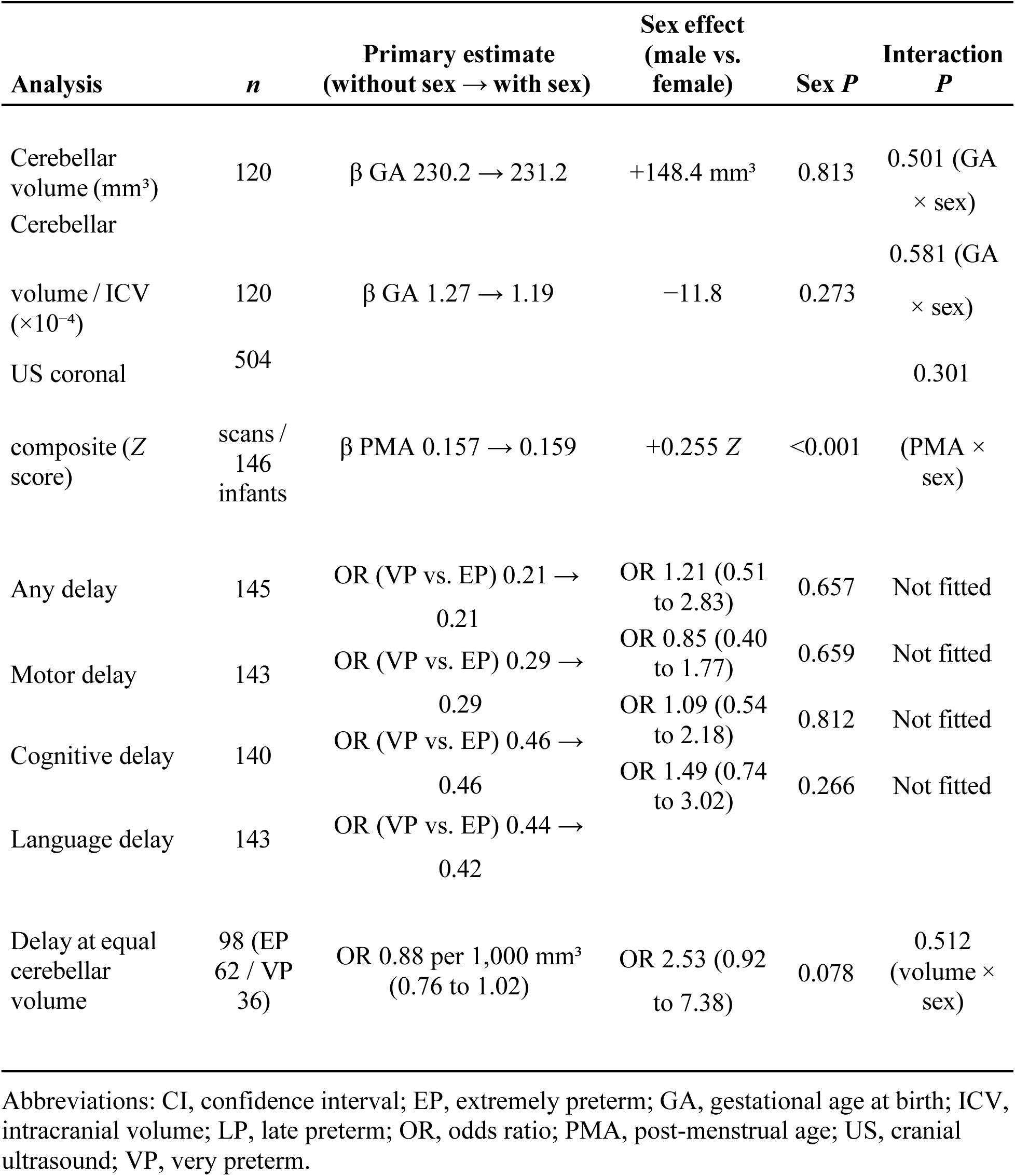
Effect of infant sex on the primary in vivo analyses. Each primary estimate is given without and with infant sex in the model. β denotes a regression slope; delay odds ratios compare VP with EP and sex odds ratios are male versus female; values in parentheses are 95% CIs, given for odds ratios only. Sex distribution: EP = 41 female / 57 male, VP = 18 female / 36 male, LP = 10 female / 14 male (*n* = 176); sex was not associated with gestational age group (χ² *P* = 0.569) or with gestational age (female 27.5 ± 3.8 versus male 27.7 ± 3.6 weeks, *P* = 0.766).

**Table S4.**
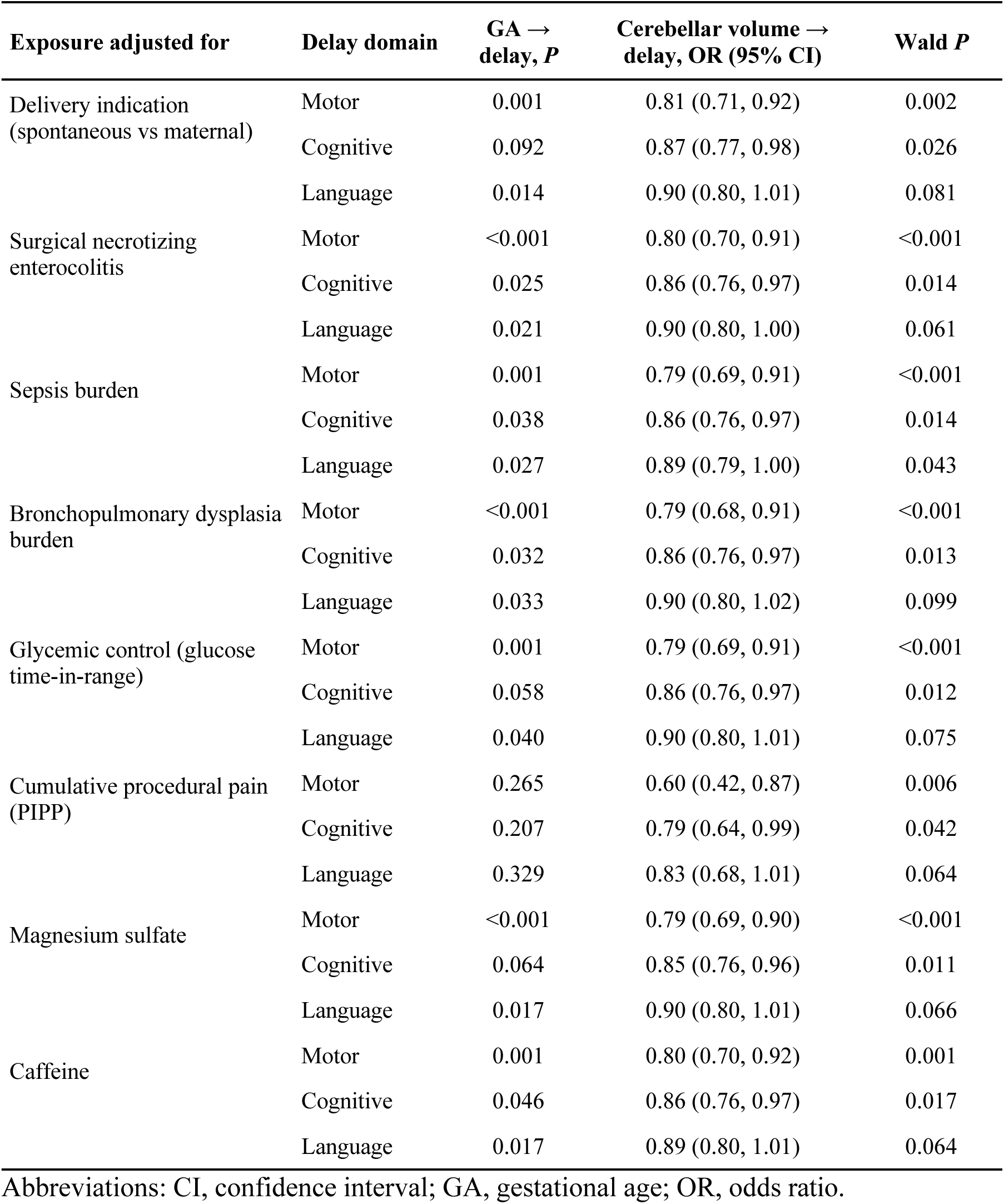
Gestational age and cerebellar volume as predictors of neurodevelopmental delay, adjusted for individual perinatal exposure in vivo cohort. Logistic regression of motor, cognitive, and language delay, with one perinatal exposure added per model, restricted to extremely and very preterm infants. Gestational age (GA) → delay: likelihood-ratio χ² *P* for the GA group term (*n* = 140-143). Cerebellar volume → delay: odds ratio (OR) per 1,000 mm³ with 95% confidence interval (CI) and Wald *P* for term-equivalent age cerebellar volume (*n* = 90–97).

**Table S5.**
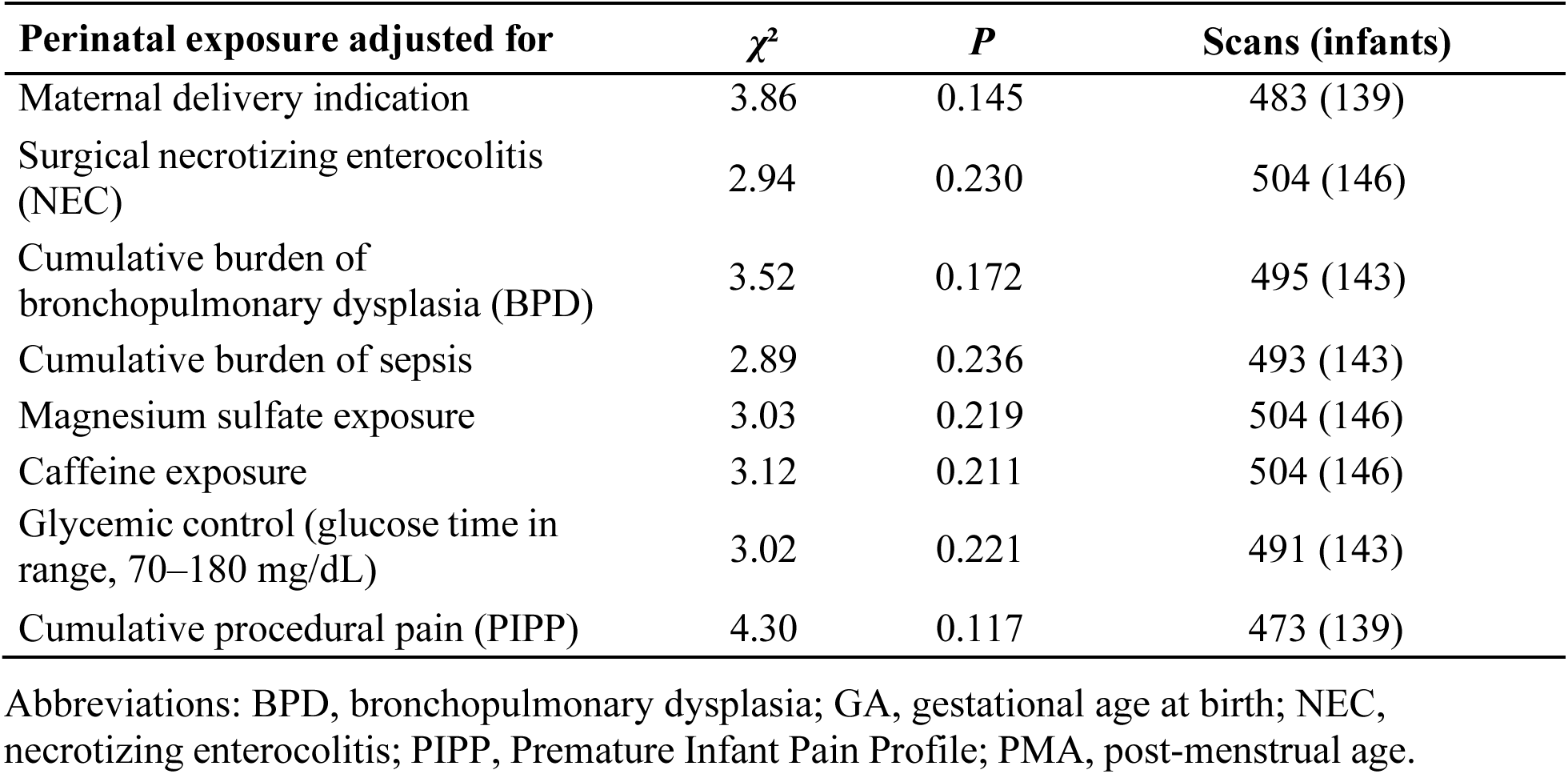
Differences in coronal cerebellar growth rate between gestational age groups, adjusted for individual perinatal exposures. Gestational age (GA) group, post-menstrual age (PMA), and the exposure were entered in a base model. For each perinatal exposure, this was compared by likelihood-ratio test with a model additionally including the GA group × PMA interaction.

**Table S6.**
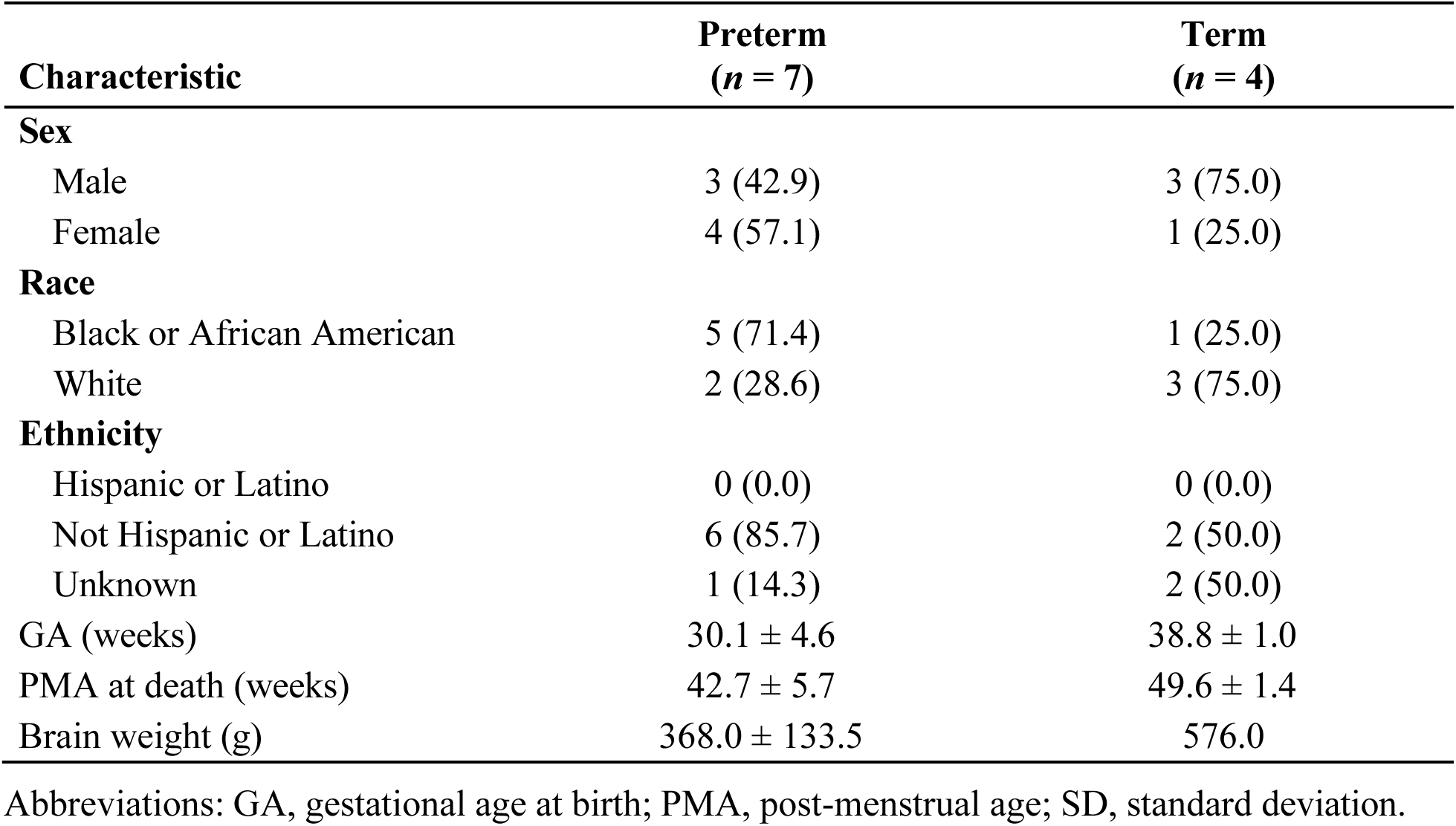
Demographic and clinical characteristics of the postmortem spatial transcriptomic cohort, by preterm and term group. Values are number (percentage) for categorical variables and mean ± SD for continuous variables.

| **Characteristic** | **Preterm**<br>**(*n* = 7)** | **Term**<br>**(*n* = 4)** |
| --- | --- | --- |
| **Sex** |  |  |
| Male | 3 (42.9) | 3 (75.0) |
| Female | 4 (57.1) | 1 (25.0) |
| **Race** |  |  |
| Black or African American | 5 (71.4) | 1 (25.0) |
| White | 2 (28.6) | 3 (75.0) |
| **Ethnicity** |  |  |
| Hispanic or Latino | 0 (0.0) | 0 (0.0) |
| Not Hispanic or Latino | 6 (85.7) | 2 (50.0) |
| Unknown | 1 (14.3) | 2 (50.0) |
| GA (weeks) | 30.1 ± 4.6 | 38.8 ± 1.0 |
| PMA at death (weeks) | 42.7 ± 5.7 | 49.6 ± 1.4 |
| Brain weight (g) | 368.0 ± 133.5 | 576.0 |

Abbreviations: GA, gestational age at birth; PMA, post-menstrual age; SD, standard deviation.

**Table S7.**
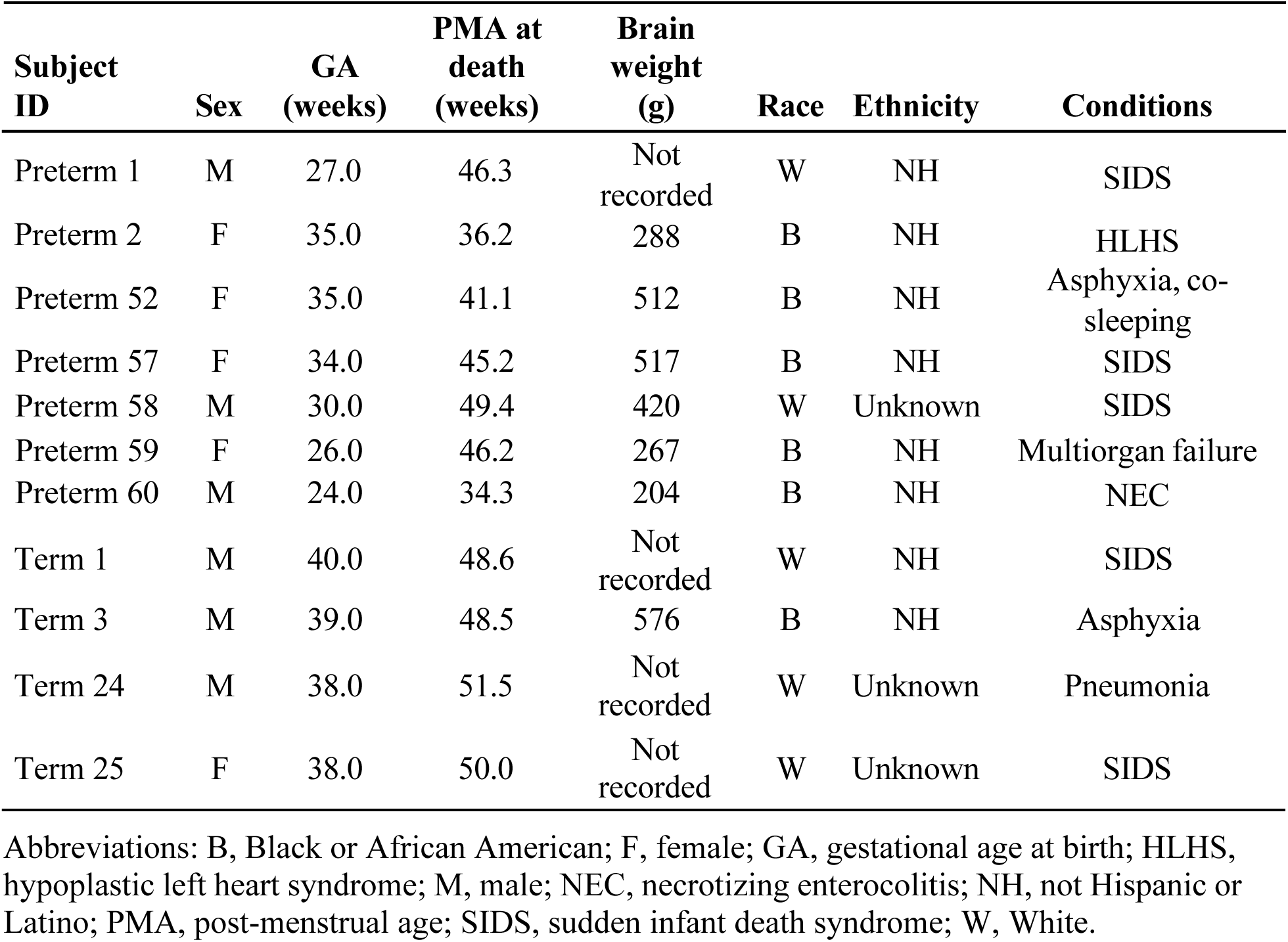
Demographic and clinical characteristics of the postmortem spatial transcriptomic cohort, by individual subject. Data for the 7 preterm and 4 term subjects analyzed by spatial transcriptomics. Conditions lists the conditions documented at or contributing to death.

**Table S8.**
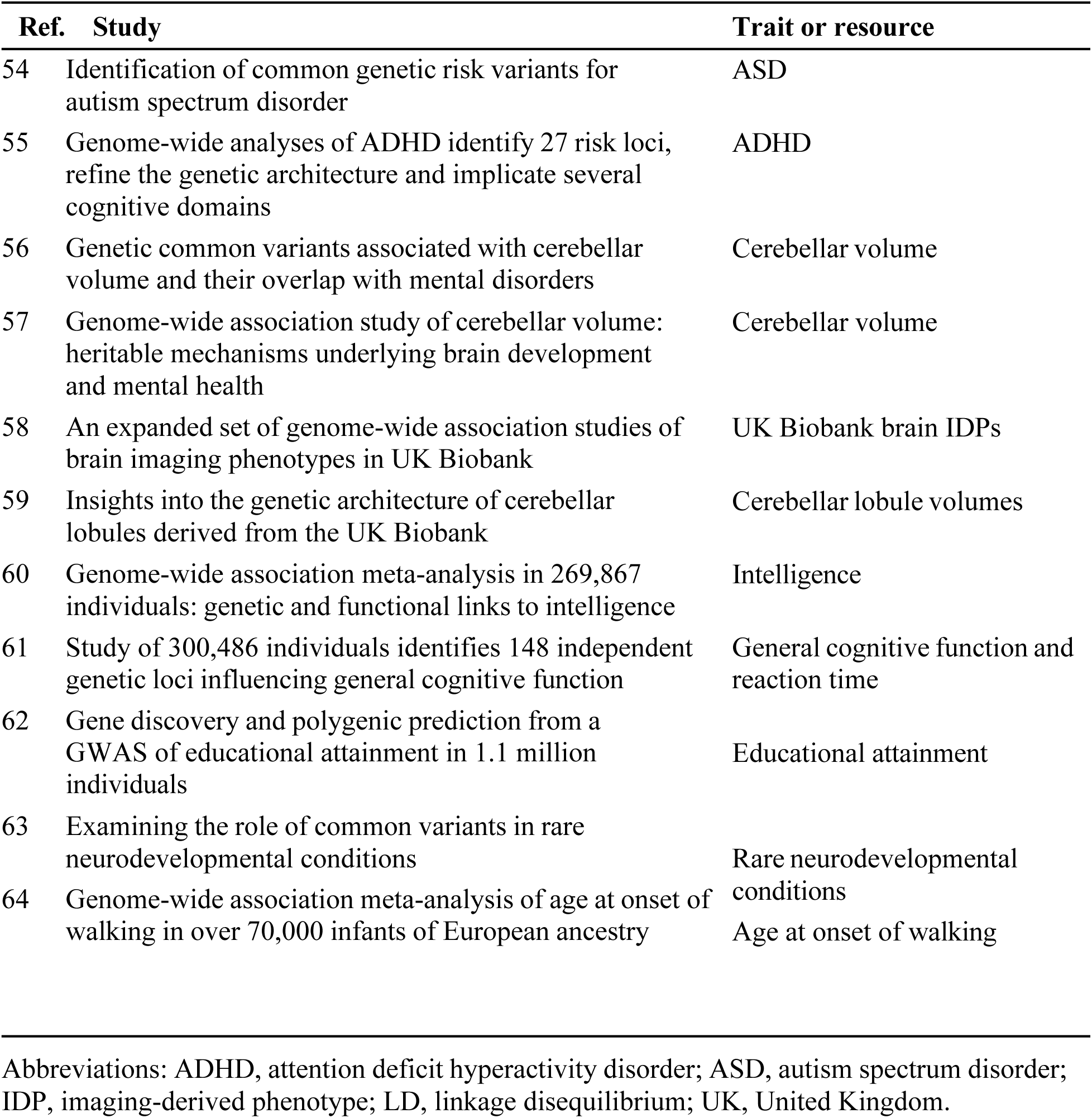
Datasets used in the partitioned heritability analysis. Genome-wide association study (GWAS) reference datasets used for partitioned heritability analysis by linkage disequilibrium score regression. Full citations appear in the main reference list under the indicated reference numbers.

**Table S9.**
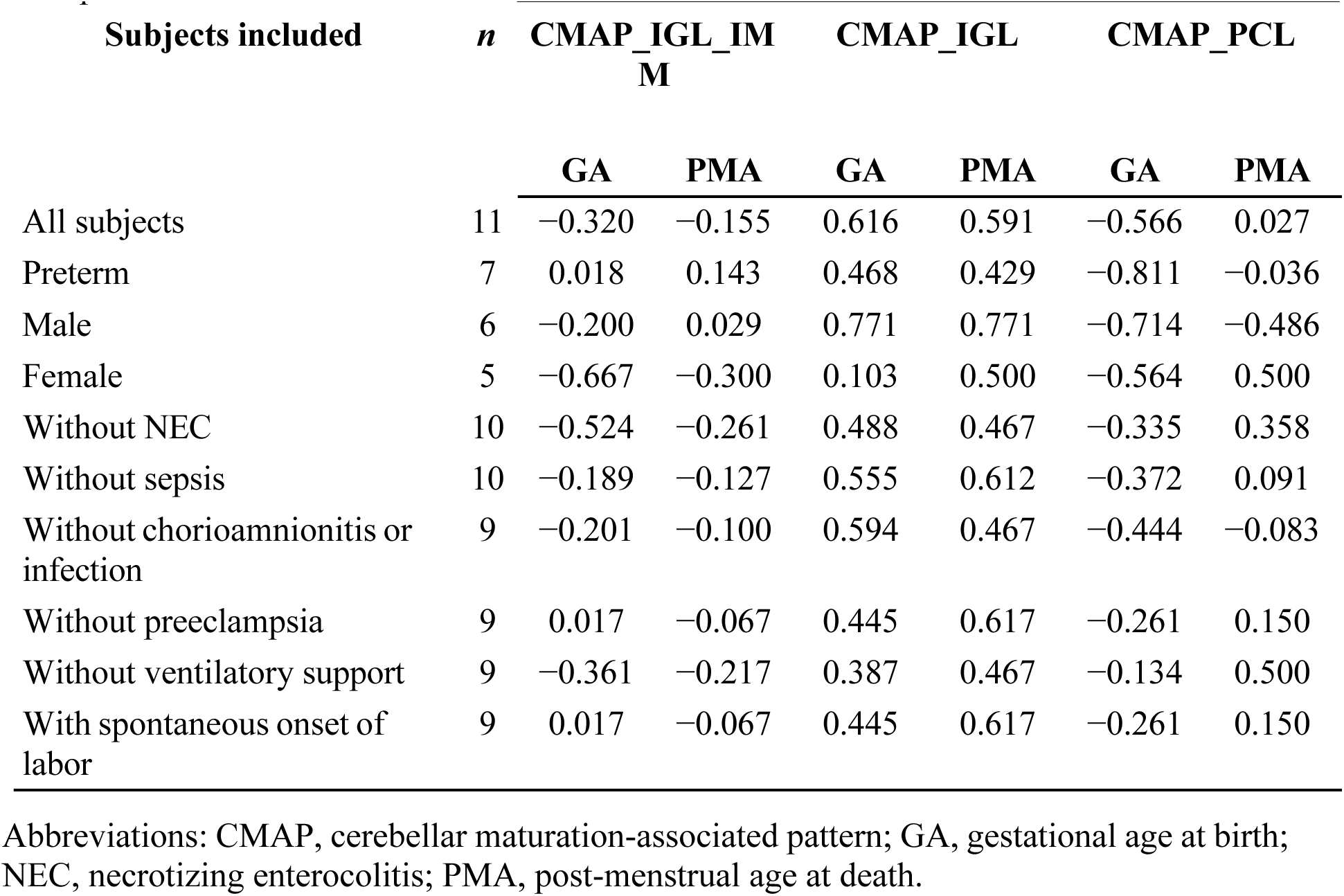
Correlation of CMAP activity with gestational age at birth and post-menstrual age at death within clinical subgroups. Spearman rank-correlation coefficients (rho) between subject-level CMAP activity and gestational age at birth or post-menstrual age at death. The first row is the full cohort; each subsequent row is restricted to the subjects indicated. Coefficients are descriptive; no *P* values are reported.

**Table S10.**
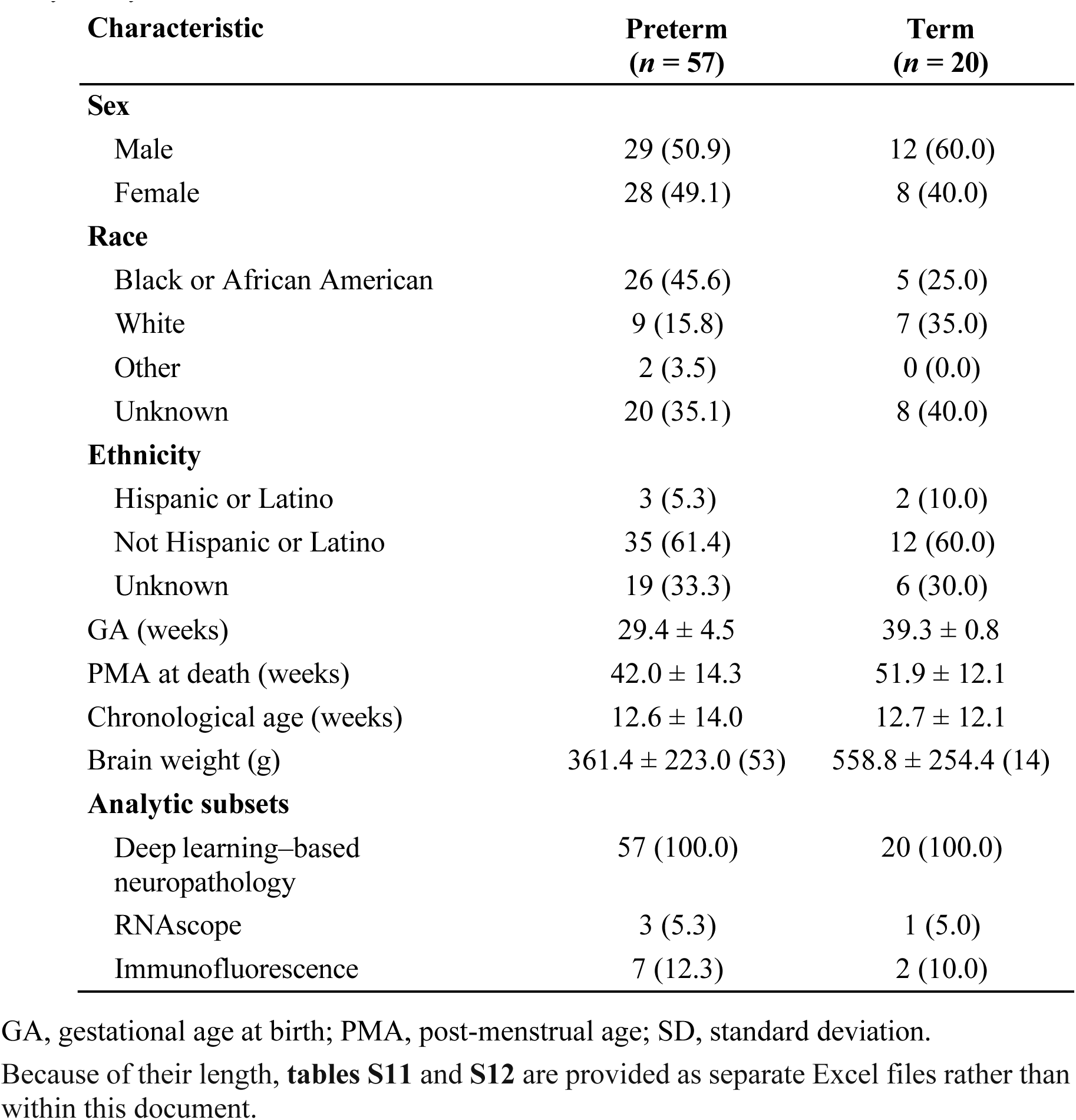
Demographic and clinical characteristics of the postmortem neuropathology cohort and its analytic subsets, by preterm and term group. Values are number (percentage) for categorical variables and mean ± SD (number of subjects with available data) for continuous variables. The final rows give the number of subjects analyz<u>ed by each method.</u>

| **Characteristic** | **Preterm**<br>**(*n* = 57)** | **Term**<br>**(*n* = 20)** |
| --- | --- | --- |
| **Sex** |  |  |
| Male | 29 (50.9) | 12 (60.0) |
| Female | 28 (49.1) | 8 (40.0) |
| **Race** |  |  |
| Black or African American | 26 (45.6) | 5 (25.0) |
| White | 9 (15.8) | 7 (35.0) |
| Other | 2 (3.5) | 0 (0.0) |
| Unknown | 20 (35.1) | 8 (40.0) |
| **Ethnicity** |  |  |
| Hispanic or Latino | 3 (5.3) | 2 (10.0) |
| Not Hispanic or Latino | 35 (61.4) | 12 (60.0) |
| Unknown | 19 (33.3) | 6 (30.0) |
| GA (weeks) | 29.4 ± 4.5 | 39.3 ± 0.8 |
| PMA at death (weeks) | 42.0 ± 14.3 | 51.9 ± 12.1 |
| Chronological age (weeks) | 12.6 ± 14.0 | 12.7 ± 12.1 |
| Brain weight (g) | 361.4 ± 223.0 (53) | 558.8 ± 254.4 (14) |
| **Analytic subsets** |  |  |
| Deep learning–based neuropathology | 57 (100.0) | 20 (100.0) |
| RNAscope | 3 (5.3) | 1 (5.0) |
| Immunofluorescence | 7 (12.3) | 2 (10.0) |

GA, gestational age at birth; PMA, post-menstrual age; SD, standard deviation.

**Table S13.**
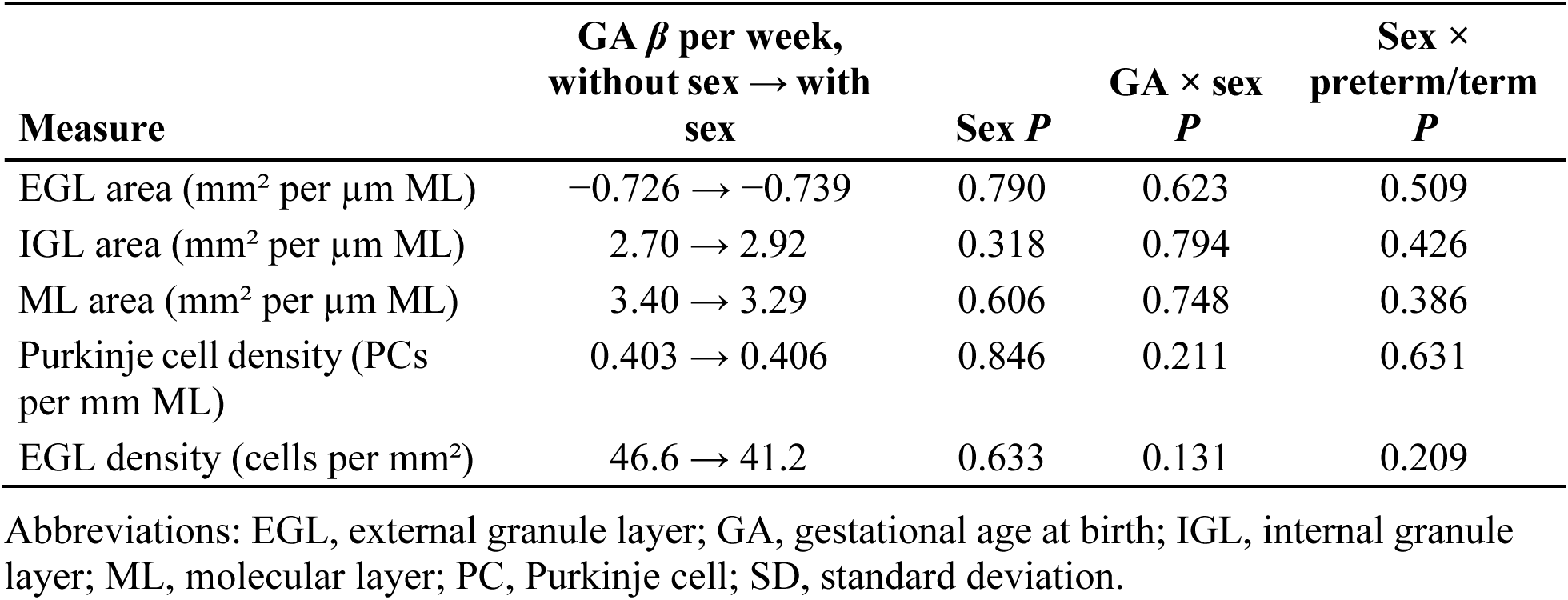
Effect of sex on the postmortem neuropathology analyses. Each gestational age (GA) coefficient is given without and with infant sex in the model, alongside *P* values for the sex effect and for the two interactions. All models include all 77 subjects. Sex was not associated with preterm/term status (Fisher exact *P* = 0.604) or with GA (female 31.0 ± 6.0 versus male 32.8 ± 5.6 weeks, *P* = 0.175).

| **Measure** | **GA $\beta$ per week,**<br>**without sex → with**<br>**sex** | **Sex *P*** | **GA × sex**<br>***P*** | **Sex ×**<br>**preterm/term**<br>***P*** |
| --- | --- | --- | --- | --- |
| EGL area (mm<sup>2</sup> per μm ML) | -0.726 → -0.739 | 0.790 | 0.623 | 0.509 |
| IGL area (mm<sup>2</sup> per μm ML) | 2.70 → 2.92 | 0.318 | 0.794 | 0.426 |
| ML area (mm<sup>2</sup> per μm ML) | 3.40 → 3.29 | 0.606 | 0.748 | 0.386 |
| Purkinje cell density (PCs<br>per mm ML) | 0.403 → 0.406 | 0.846 | 0.211 | 0.631 |
| EGL density (cells per mm<sup>2</sup>) | 46.6 → 41.2 | 0.633 | 0.131 | 0.209 |

Abbreviations: EGL, external granule layer; GA, gestational age at birth; IGL, internal granule layer; ML, molecular layer; PC, Purkinje cell; SD, standard deviation.

**Table S14.**
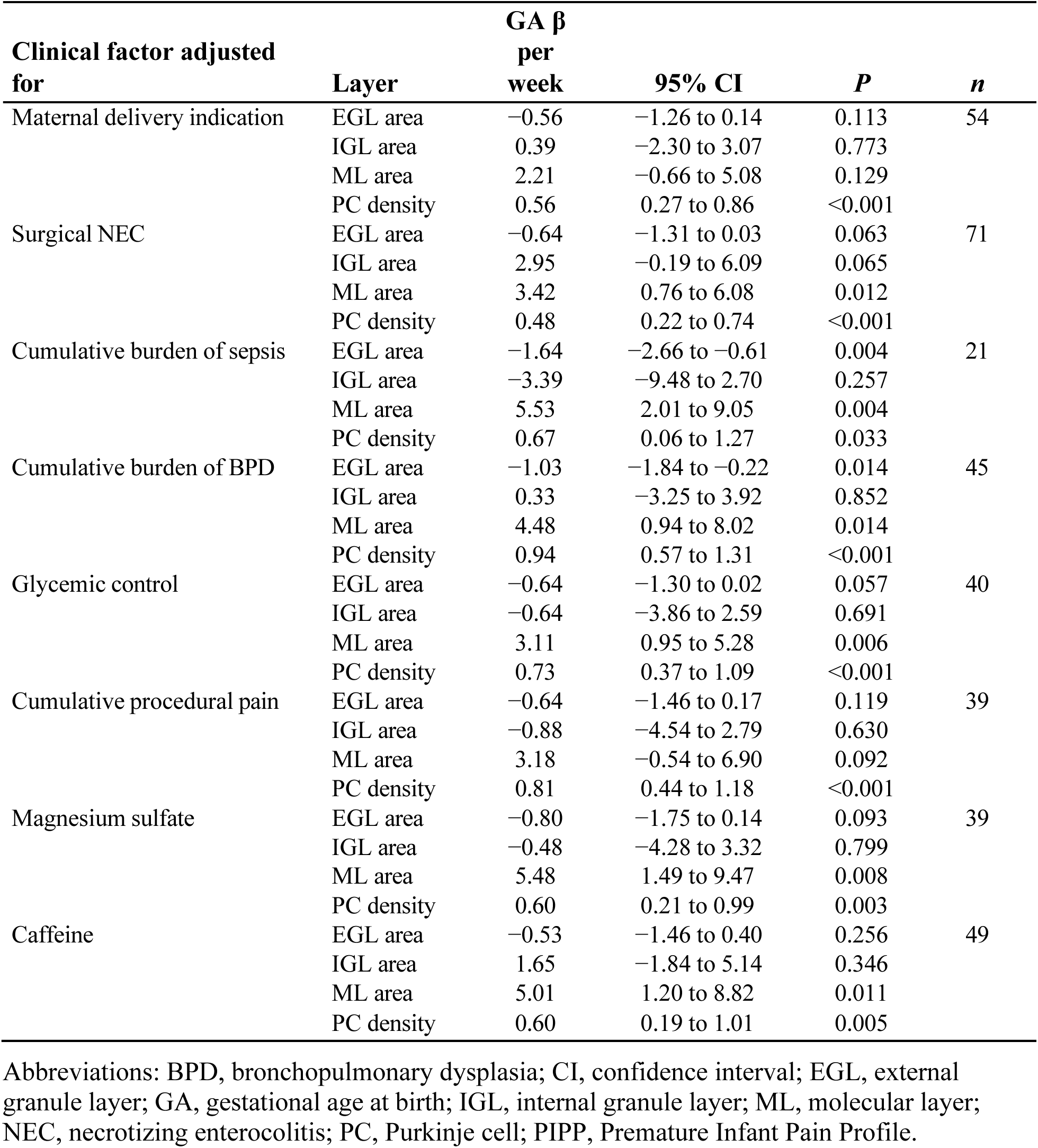
Gestational age effect on cerebellar layer architecture after adjustment for individual clinical factors in the postmortem neuropathology cohort. For each clinical factor, each layer measure was modeled on that factor and gestational age (GA); the GA term was tested by nested F test and only the GA coefficient is shown. Glycemic control is glucose time in range (70–180 mg/dL) and cumulative procedural pain is the summed Premature Infant Pain Profile (PIPP) score.

**Table S15.**
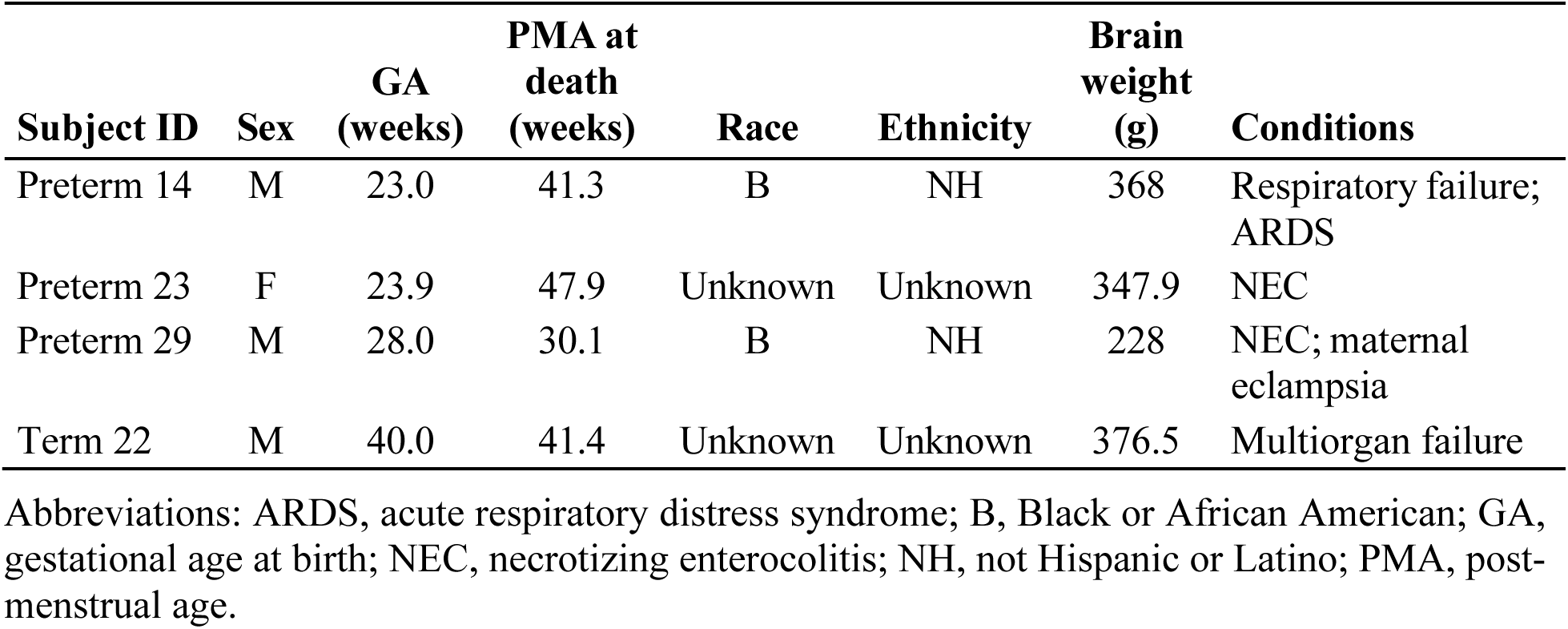
Demographic and clinical characteristics of the postmortem subjects analyzed by RNAscope in situ hybridization. The four subjects, three preterm and one term, drawn from the postmortem neuropathology cohort. Conditions lists the conditions documented at or contributing to death.

**Table S16.**
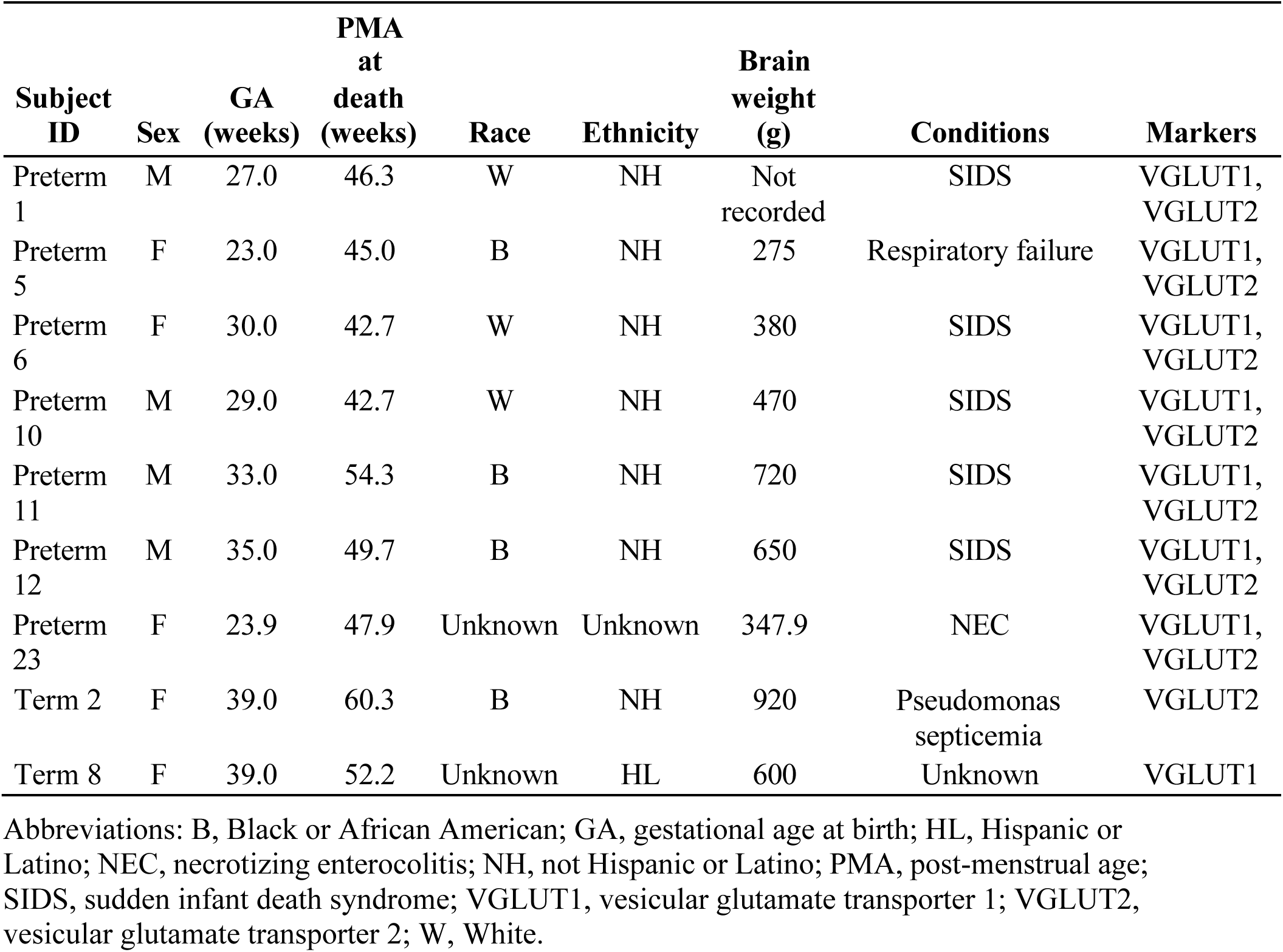
Demographic and clinical characteristics of the immunofluorescence subjects in the postmortem neuropathology cohort. Data for each of the 9 immunofluorescence subjects. Conditions lists the conditions documented at or contributing to death. All seven preterm subjects were stained for both VGLUT1 and VGLUT2; the term subjects differ by marker (Term 8 for VGLUT1, Term 2 for VGLUT2), so each marker was assessed in eight subjects and nine subjects contributed in total.

**Table S17.**
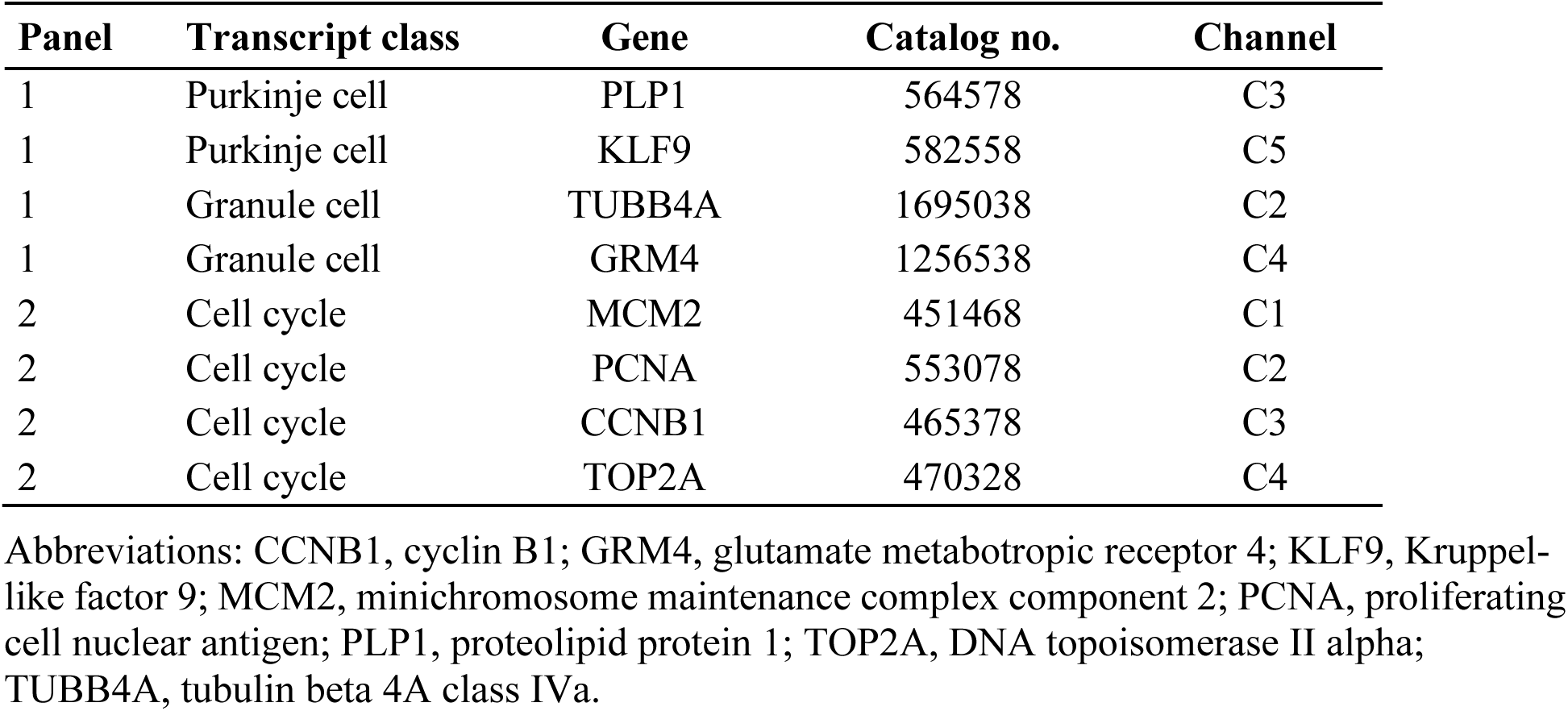
RNAscope probe panels. All probes are RNAscope 2.5 LS Probe, Homo sapiens (Advanced Cell Diagnostics).

**Table S18.**
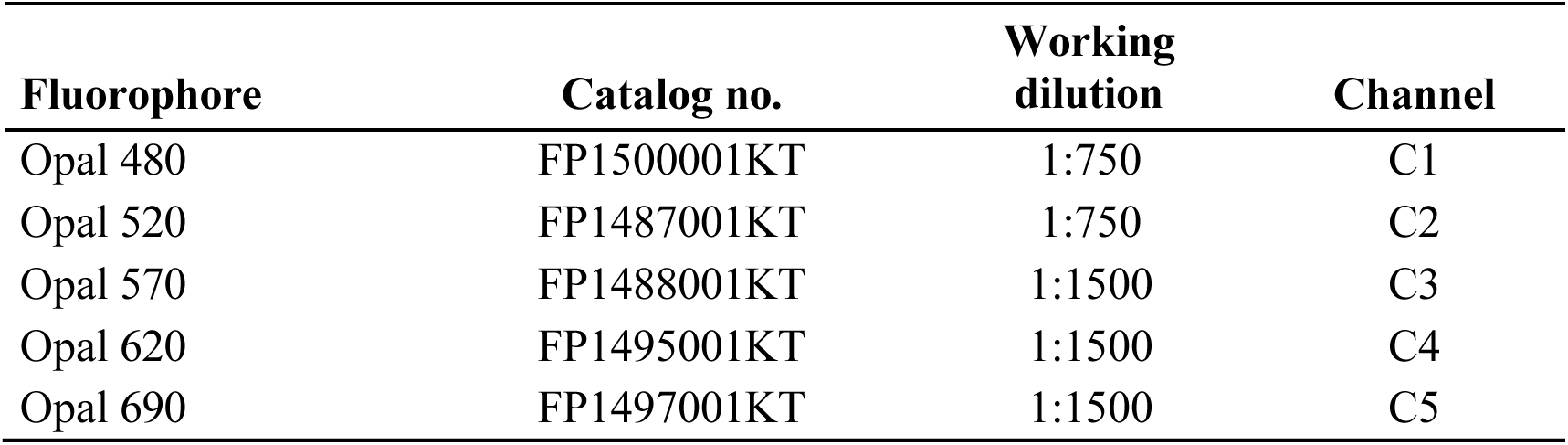
Opal fluorophore and channel assignments for the RNAscope probe panels. All reagents are Opal Reagent Packs (Akoya Biosciences).

**Table S19.**
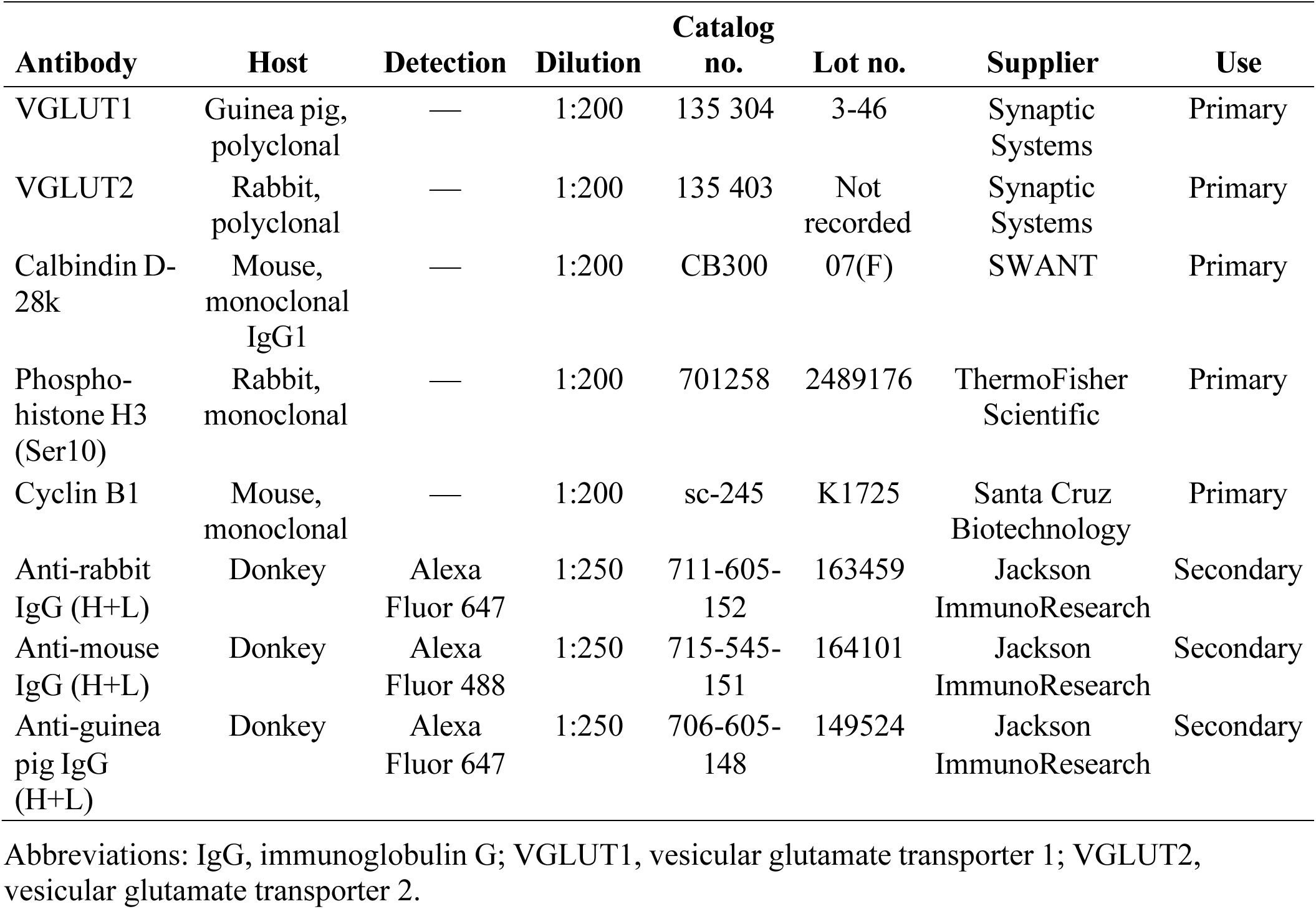
Antibodies used for immunofluorescent analyses.

**Table S20.**
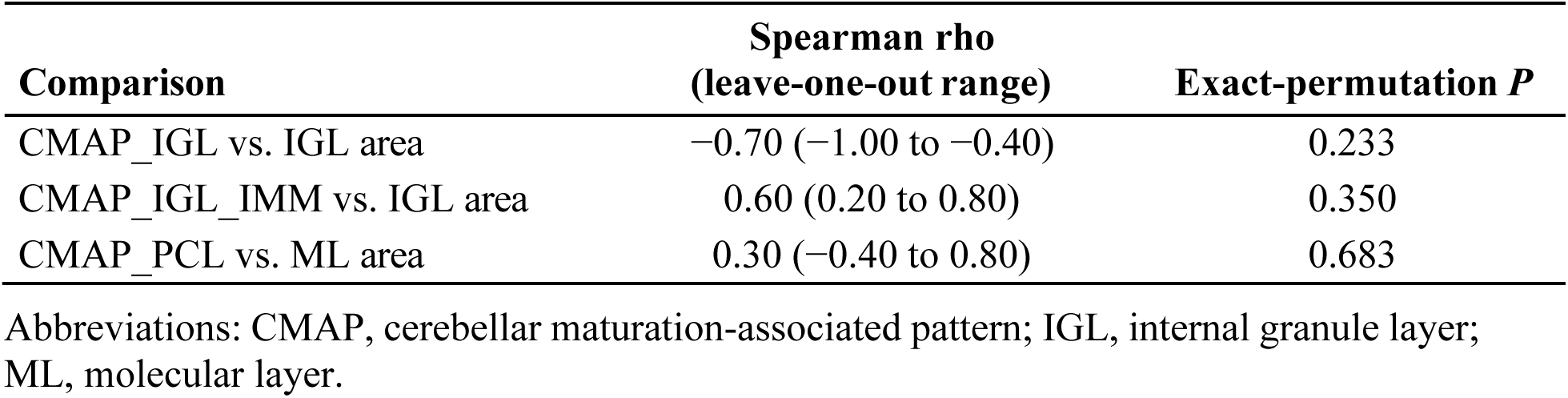
Intra-subject correlation between transcriptomic and neuropathological cerebellar metrics. Spearman rank correlation coefficients (rho) between per-subject cerebellar maturation-associated pattern (CMAP) activity and normalized cerebellar layer surface areas from deep learning–based neuropathology, in the five postmortem subjects analyzed by both modalities (Preterm 1, 2 and 52; Term 1 and 3). Per-subject CMAP activity is the mean over layer-matched spatial spots: two-sided exact-permutation *P* values (all 120 permutations) are given for reference.

