## Supplemental Material for manuscript for "Intrinsic Gestational Timing Governs Human Cerebellar Development After Preterm Birth"

[1] Center for Neuroscience Research and Sheikh Zayed Institute for Pediatric Surgical Innovation, Children's National Hospital, Washington, DC 20010, USA
[2] Department of Pediatrics, The George Washington University School of Medicine and Health Sciences, Washington, DC 20037, USA
[3] Department of Computational Medicine & Bioinformatics, University of Michigan, Ann Arbor, MI 48109, USA
[4] Johns Hopkins University School of Medicine, Baltimore, MD 21205, USA
[5] Children's Hospital of Philadelphia, University of Pennsylvania School of Medicine, Philadelphia, PA, 19104, USA
[6] Life and Health Sciences Research Institute (ICVS); Unidade Local de Saúde de Santo António, Braga 4710-057, Portugal
[7] Department of Neuroradiology, Great Ormond Street Hospital NHS Foundation Trust, London WC1N 3JH, United Kingdom
[8] Department of Pathology, St. Olavs Hospital, Trondheim University Hospital, 7030 Trondheim, Norway
[9] Department of Clinical and Molecular Medicine, Faculty of Medicine and Health Sciences, NTNU, Norwegian University of Science and Technology, 7491 Trondheim, Norway
[10] Seattle Children's Research Institute, Seattle, WA 98101, USA
[11] University of Washington School of Medicine, Seattle, WA 98195, USA
[12] Department of Neonatology, Children's National Hospital, Washington, DC 20010, USA
† These Authors contributed equally to this work
* Correspondence to:
Panagiotis Kratimenos, MD, PhD


Vittorio Gallo, PhD


**This PDF file includes:**

-Materials and Methods
-Figs. S1 to S18
-Tables S1-S10 and tables S13-S20

**Other Supplementary Materials for this manuscript include the following:**

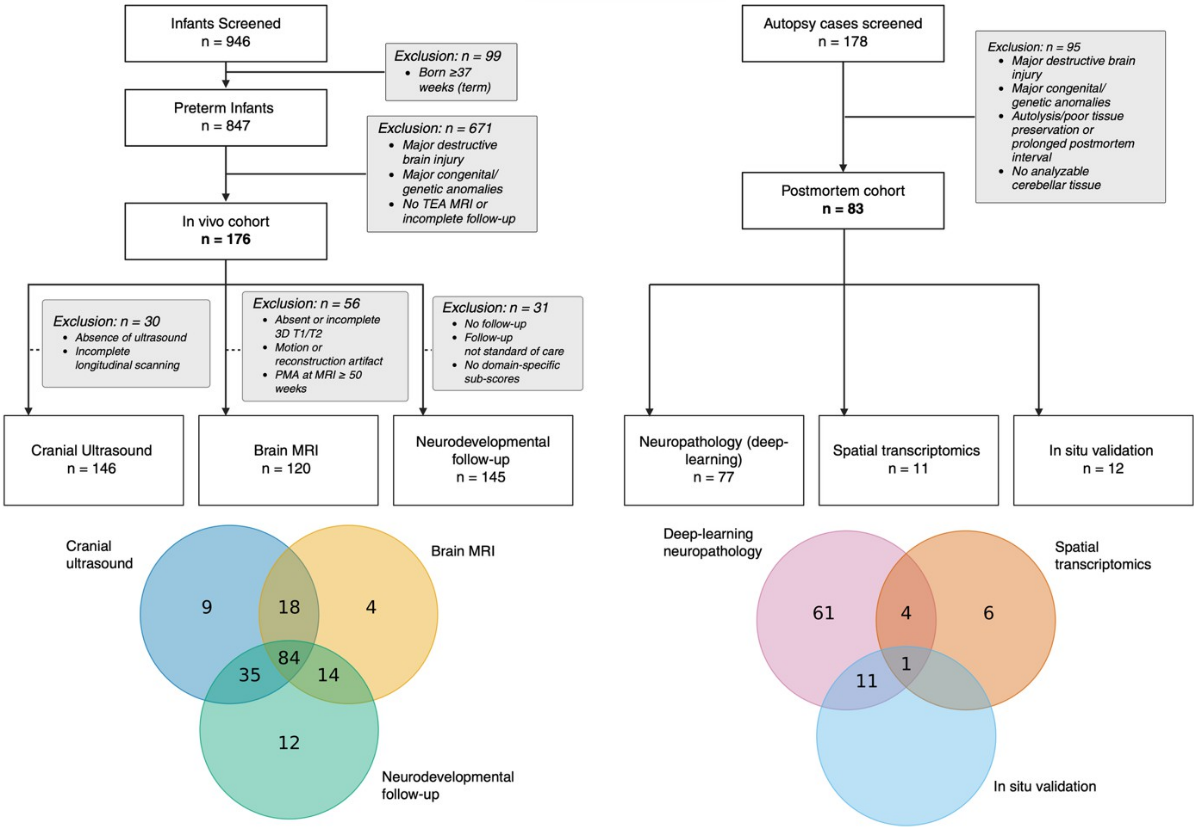


**Fig. S1. Cohort selection and subject overlap across in vivo and postmortem analyses.** Left, in vivo cohort selection. Of 946 infants screened, 99 born at ≥37 weeks were excluded; of the remaining 847 preterm infants, 671 were excluded for major destructive brain injury, major congenital or genetic anomalies, absence of term-equivalent age (TEA) MRI, or incomplete follow-up, yielding the in vivo cohort ($n$ = 176). Three analytic cohorts were derived: cranial ultrasound ($n$ = 146; EP, 80; VP, 45; LP, 21), brain MRI ($n$ = 120; EP, 65; VP, 37; LP, 18), and neurodevelopmental follow-up ($n$ = 145; EP, 94; VP, 51). Modality-specific exclusions and reasons are indicated in the flow diagram. Right, postmortem cohort selection. Of 178 autopsy cases screened, 95 were excluded for major destructive brain injury, major congenital or genetic anomalies, autolysis or poor tissue preservation, prolonged postmortem interval, or absence of analyzable cerebellar tissue, yielding the postmortem cohort ($n$ = 83). Analytic cohorts comprised deep-learning neuropathology ($n$ = 77), spatial transcriptomics ($n$ = 11), and in situ validation ($n$ = 12; immunofluorescence, $n$ = 9; RNAscope, $n$ = 4; one subject assessed by both). Bottom, Venn diagrams show subject overlap across analyses within the in vivo (left) and postmortem (right) cohorts; numbers indicate unique or shared subjects. EP, extremely preterm; VP, very preterm; LP, late preterm; MRI, magnetic resonance imaging.

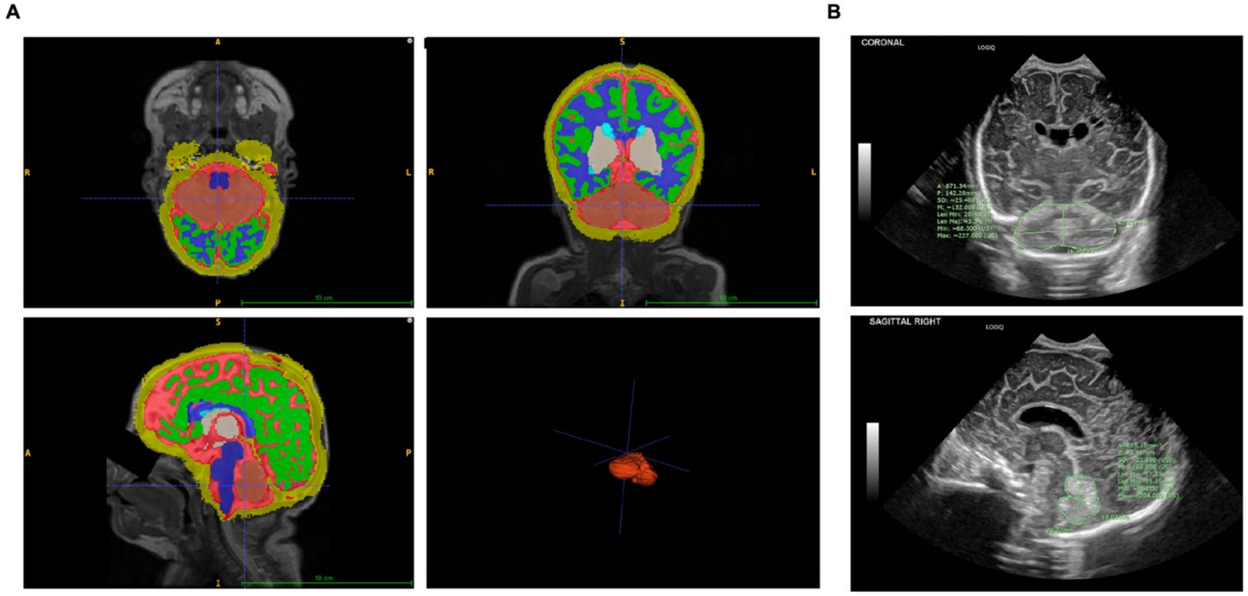


**Fig. S2. Representative MRI cerebellar segmentation and cranial ultrasound morphometry. (A)** Representative brain MRI segmentation showing segmented brain structures with the cerebellum highlighted in red in axial (top left), coronal (top right), and sagittal (bottom left) views, with the corresponding three-dimensional cerebellar reconstruction (bottom right). Segmentation labels are overlaid on the anatomical images. Scale bar, 10 cm. **(B)** Representative cranial ultrasound images showing coronal (top) and sagittal (bottom) measurement planes used for cerebellar morphometry. Coronal measurements include transcerebellar diameter and cross-sectional area; sagittal measurements include anteroposterior diameter, height, and cross-sectional area. Measurement axes and contours are shown in green.

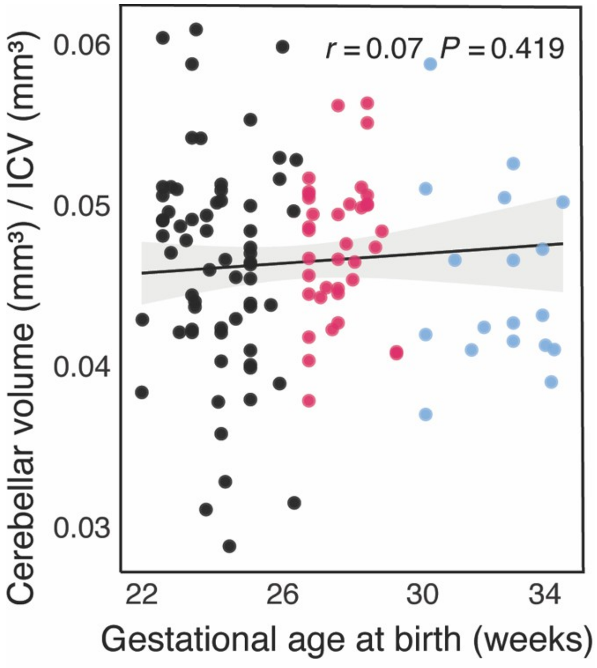


**Fig. S3. Cerebellar volume normalized to intracranial volume.**
Cerebellar volume at term-equivalent age normalized to intracranial volume (ICV) and plotted against gestational age at birth in extremely preterm (EP, black), very preterm (VP, magenta), and late preterm (LP, light blue) infants ($n$ = 120; EP = 65, VP = 37, LP = 18). Line, ordinary least-squares fit; shaded band, 95% CI; partial $r$, adjusted for post-menstrual age at MRI. The cerebellum-to-ICV ratio was not associated with gestational age at birth (partial $r = 0.07$; $P = 0.419$), indicating proportional scaling of cerebellar and intracranial volumes.

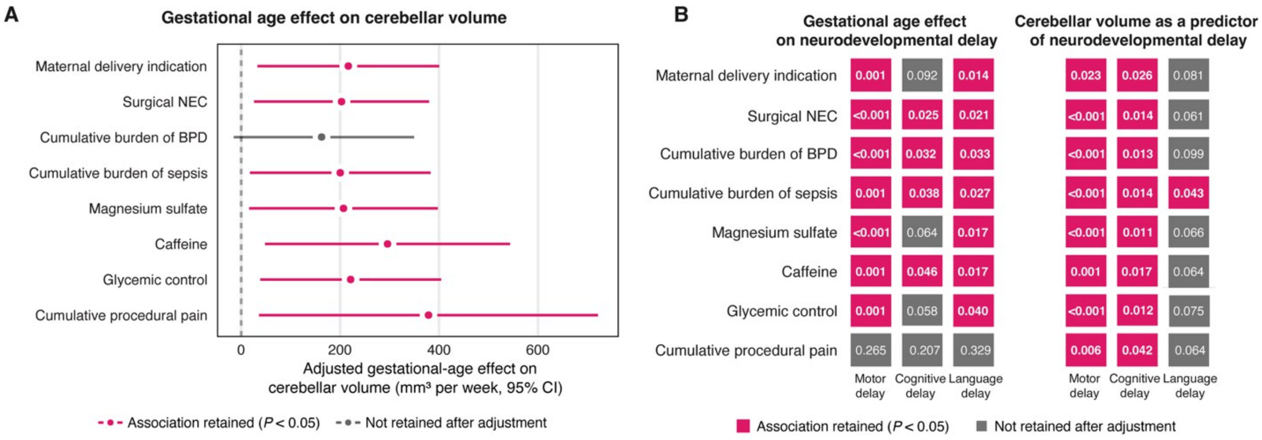


**Fig. S4. Persistence of gestational age and cerebellar volume associations after adjustment for individual perinatal exposures. (A)** Association between gestational age at birth (GA) and term-equivalent age (TEA) cerebellar volume after adjustment for individual perinatal exposures. Each row shows the GA coefficient and 95% CI from a separate model including post-menstrual age at MRI and the indicated exposure as covariates. Points, GA coefficients; horizontal lines, 95% CI; dashed line, no association. Magenta, association retained after adjustment ($P < 0.05$); gray, not retained. Model size ranged from $n = 115$ to $n = 120$. **(B)** Persistence of the associations of GA (left) and TEA cerebellar volume (right) with motor, cognitive, and language delay after adjustment for each indicated perinatal exposure. Each cell shows the $P$ value from a separate adjusted model. Magenta, association retained ($P < 0.05$); gray, not retained. Model size ranged from $n = 140$ to $n = 143$ for GA models and from $n = 90$ to $n = 97$ for cerebellar-volume models. Perinatal exposures included maternal delivery indication, surgical necrotizing enterocolitis (NEC), cumulative burden of bronchopulmonary dysplasia (BPD), cumulative burden of sepsis, magnesium sulfate, caffeine, glycemic control, and cumulative procedural pain. Each exposure was entered separately as an additive covariate; no interaction terms were included.

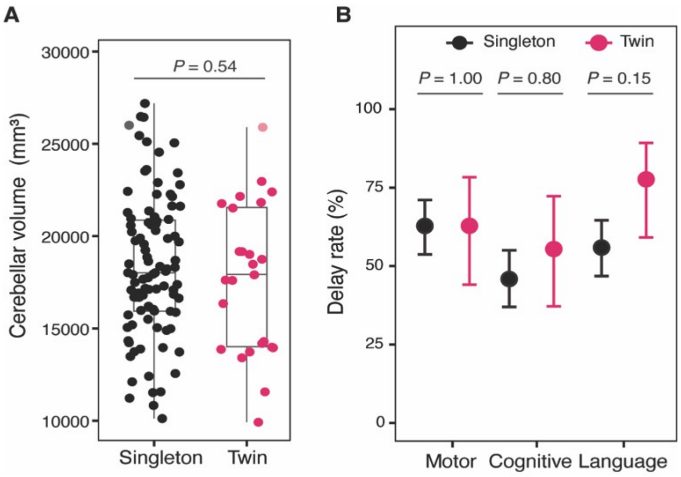


**Fig. S5. Twin status does not account for cerebellar volume or neurodevelopmental delay.** **(A)** Term-equivalent age (TEA) cerebellar volume in singletons (*n* = 95) and twins (*n* = 25) in the volumetric MRI cohort. Two-sided Wilcoxon rank-sum test; box, median and IQR; whiskers, 1.5× IQR; points, individual infants. Gestational age at birth did not differ between groups (*P* = 0.82), and inclusion of twin status did not alter the gestational age–volume association (twin-status coefficient, +136 $mm^3$; *P* = 0.86). **(B)** Motor, cognitive, and language delay rates in extremely and very preterm infants by twin status (singletons, *n* = 118; twins, *n* = 27). Points, proportion with delay; error bars, 95% Wilson CI. *P* values, two-sided Fisher's exact tests, Holm-adjusted across the three domains. No delay domain differed by twin status after adjustment. Singletons, black; twins, magenta.

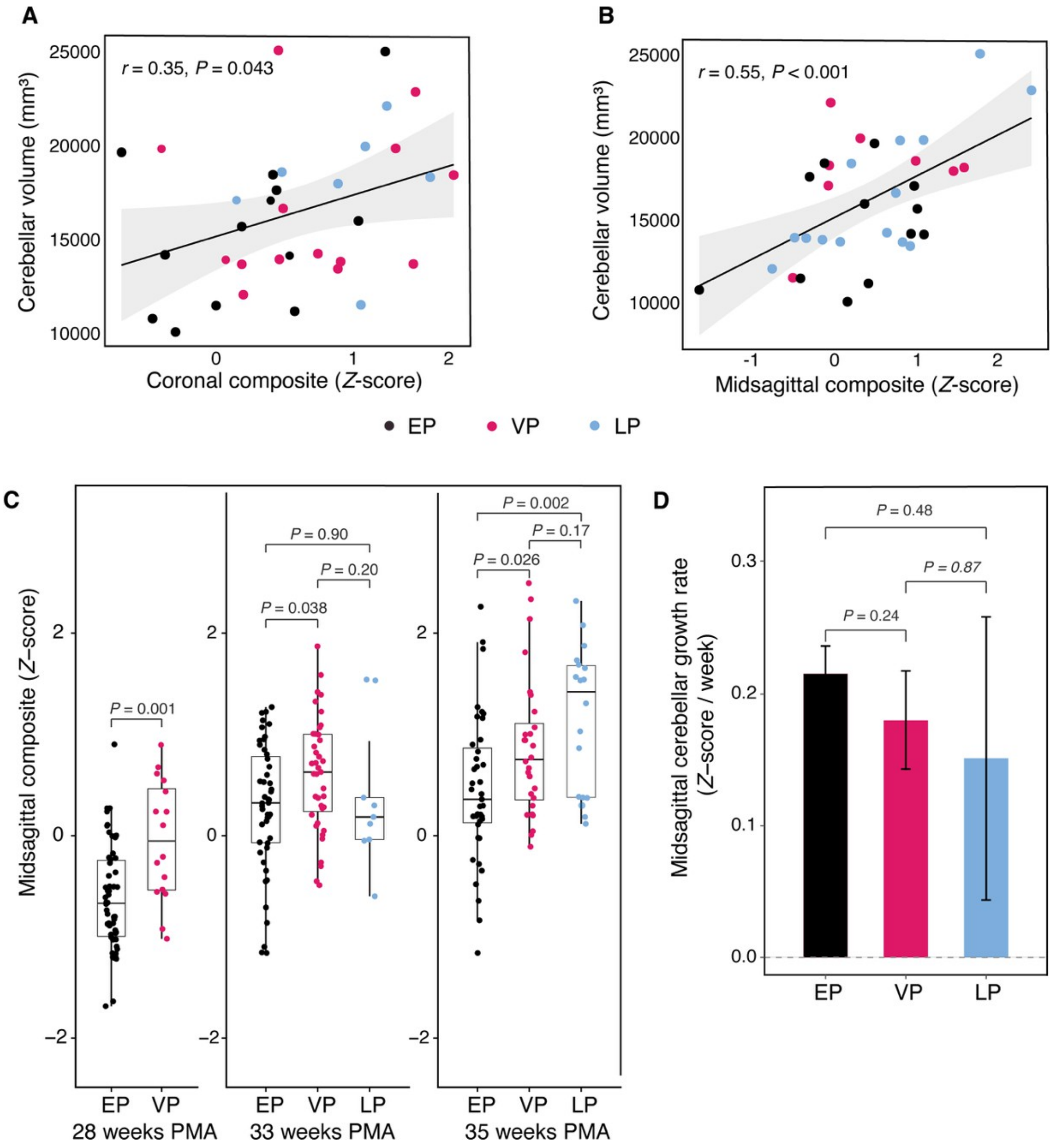


**Fig. S6. Cross-modal correlation of ultrasound and MRI measures and midsagittal cerebellar growth across preterm groups.** Extremely preterm (EP, black), very preterm (VP, magenta), and late preterm (LP, light blue) infants are shown throughout. **(A and B)** Association between bedside ultrasound measures and segmented MRI cerebellar volume in infants with ultrasound performed within 2 weeks of MRI ($n = 35$): coronal composite (A) and midsagittal composite (B), with ultrasound composites expressed as *Z*-scores; two-sided Pearson correlation. **(C)** Midsagittal composite cerebellar size (*Z*-score) at 28, 33, and 35 weeks post-menstrual age (PMA); 28 weeks includes EP and VP only. Box plots, median and IQR with individual subjects overlaid; pairwise two-sided Wilcoxon rank-sum tests. **(D)** Midsagittal cerebellar growth rate (*Z*-score per week) by group. Bars, mean; error bars, 95% CI; pairwise slope contrasts from mixed-effects models, Tukey-adjusted. (C and D), $n = 146$ (EP = 80, VP = 45, LP = 21).

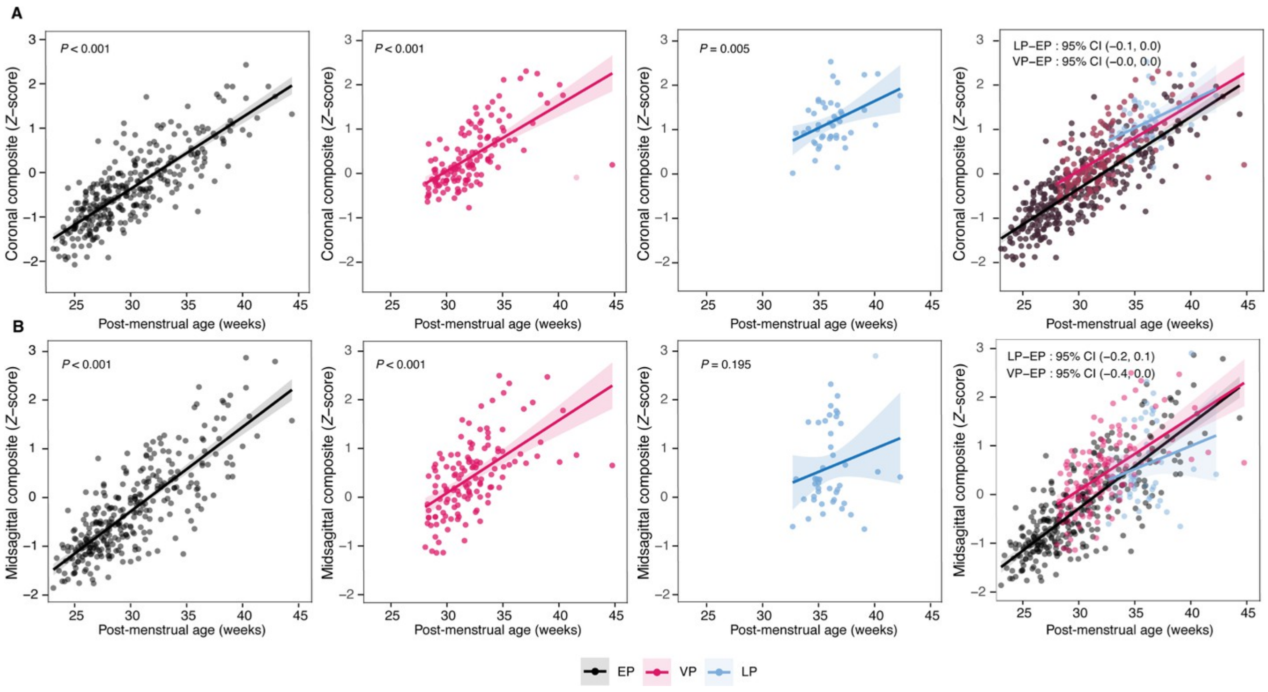


**Fig. S7. Longitudinal cerebellar growth trajectories across gestational age groups. (A and B**) Coronal composite cerebellar size (A) and midsagittal composite cerebellar size (B), each expressed as a *Z*-score and plotted against post-menstrual age (PMA) for extremely preterm (EP, black), very preterm (VP, magenta), and late preterm (LP, light blue) infants ($n$ = 146; EP = 80, VP = 45, LP = 21). The first three columns show EP, VP, and LP trajectories, respectively; the fourth column shows all groups overlaid. Each point represents one ultrasound measurement. Solid lines, linear mixed-effects model fits (PMA × group, per-infant random effect); shaded bands, 95% CI. The overlaid panels report pairwise slope differences (LP−EP and VP−EP) with Tukey-adjusted 95% CI. *P* values in the group-specific panels test the within-group PMA slope. Growth rates did not differ detectably between groups in either plane.

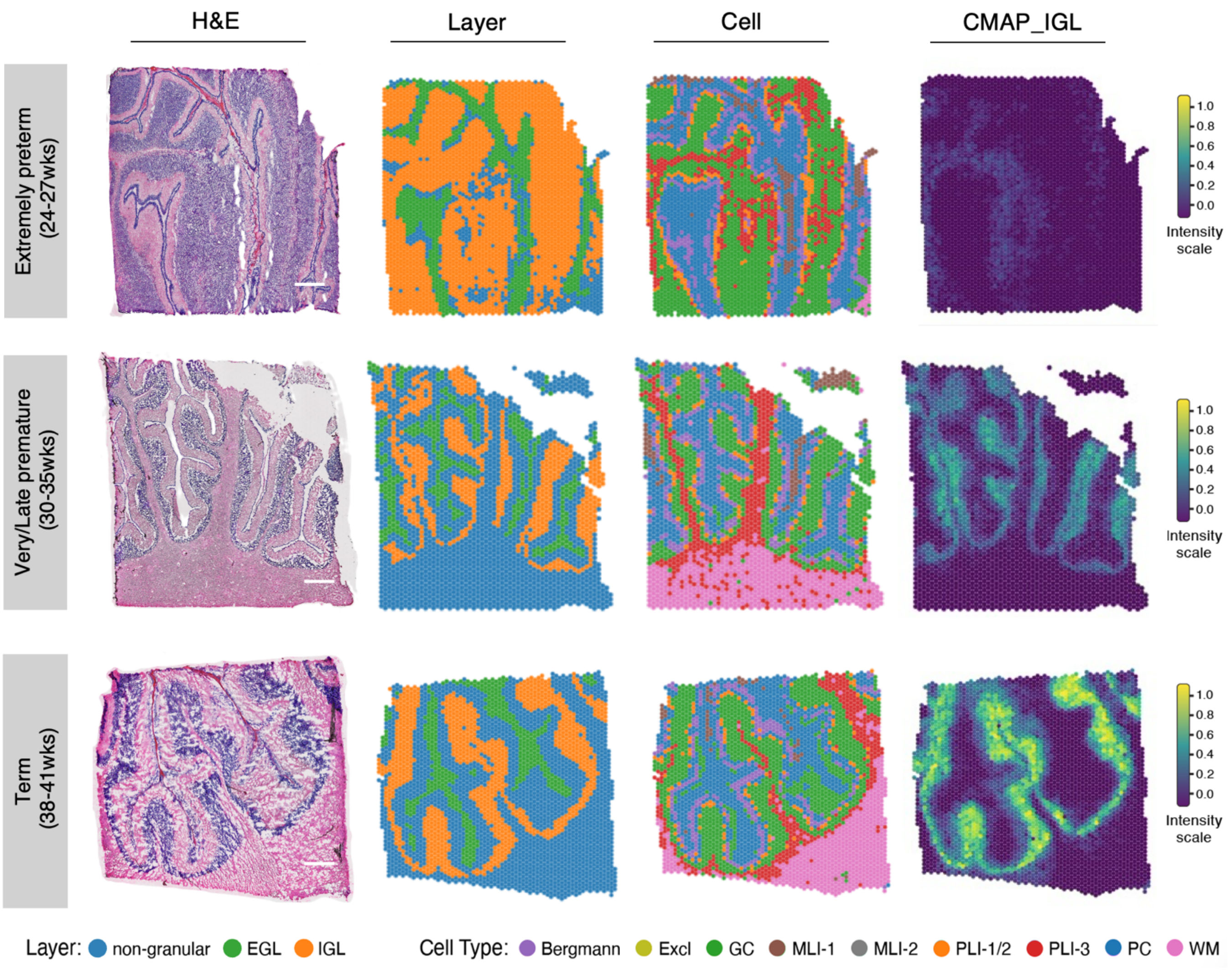


**Fig. S8. Spatial transcriptomic architecture of the human cerebellar cortex across the gestational spectrum.** Representative postmortem cerebellar sections profiled by Visium spatial transcriptomics. Rows (top to bottom): extremely preterm (24–27 weeks gestational age at birth, GA), very/late preterm (30–35 weeks GA), and term (38–40 weeks GA) subjects. Columns (left to right): hematoxylin and eosin (H&E) staining of the profiled section; spatial annotation of cortical layers (Layer); spatial annotation of cell types (Cell); and spatial activity of CMAP_IGL, shown as an example of a cerebellar maturation-associated pattern (CMAP) derived by Bayesian non-negative matrix factorization (CoGAPS). Scale bars, 1 mm. Each spot represents one Visium capture location. Layer annotations: non-granular, external granule layer (EGL), and internal granule layer (IGL). Cell-type annotations: Bergmann glia, excluded tissue (Excl), granule cells (GC), molecular layer interneurons (MLI-1, MLI-2), Purkinje layer interneurons (PLI-1/2, PLI-3), Purkinje cells (PC), and white matter (WM). CMAP activity is shown on a normalized scale from 0.0 to 1.0 (color bar). Analyses were performed on *n* = 11 subjects with two sections per subject (22 sections total; tables S6 and S7).

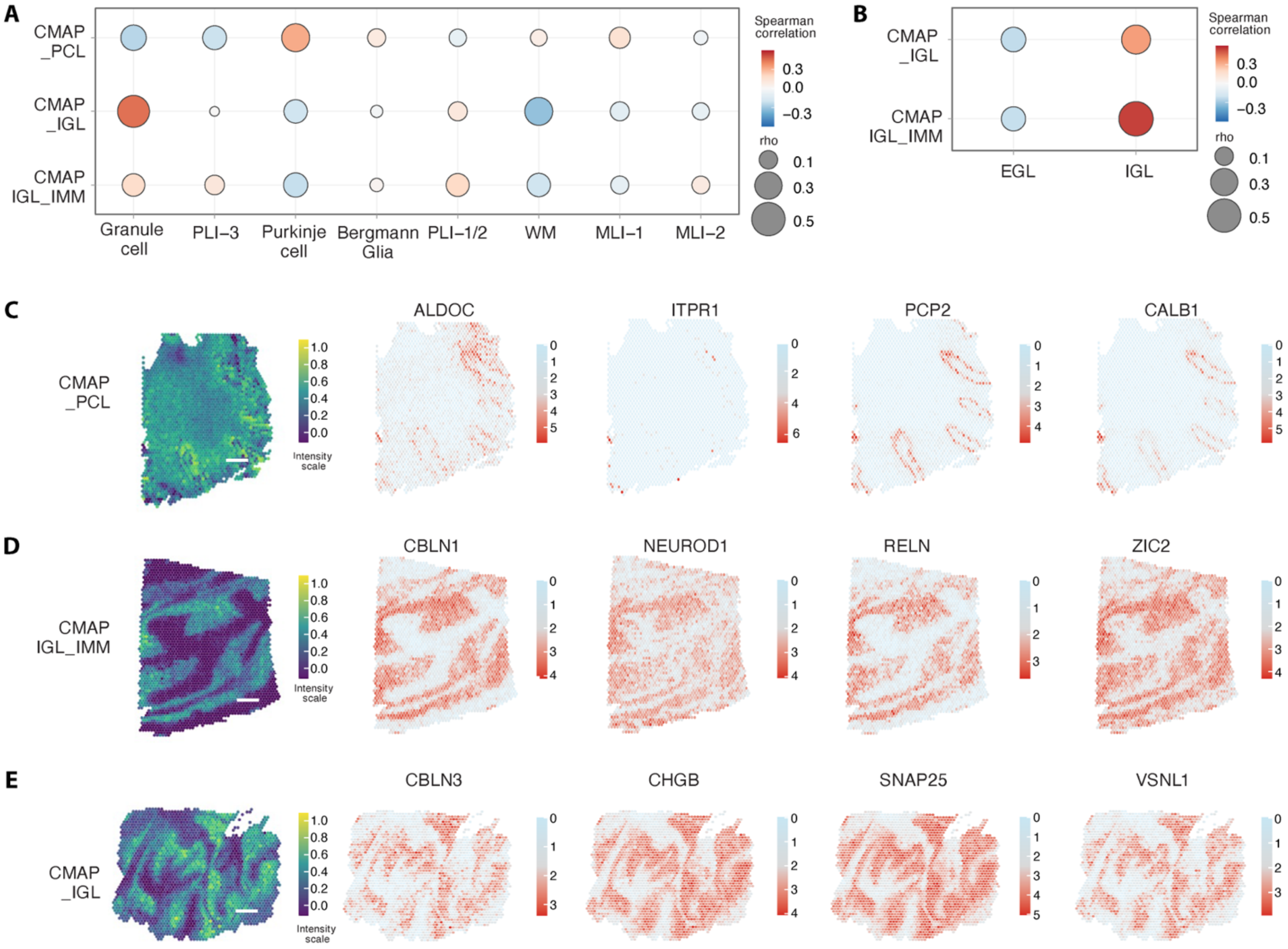


**Fig. S9. Cerebellar maturation-associated patterns localize to distinct cell types and laminar compartments. (A)** Spearman correlations between cerebellar maturation-associated pattern (CMAP) activity and cell-type annotations across the spatial transcriptomic cohort. Dot color, Spearman rho; dot size, absolute correlation magnitude (|rho|). MLI, molecular layer interneuron; PLI, Purkinje layer interneuron; WM, white matter. **(B)** Spearman correlations between CMAP_IGL and CMAP_IGL_IMM activity and laminar annotations of the external granule layer (EGL) and internal granule layer (IGL), displayed as in (A). **(C to E)** Representative spatial activity maps of CMAP_PCL (C), CMAP_IGL_IMM (D), and CMAP_IGL (E), shown alongside spot-level expression of canonical marker genes: *ALDOC*, *ITPR1*, *PCP2*, and *CALB1* (C); *CBLN1*, *NEUROD1*, *RELN*, and *ZIC2* (D); and *CBLN3*, *CHGB*, *SNAP25*, and *VSNL1* (E). Scale bars, 1 mm. Color scales indicate normalized CMAP activity (left) and spot-level gene expression (right). Representative sections are from subjects born at 24.4 (C), 26 (D), and 39 (E) weeks of gestation. (A to E) $n = 11$ subjects, two sections per subject (22 sections total).

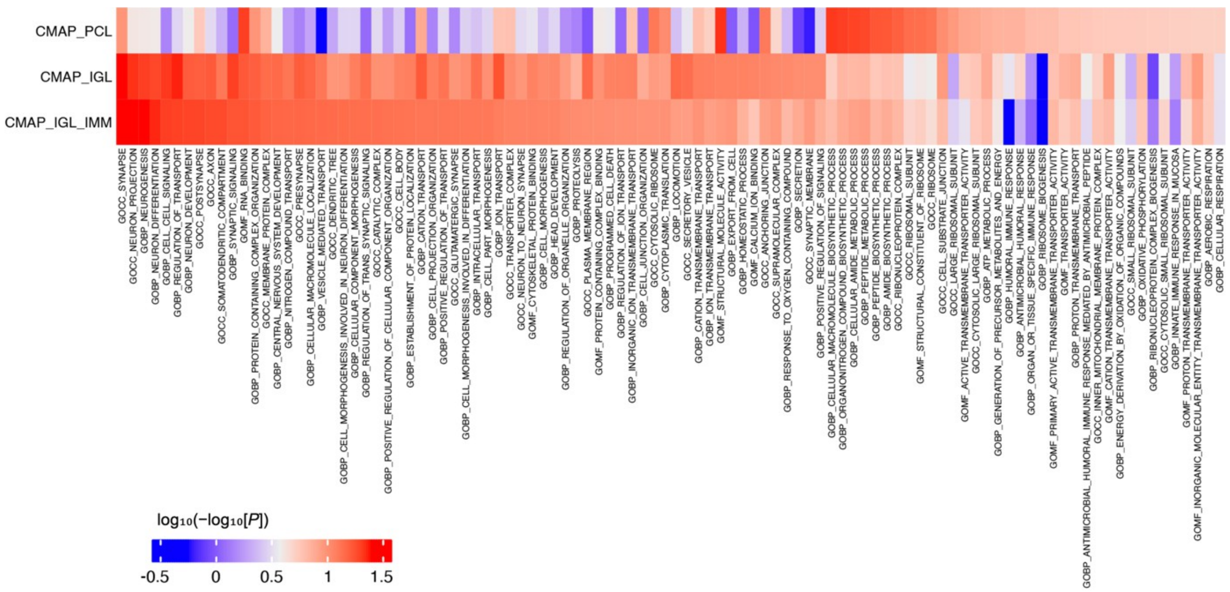


**Fig. S10. Functional enrichment of cerebellar maturation-associated patterns.** Heat map of Gene Ontology (GO) enrichment across gene sets associated with CMAP_PCL, CMAP_IGL, and CMAP_IGL_IMM. Columns represent GO terms enriched in at least one CMAP-associated gene set, spanning biological process (GOBP), cellular component (GOCC), and molecular function (GOMF) categories. Enrichment profiles differed across CMAPs, reflecting distinct developmental, synaptic, metabolic, and cellular programs. Color encodes the transformed nominal *P* value from fast gene set enrichment analysis (fGSEA), $\log_{10}(-\log_{10}[\text{nominal } P])$. CMAPs were derived from 11 subjects with two sections per subject (22 sections total).

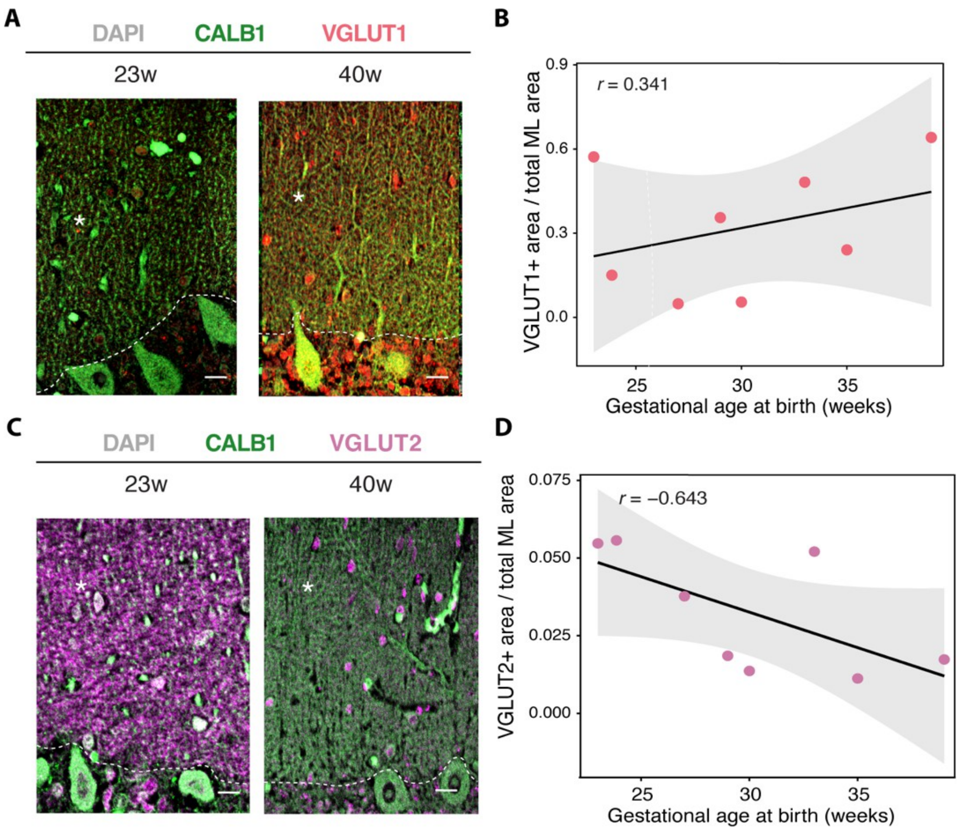


**Fig. S11. VGLUT1 and VGLUT2 immunoreactivity in the cerebellar molecular layer across gestational age at birth. (A)** Representative immunofluorescence images of the cerebellar molecular layer (ML) from subjects born at 23 and 40 weeks' gestation, stained for calbindin (CALB1, green), VGLUT1 (red), and DAPI (gray). Asterisks, molecular layer; dashed lines, Purkinje cell layer–molecular layer border. Scale bar, 10 μm. **(B)** VGLUT1-positive area as a fraction of total ML area plotted against gestational age at birth (Pearson $r = 0.341$). **(C)** Representative immunofluorescence images stained for CALB1 (green), VGLUT2 (magenta), and DAPI (gray), as in (A). Scale bar, 10 μm. **(D)** VGLUT2-positive area as a fraction of total ML area plotted against gestational age at birth (Pearson $r = -0.643$). (B and D) Each point represents one subject and the mean of five regions of interest (ROIs); line, linear fit; shaded band, 95% CI. Eight subjects were analyzed per marker (nine unique subjects; the term subject differed between markers).

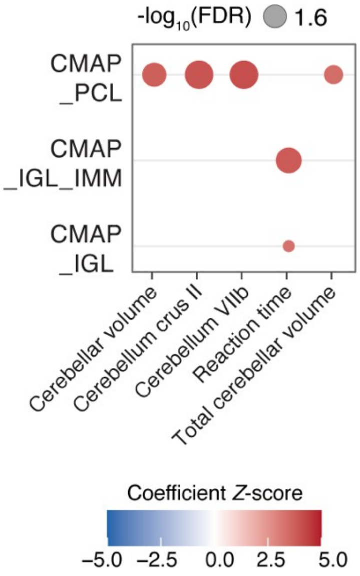


**Fig. S12. LDSC-based partitioned heritability enrichment of CMAP gene sets across neuroimaging and cognitive traits.** Partitioned heritability enrichment of CMAP_PCL, CMAP_IGL_IMM, and CMAP_IGL cerebellar maturation-associated pattern (CMAP) gene sets across cerebellar imaging and cognitive traits was assessed by linkage disequilibrium score regression (LDSC) using published genome-wide association study summary statistics (table S8). Rows indicate CMAP gene sets and columns indicate traits. Only associations passing a false discovery rate (FDR) of <0.05 are shown. Dot size, $-\log_{10}$ FDR; color, regression coefficient *Z*-score. CMAP gene sets were derived from the spatial transcriptomic cohort ($n = 11$ subjects, two sections per subject).

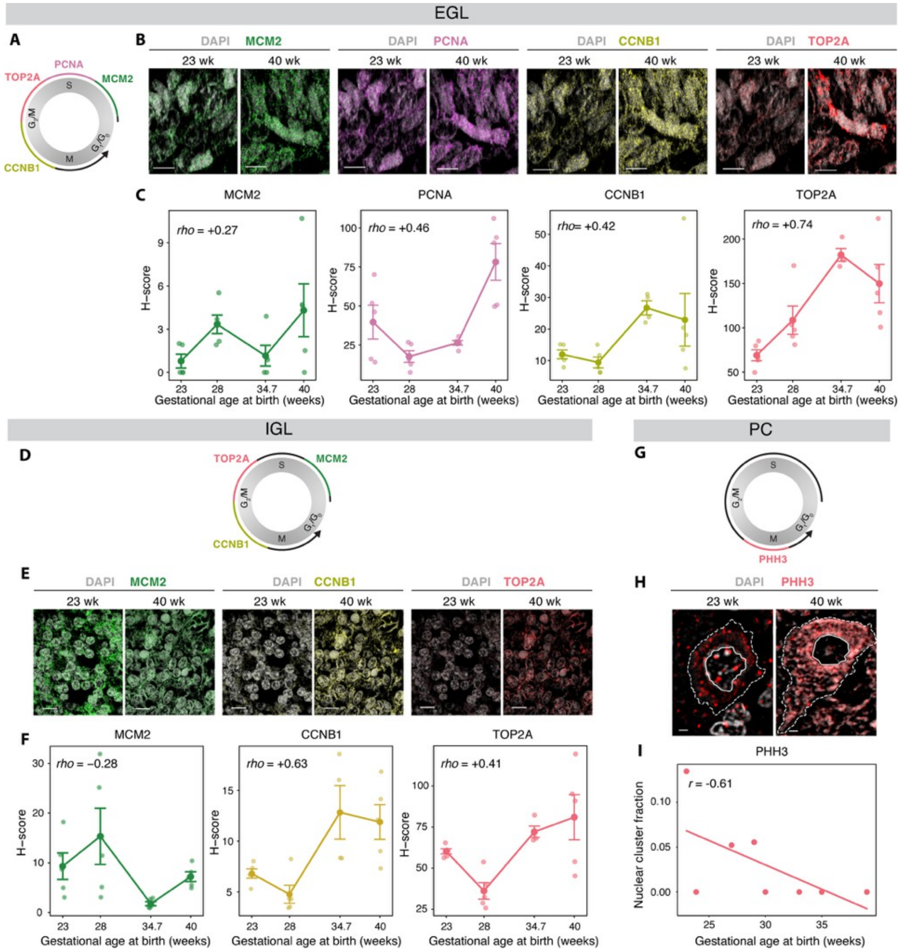


**Fig. S13. Orthogonal validation of phase-associated molecular states across cerebellar lineages and gestational ages.** **(A)** Schematic of *MCM2*, *PCNA*, *CCNB1*, and *TOP2A* transcript positions along the inferred cell cycle continuum in the external granule layer (EGL). **(B)** Representative RNAscope images of the EGL in subjects born at 23 and 40 weeks of gestation, showing *MCM2* (green), *PCNA* (purple), *CCNB1* (yellow), *TOP2A* (magenta), and DAPI (gray). Scale bar, 10 µm. **(C)** H-score quantification of EGL marker expression across gestational age at birth ($n$ = 4 subjects; 4–5 regions of interest [ROIs] per subject). Small points, individual ROIs; large points, subject means; lines connect subject means; error bars, SEM across ROIs. Spearman rho between subject-mean H-score and gestational age at birth is shown in each panel. **(D)** Schematic of *MCM2*, *TOP2A*, and *CCNB1* transcript positions along the inferred cell cycle continuum in the internal granule layer (IGL). **(E)** Representative RNAscope images of the IGL for the markers in (D), with colors and staining as in (B). Scale bar, 10 µm. **(F)** H-score quantification of IGL marker expression across gestational age at birth, as in (C). **(G)** Schematic of phosphorylated histone H3 (PHH3) localization along the phase-associated continuum in Purkinje cells (PCs). **(H)** Representative immunofluorescence images of PCs showing PHH3 (magenta) with DAPI (gray); dashed outlines, annotated cells. Scale bar, 2 µm. **(I)** PHH3 nuclear cluster fraction in PCs plotted against gestational age at birth ($n$ = 8 subjects; 5 ROIs per subject, 5–6 PCs per ROI). The nuclear cluster fraction was calculated as the number of nuclear PHH3 clusters divided by total nuclear PHH3 puncta per cell and averaged per subject. Two-sided Pearson correlation.

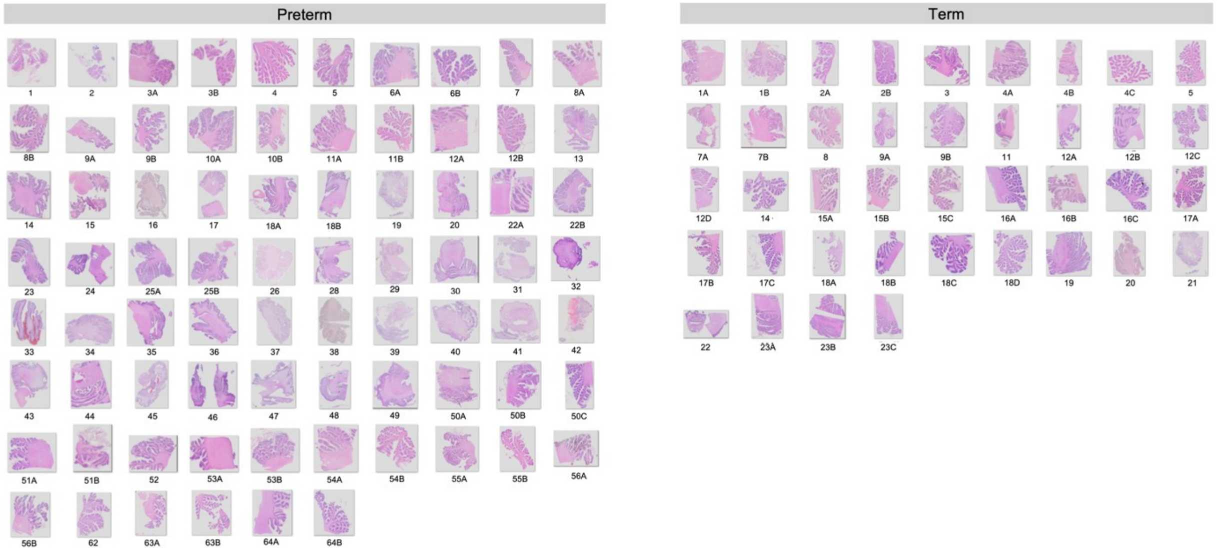


**Fig. S14. Whole-slide hematoxylin and eosin images used for deep learning–based neuropathological analysis.** Whole-slide hematoxylin and eosin (H&E) images of formalin-fixed, paraffin-embedded postmortem cerebellar sections from the deep-learning neuropathology cohort (*n* = 77; preterm, *n* = 57; term, *n* = 20). Five subjects also contributed to the spatial transcriptomic cohort (fig. S1). Each thumbnail represents one cerebellar section. Subject identifiers are numbered within group (tables S11 and S12); suffixes (A, B, C, etc.) denote multiple sections from the same subject. Thumbnails are individually scaled for layout and not shown at a common magnification; quantification was performed on calibrated whole-slide images at native resolution.

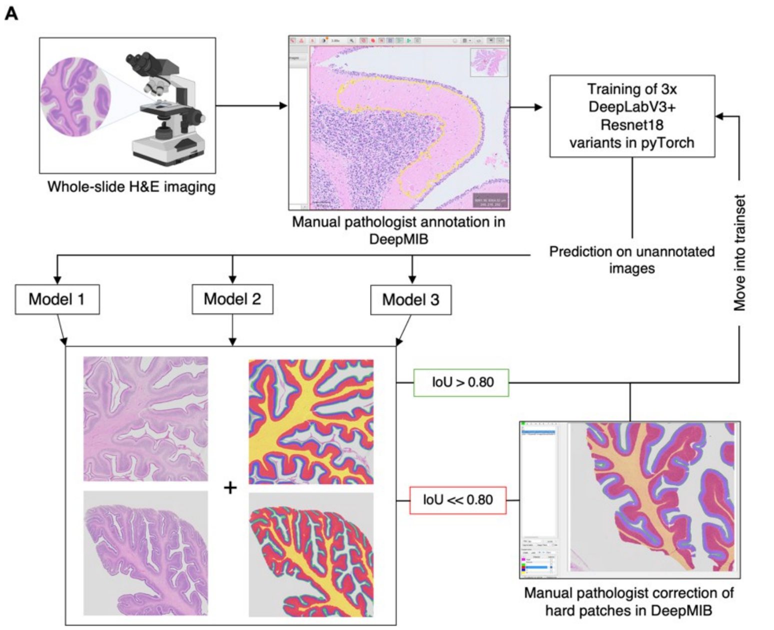


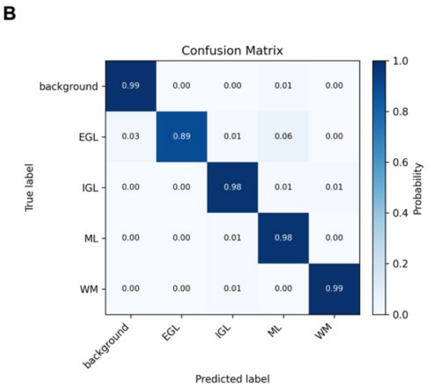


C

| Metric | Median | Mean | Q1–Q3 | SD | n (tiles) |
|---|---|---|---|---|---|
| **IoU** | 0.989 | 0.976 | 0.981–0.992 | 0.096 | 213 |
| **Dice** | 0.994 | 0.983 | 0.990–0.996 | 0.096 | 213 |
| **MSD (px)** | 5.86 | 6.30 | 4.26–7.71 | 3.29 | 211 |
| **Topology** | 0.53 | 0.85 | 0.24–1.00 | 1.08 | 213 |

**Fig. S15. Deep-learning segmentation workflow and performance. (A)** Pathologist-in-the-loop workflow for cerebellar layer segmentation from whole-slide hematoxylin and eosin (H&E) images. Manually annotated regions in DeepMIB were used to train three DeepLabV3+ ResNet18 models. Predictions were applied to previously unannotated images; regions with IoU > 0.80 were incorporated into the training set, whereas lower-IoU patches were manually reviewed and corrected by a pathologist before retraining. This iterative process was repeated until model performance stabilized. **(B)** Class-wise confusion matrix for segmentation of background, external granule layer (EGL), internal granule layer (IGL), molecular layer (ML), and white matter (WM) on a held-out validation set. Purkinje cells were quantified separately. **(C)** Segmentation performance on the held-out validation set, reported as intersection over union (IoU), Dice coefficient, mean symmetric surface distance (MSD; pixels), and topology score. Validation metrics were calculated from $n$ = 213 image tiles (MSD, $n$ = 211). Median IoU and Dice coefficients were 0.989 and 0.994, respectively.

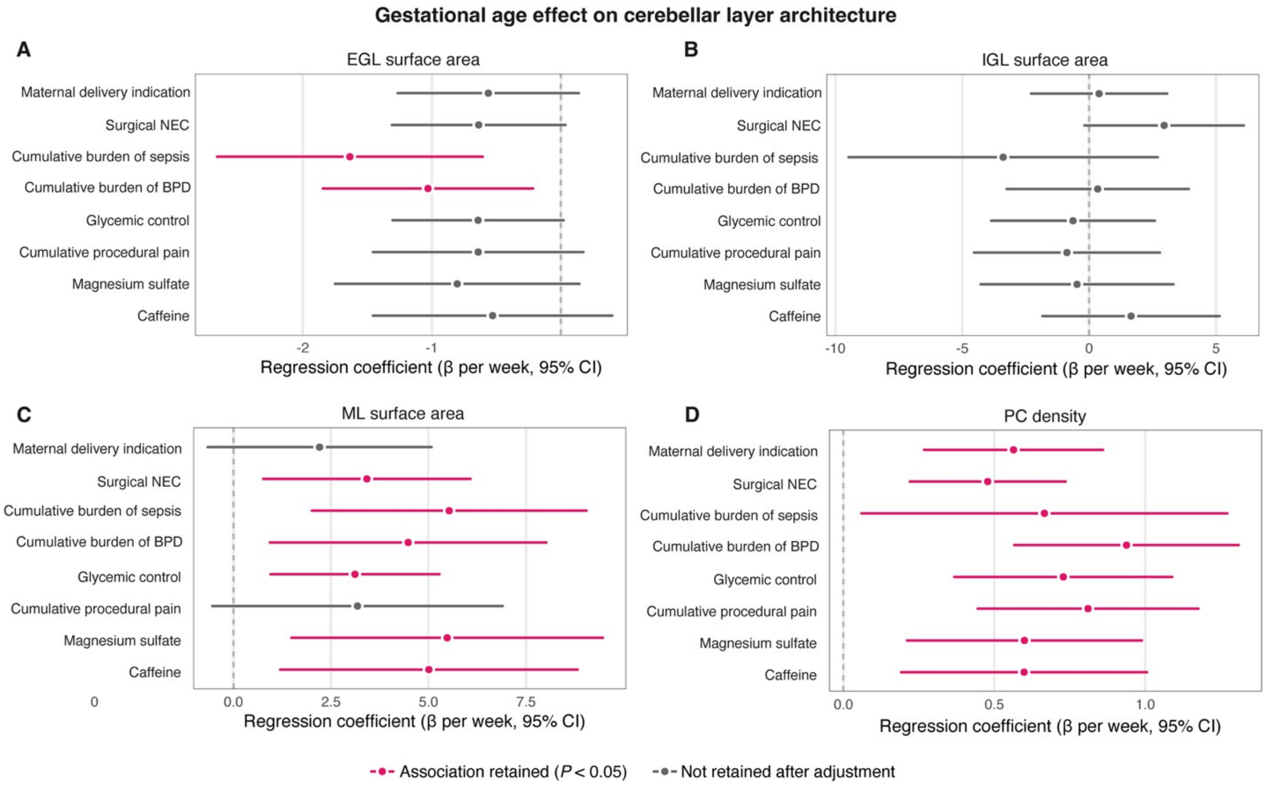


**Fig. S16. Persistence of gestational age associations with cerebellar layer architecture after adjustment for individual perinatal exposures. (A to D)** Associations between gestational age at birth (GA) and cerebellar cytoarchitecture after adjustment for individual perinatal exposures: external granule layer (EGL) surface area (A), internal granule layer (IGL) surface area (B), and molecular layer (ML) surface area (C), each normalized to ML length (mm² per µm ML), and Purkinje cell (PC) density (D; PCs per mm ML), in the deep-learning–based neuropathology cohort. Each row shows the GA regression coefficient (β per week) and 95% CI from a separate model including post-menstrual age at death and the indicated exposure as covariates. Points, β; horizontal lines, 95% CI; dashed line, β = 0. Magenta, association retained after adjustment ($P < 0.05$); gray, not retained. Perinatal exposures included maternal delivery indication, surgical necrotizing enterocolitis (NEC), cumulative burden of sepsis, cumulative burden of bronchopulmonary dysplasia (BPD), glycemic control, cumulative procedural pain, magnesium sulfate, and caffeine. Each exposure was entered separately as an additive covariate; no interaction terms were included. Model size ranged from $n$ = 21 (sepsis) to $n$ = 71 (NEC) and was the same across all four panels for each exposure. Each panel has its own x-scale.

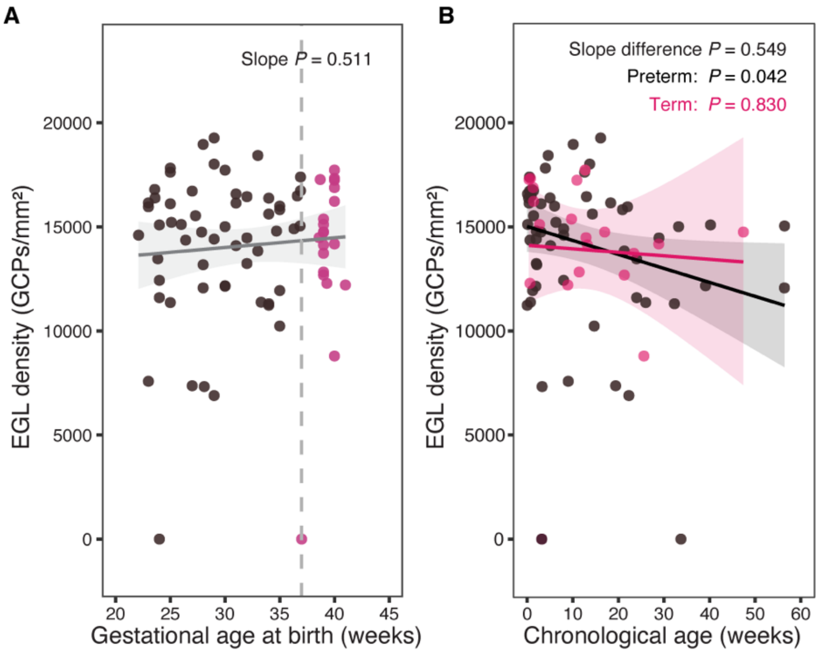


**Fig. S17. External granule layer density across gestational age at birth and chronological age.** External granule layer (EGL) granule cell precursor (GCP) density (GCPs per mm²) in the neuropathology cohort (*n* = 77 subjects; preterm, *n* = 57, black; term, *n* = 20, magenta). Each point represents one subject. **(A)** EGL density plotted against gestational age at birth. Line, ordinary least-squares fit; shaded band, 95% CI; dashed line, 37 weeks (preterm–term boundary). EGL density was not associated with gestational age at birth (slope *P* = 0.511). **(B)** EGL density plotted against chronological age, with group-specific fits (preterm slope, −67.13 [95% CI, −130.31 to −3.95], *P* = 0.042; term slope, −16.80 [95% CI, −168.03 to 134.43], *P* = 0.830). Shaded bands, 95% CI. The preterm and term slopes did not differ (slope-difference *P* = 0.549).

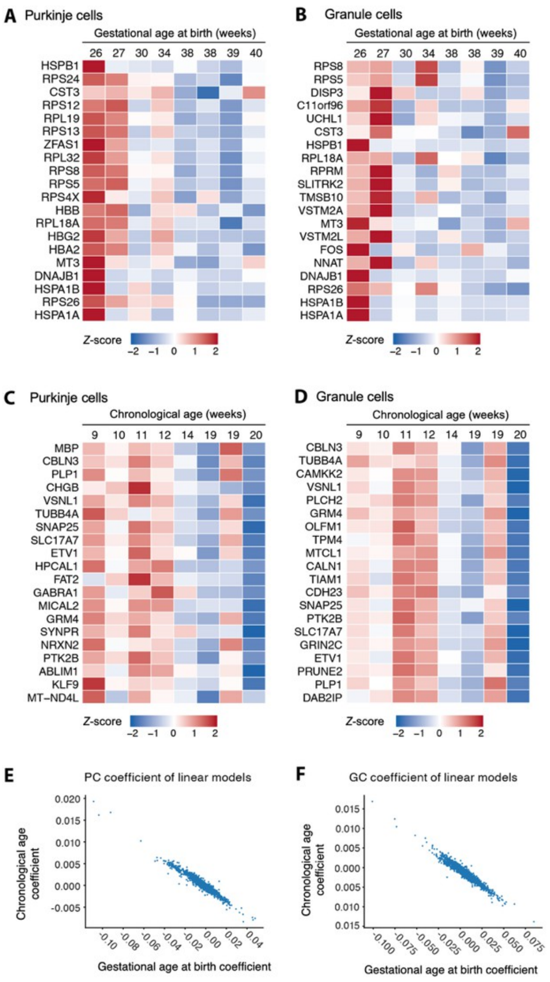


**Fig. S18. Gestational age- and chronological age-associated gene expression patterns are inversely related. (A and B)** Heat maps of genes negatively associated with gestational age at birth (GA) in Purkinje cells (A) and granule cells (B), from spatial transcriptomic samples matched by post-menstrual age at death. Values are *Z*-scored mean expression per subject, ordered by GA. **(C and D)** Genes positively associated with GA (Fig. 4A and E) displayed across chronological age (CA) in Purkinje cells (C) and granule cells (D), illustrating the inverse relationship between GA- and CA-associated expression in this matched cohort. **(E and F)** Gene-level regression coefficients for GA (x axis) versus CA (y axis) in Purkinje cells (E) and granule cells (F). Each point represents one gene. (A to F), *n* = 8 PMA-matched subjects (tables S6 and S7).

**Table S1. Demographic and clinical characteristics of the preterm infant in vivo cohort.**

Values are number (percentage) for categorical variables and mean ± SD (number of infants with available data) for continuous variables; percentages include infants of unknown status.

| **Characteristic** | **Total cohort (*n* = 176)** | **Extremely preterm (EP) <28 weeks (*n* = 98)** | **Very preterm (VP) 28 to <32 weeks (*n* = 54)** | **Late preterm (LP) 32 to <37 weeks (*n* = 24)** |
|---|---|---|---|---|
| **Sex** | | | | |
| Male | 107 (60.8) | 57 (58.2) | 36 (66.7) | 14 (58.3) |
| Female | 69 (39.2) | 41 (41.8) | 18 (33.3) | 10 (41.7) |
| **Race** | | | | |
| American Indian or Alaska Native | 2 (1.1) | 2 (2.0) | 0 (0.0) | 0 (0.0) |
| Asian | 6 (3.4) | 4 (4.1) | 1 (1.9) | 1 (4.2) |
| Black or African American | 94 (53.4) | 54 (55.1) | 29 (53.7) | 11 (45.8) |
| White | 31 (17.6) | 16 (16.3) | 9 (16.7) | 6 (25.0) |
| More than one race | 2 (1.1) | 1 (1.0) | 0 (0.0) | 1 (4.2) |
| Other | 39 (22.2) | 20 (20.4) | 14 (25.9) | 5 (20.8) |
| Unknown | 2 (1.1) | 1 (1.0) | 1 (1.9) | 0 (0.0) |
| **Ethnicity** | | | | |
| Hispanic or Latino | 25 (14.2) | 14 (14.3) | 8 (14.8) | 3 (12.5) |
| Not Hispanic or Latino | 107 (60.8) | 61 (62.2) | 31 (57.4) | 15 (62.5) |
| Unknown | 44 (25.0) | 23 (23.5) | 15 (27.8) | 6 (25.0) |
| **Insurance status** | | | | |
| Public | 86 (48.9) | 48 (49.0) | 30 (55.6) | 8 (33.3) |
| Private | 79 (44.9) | 46 (46.9) | 21 (38.9) | 12 (50.0) |
| None | 8 (4.5) | 3 (3.1) | 1 (1.9) | 4 (16.7) |
| Unknown | 3 (1.7) | 1 (1.0) | 2 (3.7) | 0 (0.0) |
| Maternal age (years) | 30.8 ± 6.3 (172) | 31.5 ± 6.2 (96) | 29.1 ± 6.0 (52) | 31.9 ± 6.7 (24) |
| GA (weeks) | 27.6 ± 3.7 (176) | 25.0 ± 1.4 (98) | 29.2 ± 1.0 (54) | 34.7 ± 1.6 (24) |
| Birth weight (g) | 1138.5 ± 705.1 (175) | 750.7 ± 262.6 (97) | 1229.4 ± 293.6 (54) | 2501.4 ± 826.1 (24) |
| Birth HC (cm) | 25.3 ± 3.8 (67) | 22.8 ± 2.6 (34) | 26.8 ± 1.2 (26) | 32.1 ± 4.0 (7) |

Abbreviations: EP, extremely preterm; GA, gestational age at birth; HC, head circumference; LP, late preterm; SD, standard deviation; VP, very preterm.

**Table S2. Gestational age effect on term-equivalent age cerebellar volume, adjusted for individual perinatal exposures in the in vivo cohort.**

Cerebellar volume was modeled on post-menstrual age at scan and one perinatal exposure at a time; gestational age (GA) was then added and tested by nested F test. Unadjusted GA effect: β = 230.2 $mm^3$ per week (95% CI 50.0 to 410.4; *n* = 120).

| **Perinatal exposure adjusted for** | **GA β ($mm^3$ per week)** | **95% CI** | ***P*** | ***n*** |
|---|---|---|---|---|
| Maternal delivery indication | 216.6 | 33.0 to 400.2 | 0.021 | 115 |
| Surgical necrotizing enterocolitis (NEC) | 202.9 | 26.2 to 379.7 | 0.025 | 120 |
| Cumulative burden of bronchopulmonary dysplasia (BPD) | 167.3 | −14.9 to 349.6 | 0.072 | 117 |
| Cumulative burden of sepsis | 200.3 | 17.9 to 382.7 | 0.032 | 118 |
| Magnesium sulfate exposure | 206.9 | 16.3 to 397.5 | 0.034 | 120 |
| Caffeine exposure | 295.9 | 48.2 to 543.6 | 0.020 | 120 |
| Glycemic control (glucose time in range, 70–180 mg/dL) | 221.3 | 38.5 to 404.0 | 0.018 | 118 |
| Cumulative procedural pain (PIPP) | 378.7 | 36.3 to 721.1 | 0.031 | 116 |

Abbreviations: BPD, bronchopulmonary dysplasia; CI, confidence interval; GA, gestational age at birth; NEC, necrotizing enterocolitis; PIPP, Premature Infant Pain Profile.

**Table S3. Effect of infant sex on the primary in vivo analyses.**

Each primary estimate is given without and with infant sex in the model. β denotes a regression slope; delay odds ratios compare VP with EP and sex odds ratios are male versus female; values in parentheses are 95% CIs, given for odds ratios only. Sex distribution: EP = 41 female / 57 male, VP = 18 female / 36 male, LP = 10 female / 14 male ($n$ = 176); sex was not associated with gestational age group ($\chi^2$ $P$ = 0.569) or with gestational age (female 27.5 ± 3.8 versus male 27.7 ± 3.6 weeks, $P$ = 0.766).

| Analysis | *n* | Primary estimate (without sex → with sex) | Sex effect (male vs. female) | Sex *P* | Interaction *P* |
|---|---|---|---|---|---|
| Cerebellar volume (mm³) | 120 | β GA 230.2 → 231.2 | +148.4 mm³ | 0.813 | 0.501 (GA × sex) |
| Cerebellar volume / ICV ($\times 10^{-4}$) | 120 | β GA 1.27 → 1.19 | −11.8 | 0.273 | 0.581 (GA × sex) |
| US coronal composite (*Z* score) | 504 scans / 146 infants | β PMA 0.157 → 0.159 | +0.255 *Z* | <0.001 | 0.301 (PMA × sex) |
| Any delay | 145 | OR (VP vs. EP) 0.21 → 0.21 | OR 1.21 (0.51 to 2.83) | 0.657 | Not fitted |
| Motor delay | 143 | OR (VP vs. EP) 0.29 → 0.29 | OR 0.85 (0.40 to 1.77) | 0.659 | Not fitted |
| Cognitive delay | 140 | OR (VP vs. EP) 0.46 → 0.46 | OR 1.09 (0.54 to 2.18) | 0.812 | Not fitted |
| Language delay | 143 | OR (VP vs. EP) 0.44 → 0.42 | OR 1.49 (0.74 to 3.02) | 0.266 | Not fitted |
| Delay at equal cerebellar volume | 98 (EP 62 / VP 36) | OR 0.88 per 1,000 mm³ (0.76 to 1.02) | OR 2.53 (0.92 to 7.38) | 0.078 | 0.512 (volume × sex) |

Abbreviations: CI, confidence interval; EP, extremely preterm; GA, gestational age at birth; ICV, intracranial volume; LP, late preterm; OR, odds ratio; PMA, post-menstrual age; US, cranial ultrasound; VP, very preterm.

**Table S4. Gestational age and cerebellar volume as predictors of neurodevelopmental delay, adjusted for individual perinatal exposure in vivo cohort.**
Logistic regression of motor, cognitive, and language delay, with one perinatal exposure added per model, restricted to extremely and very preterm infants. Gestational age (GA) → delay: likelihood-ratio $\chi^2$ *P* for the GA group term (*n* = 140-143). Cerebellar volume → delay: odds ratio (OR) per 1,000 $mm^3$ with 95% confidence interval (CI) and Wald *P* for term-equivalent age cerebellar volume (*n* = 90–97).

| Exposure adjusted for | Delay domain | GA → delay, *P* | Cerebellar volume → delay, OR (95% CI) | Wald *P* |
|---|---|---|---|---|
| Delivery indication (spontaneous vs maternal) | Motor | 0.001 | 0.81 (0.71, 0.92) | 0.002 |
| | Cognitive | 0.092 | 0.87 (0.77, 0.98) | 0.026 |
| | Language | 0.014 | 0.90 (0.80, 1.01) | 0.081 |
| Surgical necrotizing enterocolitis | Motor | <0.001 | 0.80 (0.70, 0.91) | <0.001 |
| | Cognitive | 0.025 | 0.86 (0.76, 0.97) | 0.014 |
| | Language | 0.021 | 0.90 (0.80, 1.00) | 0.061 |
| Sepsis burden | Motor | 0.001 | 0.79 (0.69, 0.91) | <0.001 |
| | Cognitive | 0.038 | 0.86 (0.76, 0.97) | 0.014 |
| | Language | 0.027 | 0.89 (0.79, 1.00) | 0.043 |
| Bronchopulmonary dysplasia burden | Motor | <0.001 | 0.79 (0.68, 0.91) | <0.001 |
| | Cognitive | 0.032 | 0.86 (0.76, 0.97) | 0.013 |
| | Language | 0.033 | 0.90 (0.80, 1.02) | 0.099 |
| Glycemic control (glucose time-in-range) | Motor | 0.001 | 0.79 (0.69, 0.91) | <0.001 |
| | Cognitive | 0.058 | 0.86 (0.76, 0.97) | 0.012 |
| | Language | 0.040 | 0.90 (0.80, 1.01) | 0.075 |
| Cumulative procedural pain (PIPP) | Motor | 0.265 | 0.60 (0.42, 0.87) | 0.006 |
| | Cognitive | 0.207 | 0.79 (0.64, 0.99) | 0.042 |
| | Language | 0.329 | 0.83 (0.68, 1.01) | 0.064 |
| Magnesium sulfate | Motor | <0.001 | 0.79 (0.69, 0.90) | <0.001 |
| | Cognitive | 0.064 | 0.85 (0.76, 0.96) | 0.011 |
| | Language | 0.017 | 0.90 (0.80, 1.01) | 0.066 |
| Caffeine | Motor | 0.001 | 0.80 (0.70, 0.92) | 0.001 |
| | Cognitive | 0.046 | 0.86 (0.76, 0.97) | 0.017 |
| | Language | 0.017 | 0.89 (0.80, 1.01) | 0.064 |

Abbreviations: CI, confidence interval; GA, gestational age; OR, odds ratio.

**Table S5. Differences in coronal cerebellar growth rate between gestational age groups, adjusted for individual perinatal exposures.**

Gestational age (GA) group, post-menstrual age (PMA), and the exposure were entered in a base model. For each perinatal exposure, this was compared by likelihood-ratio test with a model additionally including the GA group × PMA interaction.

| Perinatal exposure adjusted for | $\chi^2$ | *P* | Scans (infants) |
|---|---|---|---|
| Maternal delivery indication | 3.86 | 0.145 | 483 (139) |
| Surgical necrotizing enterocolitis (NEC) | 2.94 | 0.230 | 504 (146) |
| Cumulative burden of bronchopulmonary dysplasia (BPD) | 3.52 | 0.172 | 495 (143) |
| Cumulative burden of sepsis | 2.89 | 0.236 | 493 (143) |
| Magnesium sulfate exposure | 3.03 | 0.219 | 504 (146) |
| Caffeine exposure | 3.12 | 0.211 | 504 (146) |
| Glycemic control (glucose time in range, 70–180 mg/dL) | 3.02 | 0.221 | 491 (143) |
| Cumulative procedural pain (PIPP) | 4.30 | 0.117 | 473 (139) |

Abbreviations: BPD, bronchopulmonary dysplasia; GA, gestational age at birth; NEC, necrotizing enterocolitis; PIPP, Premature Infant Pain Profile; PMA, post-menstrual age.

**Table S6. Demographic and clinical characteristics of the postmortem spatial transcriptomic cohort, by preterm and term group.**

Values are number (percentage) for categorical variables and mean ± SD for continuous variables.

| Characteristic | Preterm ($n$ = 7) | Term ($n$ = 4) |
|---|---|---|
| **Sex** | | |
| Male | 3 (42.9) | 3 (75.0) |
| Female | 4 (57.1) | 1 (25.0) |
| **Race** | | |
| Black or African American | 5 (71.4) | 1 (25.0) |
| White | 2 (28.6) | 3 (75.0) |
| **Ethnicity** | | |
| Hispanic or Latino | 0 (0.0) | 0 (0.0) |
| Not Hispanic or Latino | 6 (85.7) | 2 (50.0) |
| Unknown | 1 (14.3) | 2 (50.0) |
| GA (weeks) | 30.1 ± 4.6 | 38.8 ± 1.0 |
| PMA at death (weeks) | 42.7 ± 5.7 | 49.6 ± 1.4 |
| Brain weight (g) | 368.0 ± 133.5 | 576.0 |

Abbreviations: GA, gestational age at birth; PMA, post-menstrual age; SD, standard deviation.

**Table S7. Demographic and clinical characteristics of the postmortem spatial transcriptomic cohort, by individual subject.**

Data for the 7 preterm and 4 term subjects analyzed by spatial transcriptomics. Conditions lists the conditions documented at or contributing to death.

| Subject ID | Sex | GA (weeks) | PMA at death (weeks) | Brain weight (g) | Race | Ethnicity | Conditions |
|---|---|---|---|---|---|---|---|
| Preterm 1 | M | 27.0 | 46.3 | Not recorded | W | NH | SIDS |
| Preterm 2 | F | 35.0 | 36.2 | 288 | B | NH | HLHS |
| Preterm 52 | F | 35.0 | 41.1 | 512 | B | NH | Asphyxia, co-sleeping |
| Preterm 57 | F | 34.0 | 45.2 | 517 | B | NH | SIDS |
| Preterm 58 | M | 30.0 | 49.4 | 420 | W | Unknown | SIDS |
| Preterm 59 | F | 26.0 | 46.2 | 267 | B | NH | Multiorgan failure |
| Preterm 60 | M | 24.0 | 34.3 | 204 | B | NH | NEC |
| Term 1 | M | 40.0 | 48.6 | Not recorded | W | NH | SIDS |
| Term 3 | M | 39.0 | 48.5 | 576 | B | NH | Asphyxia |
| Term 24 | M | 38.0 | 51.5 | Not recorded | W | Unknown | Pneumonia |
| Term 25 | F | 38.0 | 50.0 | Not recorded | W | Unknown | SIDS |

Abbreviations: B, Black or African American; F, female; GA, gestational age at birth; HLHS, hypoplastic left heart syndrome; M, male; NEC, necrotizing enterocolitis; NH, not Hispanic or Latino; PMA, post-menstrual age; SIDS, sudden infant death syndrome; W, White.

**Table S8. Datasets used in the partitioned heritability analysis.**

Genome-wide association study (GWAS) reference datasets used for partitioned heritability analysis by linkage disequilibrium score regression. Full citations appear in the main reference list under the indicated reference numbers.

| Ref. | Study | Trait or resource |
|---|---|---|
| 54 | Identification of common genetic risk variants for autism spectrum disorder | ASD |
| 55 | Genome-wide analyses of ADHD identify 27 risk loci, refine the genetic architecture and implicate several cognitive domains | ADHD |
| 56 | Genetic common variants associated with cerebellar volume and their overlap with mental disorders | Cerebellar volume |
| 57 | Genome-wide association study of cerebellar volume: heritable mechanisms underlying brain development and mental health | Cerebellar volume |
| 58 | An expanded set of genome-wide association studies of brain imaging phenotypes in UK Biobank | UK Biobank brain IDPs |
| 59 | Insights into the genetic architecture of cerebellar lobules derived from the UK Biobank | Cerebellar lobule volumes |
| 60 | Genome-wide association meta-analysis in 269,867 individuals: genetic and functional links to intelligence | Intelligence |
| 61 | Study of 300,486 individuals identifies 148 independent genetic loci influencing general cognitive function | General cognitive function and reaction time |
| 62 | Gene discovery and polygenic prediction from a GWAS of educational attainment in 1.1 million individuals | Educational attainment |
| 63 | Examining the role of common variants in rare neurodevelopmental conditions | Rare neurodevelopmental conditions |
| 64 | Genome-wide association meta-analysis of age at onset of walking in over 70,000 infants of European ancestry | Age at onset of walking |

Abbreviations: ADHD, attention deficit hyperactivity disorder; ASD, autism spectrum disorder; IDP, imaging-derived phenotype; LD, linkage disequilibrium; UK, United Kingdom.

**Table S9. Correlation of CMAP activity with gestational age at birth and post-menstrual age at death within clinical subgroups.**

Spearman rank-correlation coefficients (rho) between subject-level CMAP activity and gestational age at birth or post-menstrual age at death. The first row is the full cohort; each subsequent row is restricted to the subjects indicated. Coefficients are descriptive; no *P* values are reported.

| **Subjects included** | ***n*** | **CMAP_IGL_IMM** | | **CMAP_IGL** | | **CMAP_PCL** | |
|---|---|---|---|---|---|---|---|
| | | **GA** | **PMA** | **GA** | **PMA** | **GA** | **PMA** |
| All subjects | 11 | −0.320 | −0.155 | 0.616 | 0.591 | −0.566 | 0.027 |
| Preterm | 7 | 0.018 | 0.143 | 0.468 | 0.429 | −0.811 | −0.036 |
| Male | 6 | −0.200 | 0.029 | 0.771 | 0.771 | −0.714 | −0.486 |
| Female | 5 | −0.667 | −0.300 | 0.103 | 0.500 | −0.564 | 0.500 |
| Without NEC | 10 | −0.524 | −0.261 | 0.488 | 0.467 | −0.335 | 0.358 |
| Without sepsis | 10 | −0.189 | −0.127 | 0.555 | 0.612 | −0.372 | 0.091 |
| Without chorioamnionitis or infection | 9 | −0.201 | −0.100 | 0.594 | 0.467 | −0.444 | −0.083 |
| Without preeclampsia | 9 | 0.017 | −0.067 | 0.445 | 0.617 | −0.261 | 0.150 |
| Without ventilatory support | 9 | −0.361 | −0.217 | 0.387 | 0.467 | −0.134 | 0.500 |
| With spontaneous onset of labor | 9 | 0.017 | −0.067 | 0.445 | 0.617 | −0.261 | 0.150 |

Abbreviations: CMAP, cerebellar maturation-associated pattern; GA, gestational age at birth; NEC, necrotizing enterocolitis; PMA, post-menstrual age at death.

**Table S10. Demographic and clinical characteristics of the postmortem neuropathology cohort and its analytic subsets, by preterm and term group.**

Values are number (percentage) for categorical variables and mean ± SD (number of subjects with available data) for continuous variables. The final rows give the number of subjects analyzed by each method.

| Characteristic | Preterm ($n$ = 57) | Term ($n$ = 20) |
|---|---|---|
| **Sex** | | |
| Male | 29 (50.9) | 12 (60.0) |
| Female | 28 (49.1) | 8 (40.0) |
| **Race** | | |
| Black or African American | 26 (45.6) | 5 (25.0) |
| White | 9 (15.8) | 7 (35.0) |
| Other | 2 (3.5) | 0 (0.0) |
| Unknown | 20 (35.1) | 8 (40.0) |
| **Ethnicity** | | |
| Hispanic or Latino | 3 (5.3) | 2 (10.0) |
| Not Hispanic or Latino | 35 (61.4) | 12 (60.0) |
| Unknown | 19 (33.3) | 6 (30.0) |
| GA (weeks) | 29.4 ± 4.5 | 39.3 ± 0.8 |
| PMA at death (weeks) | 42.0 ± 14.3 | 51.9 ± 12.1 |
| Chronological age (weeks) | 12.6 ± 14.0 | 12.7 ± 12.1 |
| Brain weight (g) | 361.4 ± 223.0 (53) | 558.8 ± 254.4 (14) |
| **Analytic subsets** | | |
| Deep learning–based neuropathology | 57 (100.0) | 20 (100.0) |
| RNAscope | 3 (5.3) | 1 (5.0) |
| Immunofluorescence | 7 (12.3) | 2 (10.0) |

GA, gestational age at birth; PMA, post-menstrual age; SD, standard deviation.

Because of their length, **tables S11** and **S12** are provided as separate Excel files rather than within this document.

**Table S13. Effect of sex on the postmortem neuropathology analyses.**

Each gestational age (GA) coefficient is given without and with infant sex in the model, alongside *P* values for the sex effect and for the two interactions. All models include all 77 subjects. Sex was not associated with preterm/term status (Fisher exact $P = 0.604$) or with GA (female 31.0 ± 6.0 versus male 32.8 ± 5.6 weeks, $P = 0.175$).

| **Measure** | **GA *β* per week, without sex → with sex** | **Sex *P*** | **GA × sex *P*** | **Sex × preterm/term *P*** |
|---|---|---|---|---|
| EGL area ($mm^2$ per µm ML) | −0.726 → −0.739 | 0.790 | 0.623 | 0.509 |
| IGL area ($mm^2$ per µm ML) | 2.70 → 2.92 | 0.318 | 0.794 | 0.426 |
| ML area ($mm^2$ per µm ML) | 3.40 → 3.29 | 0.606 | 0.748 | 0.386 |
| Purkinje cell density (PCs per mm ML) | 0.403 → 0.406 | 0.846 | 0.211 | 0.631 |
| EGL density (cells per $mm^2$) | 46.6 → 41.2 | 0.633 | 0.131 | 0.209 |

Abbreviations: EGL, external granule layer; GA, gestational age at birth; IGL, internal granule layer; ML, molecular layer; PC, Purkinje cell; SD, standard deviation.

**Table S14. Gestational age effect on cerebellar layer architecture after adjustment for individual clinical factors in the postmortem neuropathology cohort.**

For each clinical factor, each layer measure was modeled on that factor and gestational age (GA); the GA term was tested by nested F test and only the GA coefficient is shown. Glycemic control is glucose time in range (70–180 mg/dL) and cumulative procedural pain is the summed Premature Infant Pain Profile (PIPP) score.

| Clinical factor adjusted for | Layer | GA β per week | 95% CI | *P* | *n* |
|---|---|---|---|---|---|
| Maternal delivery indication | EGL area | −0.56 | −1.26 to 0.14 | 0.113 | 54 |
| | IGL area | 0.39 | −2.30 to 3.07 | 0.773 | |
| | ML area | 2.21 | −0.66 to 5.08 | 0.129 | |
| | PC density | 0.56 | 0.27 to 0.86 | <0.001 | |
| Surgical NEC | EGL area | −0.64 | −1.31 to 0.03 | 0.063 | 71 |
| | IGL area | 2.95 | −0.19 to 6.09 | 0.065 | |
| | ML area | 3.42 | 0.76 to 6.08 | 0.012 | |
| | PC density | 0.48 | 0.22 to 0.74 | <0.001 | |
| Cumulative burden of sepsis | EGL area | −1.64 | −2.66 to −0.61 | 0.004 | 21 |
| | IGL area | −3.39 | −9.48 to 2.70 | 0.257 | |
| | ML area | 5.53 | 2.01 to 9.05 | 0.004 | |
| | PC density | 0.67 | 0.06 to 1.27 | 0.033 | |
| Cumulative burden of BPD | EGL area | −1.03 | −1.84 to −0.22 | 0.014 | 45 |
| | IGL area | 0.33 | −3.25 to 3.92 | 0.852 | |
| | ML area | 4.48 | 0.94 to 8.02 | 0.014 | |
| | PC density | 0.94 | 0.57 to 1.31 | <0.001 | |
| Glycemic control | EGL area | −0.64 | −1.30 to 0.02 | 0.057 | 40 |
| | IGL area | −0.64 | −3.86 to 2.59 | 0.691 | |
| | ML area | 3.11 | 0.95 to 5.28 | 0.006 | |
| | PC density | 0.73 | 0.37 to 1.09 | <0.001 | |
| Cumulative procedural pain | EGL area | −0.64 | −1.46 to 0.17 | 0.119 | 39 |
| | IGL area | −0.88 | −4.54 to 2.79 | 0.630 | |
| | ML area | 3.18 | −0.54 to 6.90 | 0.092 | |
| | PC density | 0.81 | 0.44 to 1.18 | <0.001 | |
| Magnesium sulfate | EGL area | −0.80 | −1.75 to 0.14 | 0.093 | 39 |
| | IGL area | −0.48 | −4.28 to 3.32 | 0.799 | |
| | ML area | 5.48 | 1.49 to 9.47 | 0.008 | |
| | PC density | 0.60 | 0.21 to 0.99 | 0.003 | |
| Caffeine | EGL area | −0.53 | −1.46 to 0.40 | 0.256 | 49 |
| | IGL area | 1.65 | −1.84 to 5.14 | 0.346 | |
| | ML area | 5.01 | 1.20 to 8.82 | 0.011 | |
| | PC density | 0.60 | 0.19 to 1.01 | 0.005 | |

Abbreviations: BPD, bronchopulmonary dysplasia; CI, confidence interval; EGL, external granule layer; GA, gestational age at birth; IGL, internal granule layer; ML, molecular layer; NEC, necrotizing enterocolitis; PC, Purkinje cell; PIPP, Premature Infant Pain Profile.

**Table S15. Demographic and clinical characteristics of the postmortem subjects analyzed by RNAscope in situ hybridization.**

The four subjects, three preterm and one term, drawn from the postmortem neuropathology cohort. Conditions lists the conditions documented at or contributing to death.

| Subject ID | Sex | GA (weeks) | PMA at death (weeks) | Race | Ethnicity | Brain weight (g) | Conditions |
|---|---|---|---|---|---|---|---|
| Preterm 14 | M | 23.0 | 41.3 | B | NH | 368 | Respiratory failure; ARDS |
| Preterm 23 | F | 23.9 | 47.9 | Unknown | Unknown | 347.9 | NEC |
| Preterm 29 | M | 28.0 | 30.1 | B | NH | 228 | NEC; maternal eclampsia |
| Term 22 | M | 40.0 | 41.4 | Unknown | Unknown | 376.5 | Multiorgan failure |

Abbreviations: ARDS, acute respiratory distress syndrome; B, Black or African American; GA, gestational age at birth; NEC, necrotizing enterocolitis; NH, not Hispanic or Latino; PMA, post-menstrual age.

**Table S16. Demographic and clinical characteristics of the immunofluorescence subjects in the postmortem neuropathology cohort.**

Data for each of the 9 immunofluorescence subjects. Conditions lists the conditions documented at or contributing to death. All seven preterm subjects were stained for both VGLUT1 and VGLUT2; the term subjects differ by marker (Term 8 for VGLUT1, Term 2 for VGLUT2), so each marker was assessed in eight subjects and nine subjects contributed in total.

| Subject ID | Sex | GA (weeks) | PMA at death (weeks) | Race | Ethnicity | Brain weight (g) | Conditions | Markers |
|---|---|---|---|---|---|---|---|---|
| Preterm 1 | M | 27.0 | 46.3 | W | NH | Not recorded | SIDS | VGLUT1, VGLUT2 |
| Preterm 5 | F | 23.0 | 45.0 | B | NH | 275 | Respiratory failure | VGLUT1, VGLUT2 |
| Preterm 6 | F | 30.0 | 42.7 | W | NH | 380 | SIDS | VGLUT1, VGLUT2 |
| Preterm 10 | M | 29.0 | 42.7 | W | NH | 470 | SIDS | VGLUT1, VGLUT2 |
| Preterm 11 | M | 33.0 | 54.3 | B | NH | 720 | SIDS | VGLUT1, VGLUT2 |
| Preterm 12 | M | 35.0 | 49.7 | B | NH | 650 | SIDS | VGLUT1, VGLUT2 |
| Preterm 23 | F | 23.9 | 47.9 | Unknown | Unknown | 347.9 | NEC | VGLUT1, VGLUT2 |
| Term 2 | F | 39.0 | 60.3 | B | NH | 920 | Pseudomonas septicemia | VGLUT2 |
| Term 8 | F | 39.0 | 52.2 | Unknown | HL | 600 | Unknown | VGLUT1 |

Abbreviations: B, Black or African American; GA, gestational age at birth; HL, Hispanic or Latino; NEC, necrotizing enterocolitis; NH, not Hispanic or Latino; PMA, post-menstrual age; SIDS, sudden infant death syndrome; VGLUT1, vesicular glutamate transporter 1; VGLUT2, vesicular glutamate transporter 2; W, White.

**Table S17. RNAscope probe panels.**

All probes are RNAscope 2.5 LS Probe, Homo sapiens (Advanced Cell Diagnostics).

| Panel | Transcript class | Gene | Catalog no. | Channel |
|---|---|---|---|---|
| 1 | Purkinje cell | PLP1 | 564578 | C3 |
| 1 | Purkinje cell | KLF9 | 582558 | C5 |
| 1 | Granule cell | TUBB4A | 1695038 | C2 |
| 1 | Granule cell | GRM4 | 1256538 | C4 |
| 2 | Cell cycle | MCM2 | 451468 | C1 |
| 2 | Cell cycle | PCNA | 553078 | C2 |
| 2 | Cell cycle | CCNB1 | 465378 | C3 |
| 2 | Cell cycle | TOP2A | 470328 | C4 |

Abbreviations: CCNB1, cyclin B1; GRM4, glutamate metabotropic receptor 4; KLF9, Kruppel-like factor 9; MCM2, minichromosome maintenance complex component 2; PCNA, proliferating cell nuclear antigen; PLP1, proteolipid protein 1; TOP2A, DNA topoisomerase II alpha; TUBB4A, tubulin beta 4A class IVa.

**Table S18. Opal fluorophore and channel assignments for the RNAscope probe panels.**

All reagents are Opal Reagent Packs (Akoya Biosciences).

| Fluorophore | Catalog no. | Working dilution | Channel |
|---|---|---|---|
| Opal 480 | FP1500001KT | 1:750 | C1 |
| Opal 520 | FP1487001KT | 1:750 | C2 |
| Opal 570 | FP1488001KT | 1:1500 | C3 |
| Opal 620 | FP1495001KT | 1:1500 | C4 |
| Opal 690 | FP1497001KT | 1:1500 | C5 |

**Table S19. Antibodies used for immunofluorescent analyses.**

| Antibody | Host | Detection | Dilution | Catalog no. | Lot no. | Supplier | Use |
|---|---|---|---|---|---|---|---|
| VGLUT1 | Guinea pig, polyclonal | — | 1:200 | 135 304 | 3-46 | Synaptic Systems | Primary |
| VGLUT2 | Rabbit, polyclonal | — | 1:200 | 135 403 | Not recorded | Synaptic Systems | Primary |
| Calbindin D-28k | Mouse, monoclonal IgG1 | — | 1:200 | CB300 | 07(F) | SWANT | Primary |
| Phospho-histone H3 (Ser10) | Rabbit, monoclonal | — | 1:200 | 701258 | 2489176 | ThermoFisher Scientific | Primary |
| Cyclin B1 | Mouse, monoclonal | — | 1:200 | sc-245 | K1725 | Santa Cruz Biotechnology | Primary |
| Anti-rabbit IgG (H+L) | Donkey | Alexa Fluor 647 | 1:250 | 711-605-152 | 163459 | Jackson ImmunoResearch | Secondary |
| Anti-mouse IgG (H+L) | Donkey | Alexa Fluor 488 | 1:250 | 715-545-151 | 164101 | Jackson ImmunoResearch | Secondary |
| Anti-guinea pig IgG (H+L) | Donkey | Alexa Fluor 647 | 1:250 | 706-605-148 | 149524 | Jackson ImmunoResearch | Secondary |

Abbreviations: IgG, immunoglobulin G; VGLUT1, vesicular glutamate transporter 1; VGLUT2, vesicular glutamate transporter 2.

**Table S20. Intra-subject correlation between transcriptomic and neuropathological cerebellar metrics.**

Spearman rank correlation coefficients (rho) between per-subject cerebellar maturation-associated pattern (CMAP) activity and normalized cerebellar layer surface areas from deep learning–based neuropathology, in the five postmortem subjects analyzed by both modalities (Preterm 1, 2 and 52; Term 1 and 3). Per-subject CMAP activity is the mean over layer-matched spatial spots: two-sided exact-permutation *P* values (all 120 permutations) are given for reference.

| Comparison | Spearman rho (leave-one-out range) | Exact-permutation *P* |
|---|---|---|
| CMAP_IGL vs. IGL area | −0.70 (−1.00 to −0.40) | 0.233 |
| CMAP_IGL_IMM vs. IGL area | 0.60 (0.20 to 0.80) | 0.350 |
| CMAP_PCL vs. ML area | 0.30 (−0.40 to 0.80) | 0.683 |

Abbreviations: CMAP, cerebellar maturation-associated pattern; IGL, internal granule layer; ML, molecular layer.
